# High-throughput heterospheroid-based screening identifies drugs that reprogram tumor-associated macrophages

**DOI:** 10.1101/2025.11.04.686451

**Authors:** Hiroyuki Tsuchiya, Takehiko Hanaki, Mayu Obora, Jun Yoshida, Mikiya Kishino, Yoshiyuki Fujiwara, Daisuke Nanba

**Affiliations:** Division of Regenerative Medicine and Therapeutics, Department of Genomic Medicine and Regenerative Therapy, Faculty of Medicine, Tottori University, 86 Nishi-cho, Yonago, Tottori 683-8503, Japan; Division of Gastrointestinal and Pediatric Surgery, Department of Surgery, Faculty of Medicine, Tottori University, 86 Nishi-cho, Yonago, Tottori 683-8503, Japan; Division of Medical Education, Department of Medical Education, Faculty of Medicine, Tottori University, 86 Nishi-cho, Yonago, Tottori 683-8503, Japan

**Keywords:** tumor-associated macrophage, macrophage reprogramming, high-content imaging analysis, heterospheroid, screening

## Abstract

The tumor microenvironment (TME) provides a niche for immune evasion and immunotherapy resistance, in part, by recruiting pro-tumor M2-like macrophages. In the present study, using heterospheroids consisting of cancer cells and macrophages, we identified TAM activators, which are compounds that reprogram M2-like tumor-associated macrophages (TAMs) toward the antitumor M1-like phenotype. THP-1- or human peripheral monocyte-derived macrophages were co-cultured with liver cancer cells in an ultra-low attachment dish to generate heterospheroids. Cell surface marker expression and macrophage infiltration into the heterospheroids were assessed by flow cytometry and fluorescence microscopy, respectively. Lipopolysaccharide (LPS) and interferon-γ (IFNγ)-induced M1 marker expression was observed on the macrophages in the homospheroids; however, this induction was suppressed in heterospheroids. Microscopic imaging revealed that macrophage infiltration into the heterospheroids was decreased in the presence of LPS and IFNγ, which prompted us to develop a high-content imaging screen. We identified two compounds [alprostadil (prostaglandin E1) and HX531] with TAM-activating activity. RNA-seq analysis revealed that HX531 modulated the immune and IFN response in cancer cells and cell division in macrophages. Moreover, alprostadil promoted the M1-like polarization of TAMs, increased tumor-infiltrating CD8+ T cells, and enhanced anti-PD-1 antibody therapeutic efficacy in a syngeneic mouse xenograft model. In conclusion, the heterospheroid culture recapitulates the immunosuppressive TME, which prevents the M1 polarization of TAMs. It provides a new platform for screening TAM activators and will enable the development of novel cancer immunotherapeutics when combined with high-content imaging analysis.

## Introduction

Immune checkpoint inhibitors (ICIs) have had a significant impact on cancer therapy; however, a significant proportion of patients do not benefit from ICIs targeting exhausted T-cell immunity [1]. Therefore, drugs with different mechanisms of action are needed for those who relapse or are refractory to ICI therapy.

The tumor microenvironment (TME) contains various types of noncancerous stromal cells, including tumor-associated macrophages (TAMs). The cancer cells influence the behavior of these cells through humoral factors and cell-to-cell interactions to support tumor growth, progression, and immune evasion [2,3]; thus, tumor-associated stromal cells represent an alternative target for cancer immunotherapy. Macrophages phagocytose apoptotic cells and, under certain circumstances, live, non-apoptotic cells, such as transformed or infected cells. They produce inflammatory cytokines that promote T-cell effector function and tumor immunity [3,4]. However, they exhibit plasticity as they polarize between pro-inflammatory/anti-tumor (M1) and anti-inflammatory/pro-tumor (M2) phenotypes [3,4]. In most cases, TAMs are inclined to adopt the M2-like phenotype and exhibit pro-tumor functions, including the promotion of T cell exhaustion and regulatory T cell differentiation [5,6], pro-tumor and immunosuppressive cytokine secretion [7], impaired antigen presentation [8], and the induction of chemoresistance, angiogenesis, and metastasis [9–11]. Thus, M2-like TAMs are associated with poor prognosis in patients with cancer [8].

Several mechanisms underlying TAM polarization have been proposed [12–14], and drugs targeting these pathways are capable of reprogramming M2-like TAMs into the M1-like phenotype [14–18]. These agents (hereafter referred to as TAM activators) are expected to suppress tumor growth and enhance the therapeutic effects of ICIs; however, to establish macrophage-targeted therapy [3], a high-throughput and efficient screening system for TAM activators is required.

Previous studies have indicated that heterospheroids consisting of cancer cells and macrophages have several advantages for the *in vitro* study of TAMs and the TME [19–21], suggesting that a heterospheroid culture is an ideal platform for screening TAM activators. However, flow cytometry and immunostaining of cell surface markers, which are widely used to verify macrophage polarity, compromise throughput performance because of their more complex procedures. In contrast, a previous study established an efficient screening system using bone marrow-derived macrophages (BMDMs) derived from interleukin 1β (Il1β) reporter mice, in which the *luciferase* gene was expressed under the control of the *Il1β* promoter [16]. Carfilzomib, a proteasome inhibitor, was identified as a TAM activator using this screening system. However, this system requires a specific reporter mouse and, more importantly, does not consider the immunosuppressive effects of the TME.

In this study, we demonstrated that heterospheroids consisting of liver cancer (LC) cells and macrophages can reproduce the immunosuppressive milieu that inhibits polarization of macrophages to the M1 phenotype. Moreover, we found that the frequency of macrophages infiltrating into a heterospheroid was reduced in the presence of interferon-γ (IFNγ) and lipopolysaccharide (LPS) (M1 inducers). Based on this result, we constructed a high-content imaging screening system and used an FDA-approved drug library. We identified alprostadil [prostaglandin E1 (PGE1)] and HX531 as TAM activators. RNA sequencing (RNA-seq) analysis of the HX531-treated heterospheroids provided mechanistic insight into the molecular targets of HX531 in the reprogramming of macrophages. The therapeutic effects of alprostadil were confirmed *in vivo*. These results indicate that heterospheroid culture in combination with high-content imaging analysis provides an efficient platform for screening TAM activators and studying the mechanisms underlying TAM polarization in the TME.

## Material and Methods

### Materials

The key research resources Research Resource Identifiers (RRIDs) are listed in Table S1.

### Cell lines

The human male LC cell lines, HLF and HuH6, were purchased from the Japanese Collection of Research Bioresources Cell Bank in August 2018. The human male acute monocytic leukemia cell line THP-1 and the mouse female hepatoblastoma cell line Hepa1-6 were obtained from the RIKEN Bioresource Center in April 2021 and June 2019, respectively. These cell lines have not been previously reported as misidentified or contaminated. LC and Hepa1-6 cells were cultured in DMEM (Shimadzu Diagnostics, Tokyo, Japan) containing 10% FBS (Sigma-Aldrich, St. Louis, MO, USA). The THP-1 cells were cultured in RPMI1640 (Shimadzu Diagnostics) containing 10% FBS. Mycoplasma negativity was assessed by PCR using the VenorGeM Classic Mycoplasma Detection Kit (Minerva BioLabs GmbH, Berlin, Germany). The cell lines were authenticated by STR analysis [Fasmac (Kanagawa, Japan), for human cell lines; American Type Culture Collection (Manassas, VA, USA), for Hepa1-6 cell line]. The LC cells were stably transfected with pAcGFP1-C1 (TaKaRa Bio, Kusatsu, Japan) followed by G418 (Nacalai, Kyoto, Japan) selection. The THP-1 cells were infected with lentivirus expressing hKO1. AcGFP- and hKO1-positive clones were isolated by limited dilution.

### THP-1-derived macrophage preparation

hKO1-expressing THP-1 cells (3 × 10^6^ cells) were seeded into a 10-cm dish and incubated for 72 h in the presence of 200 nM phorbol 12-myristate 13-acetate (PMA; Nacalai). The cells were cultured in PMA-free RPMI1640 containing 10% FBS. The medium was replaced with fresh medium every day for three days to obtain mature macrophages.

### Preparation of the primary macrophages

The isolation of peripheral blood monocytes from a healthy male volunteer was done based on the Declarations of Helsinki and Istanbul and approved by the Ethical Committee of Tottori University (22B016). Written informed consent was obtained from the donor before blood collection. Peripheral blood (60 mL) containing EDTA-2Na was centrifuged at 800 × *g* for 15 min. After discarding the plasma, the buffy coat was removed, diluted more than twice with RPMI1640 (FBS-free), and layered onto Monocytes Spin Medium (pluriSelect, Leipzig, Germany). After centrifugation at 800 × *g* for 20 min, the cells in the middle layer were removed and washed twice with RPMI1640. The cells were then seeded into a 6-well plate and incubated at 37°C in a 5% CO_2_ atmosphere for 1.5 h. The unattached cells were removed by washing three times with PBS, and the attached cells were incubated in RPMI1640 medium supplemented with 10% FBS, 4.5g/L glucose, and 25 ng/mL M-CSF (PeproTech, Rocky Hill, NJ, USA). Half of the medium was replaced with fresh medium every three days for 10 days to obtain mature primary macrophages. The cells were labeled using CellTracker CM-DiI Dye (Thermo Fisher Scientific, Waltham, MA, USA) before spheroid culture.

### Heterospheroid culture for screening and imaging

The cells were detached from the culture dishes with Accutase (Nacalai) for LC cells and THP-1-derived macrophages or trypsin/EDTA (Nacalai) for primary macrophages. For the CD24 blocking experiments, LC cells were incubated with 4 µg/mL of IgG1κ or anti-CD24 antibody (both from Thermo Fisher Scientific), whereas the macrophages were incubated with Clear Back (MBL, Nagoya, Japan) for 30 min on ice. LC cells and macrophages were thoroughly mixed in 0.52% methyl cellulose-containing Ham’s F-12 supplemented with bFGF and EGF (20 ng/mL, from PeproTech), and seeded into an ultra-low attachment 96-well plate (Corning, Kennebunk, ME, USA). The reagents were then added to the cells. The cytokine (IFNγ, IL4, and IL13) and LPS (Sigma-Aldrich) concentrations were 20 and 10 ng/mL, respectively. For screening, drugs in an FDA-approved drug library (1,136 compounds) (Table S1) were added at a final concentration of 1 µM. The library was kindly provided by the Center for Supporting Drug Discovery and Life Science Research, Osaka University, Osaka, Japan. The effects of alprostadil (Selleck, Houston, TX, USA) were determined at a final concentration of 2 µM. The cells were incubated for six days, and imaging analysis was performed by fluorescence microscopy (BZ-X800, KEYENCE, Osaka, Japan).

### Heterospheroid culture for flow cytometry analysis

LC cells and macrophages were mixed and incubated in a 3.5-cm ultralow attachment dish (Sumitomo Bakelite, Tokyo, Japan) for 3 days in the presence of the compounds (Table S1). After dissociation of the heterospheroids with Accutase, the cells were incubated with 4 ng/mL of antibodies in 1% bovine serum albumin/PBS for 30 min on ice. After filtering through a 30-µm mesh membrane (CS CRIE, Kyoto, Japan), the cells were analyzed using LSRFortessa X-20 (BD Biosciences, San Jose, CA, USA) or CytoFLEX S (Beckman Coulter, Brea, CA, USA) flow cytometers. The antibodies used in the study are listed in Table S1.

### RNA-sequencing (RNA-seq)

Heterospheroids were dissociated with Accutase following incubation with DMSO or 1 µM HX531 for two days. HLF cells and macrophages were sorted using FACSAria Fusion (BD Biosciences). Total RNA was isolated using Sepasol-RNA I Super G (Nacalai). Poly(A) mRNA purification and library preparation were performed using the NEBNext Poly(A) mRNA Magnetic Isolation Module and NEBNext Ultra II Directional RNA Library Prep Kit for Illumina (New England BioLabs, Ipswich, MA, USA) based on the manufacturer’s instructions. RNA-seq was performed on NovaSeq 6000 (Illumina, San Diego, CA, USA) with 45–64 million paired-end reads of 50 bases, which were trimmed and mapped to the GRCh39/hg19 human reference genome using the CLC Genomics Workbench (QIAGEN, Hilden, Germany). The counts for the mapped reads were normalized by transcripts per million. The RNA-seq data were submitted to the DDBJ Sequence Read Archive under accession number PRJDB35411.

### Reverse transcription-quantitative polymerase chain reaction (RT-qPCR)

Complementary DNA from the total mRNA was synthesized using ReverTra Ace (Toyobo, Osaka, Japan) and random nanomer oligo DNA. The cDNAs were subjected to qPCR using THUNDERBIRD SYBR qPCR Mix (Toyobo) and ViiA 7 (Thermo Fisher Scientific). The primers were synthesized by Fasmac and are listed in Table S1.

### Animal study

The animal experiments were conducted in compliance with the ARRIVE guidelines and were approved by the Tottori University Animal Use Committee (23-Y-48). The sample size was determined based on the distribution. No exclusion or inclusion criteria were applied, all animals were used in the experiments, and the researchers were aware of the group allocation throughout the experiments. Six-week-old male C57BL/6J mice were purchased from CLEA Japan (Tokyo, Japan). The mice were housed under pathogen-free conditions in a temperature-controlled room illuminated for 12 h daily and allowed free access to food (CE-2; CLEA Japan) and water. Animals received humane care based on the study guidelines established by the Tottori University Subcommittee on Laboratory Animal Care. The mice (3-6 mice/cage) were acclimatized for one week and subcutaneously injected with Hepa1-6 cells. Tumor size was measured with a caliper, and the volume was calculated using the following formula: V = a × b^2^/2, where a and b are the tumor length and width, respectively. Ten days after transplantation, the mice were randomized by tumor size to ensure almost equal mean sizes. To assess the effects of alprostadil, the mice were intraperitoneally injected with 6 ng/g body weight of alprostadil or an equivalent amount of DMSO in corn oil every day for 14 days (n = 5, each). The experiments were performed independently twice. For combination studies, anti-Pd-1 antibody or IgG2aκ (both, Selleck) in combination with alprostadil or DMSO was administered (100 µg per mouse) intraperitoneally once every three days for a total of two or five doses (n = 4-6). The mice were anesthetized with isoflurane and sacrificed by cervical dislocation. To assess the tumor-infiltrated immune cells, tumor tissues were removed, weighed, and dissociated in Accumax (Nacalai) with shaking at 37°C for 30 min. After filtering through a 300-µm mesh membrane (CS CRIE), the cells were subjected to flow cytometry with the antibodies listed in Table S1.

### Statistical analysis

Data were analyzed using SPSS software (ver. 28.0, IBM, Armonk, NY, USA), and graphs were generated using Excel software (ver. 16.87, Microsoft, Redmond, WA, USA). All experimental values are expressed as the mean ± SD. Three or more independent samples were analyzed for each experiment as indicated in the figure legends. Differences between two groups were determined using a Student’s t-test, whereas multiple comparisons were made using Dunnett’s or Tukey’s tests. The cumulative tumor-free rate was analyzed using the log-rank test. *P* < 0.05 was considered statistically significant.

## Results

### The suppression of M1 polarization in heterospheroids

A monocytic leukemia cell line, THP-1, and a liver cancer (LC) cell line, HLF, were used. The cells were stably transfected with hKO1 and AcGFP, respectively, for imaging spheroids. THP-1-derived macrophages (hKO1^+^) were cultured alone to form homospheroids or with HLF cells (AcGFP^+^) to form heterospheroids in ultra-low attachment dishes (Figure 1A). Treatment with M1 inducers significantly induced the expression of CD11B, CD80, CD86, and HLA-DR in the homospheroids (Figure 1B). Although upregulation was also observed in heterospheroids, it was significantly lower compared with that in homospheroids (Figure 1B), suggesting that M1 polarization was suppressed by LC cells. CD14 expression, a mature monocyte marker, was upregulated in heterospheroids and further increased by M1 inducers.

**Figure 1.**
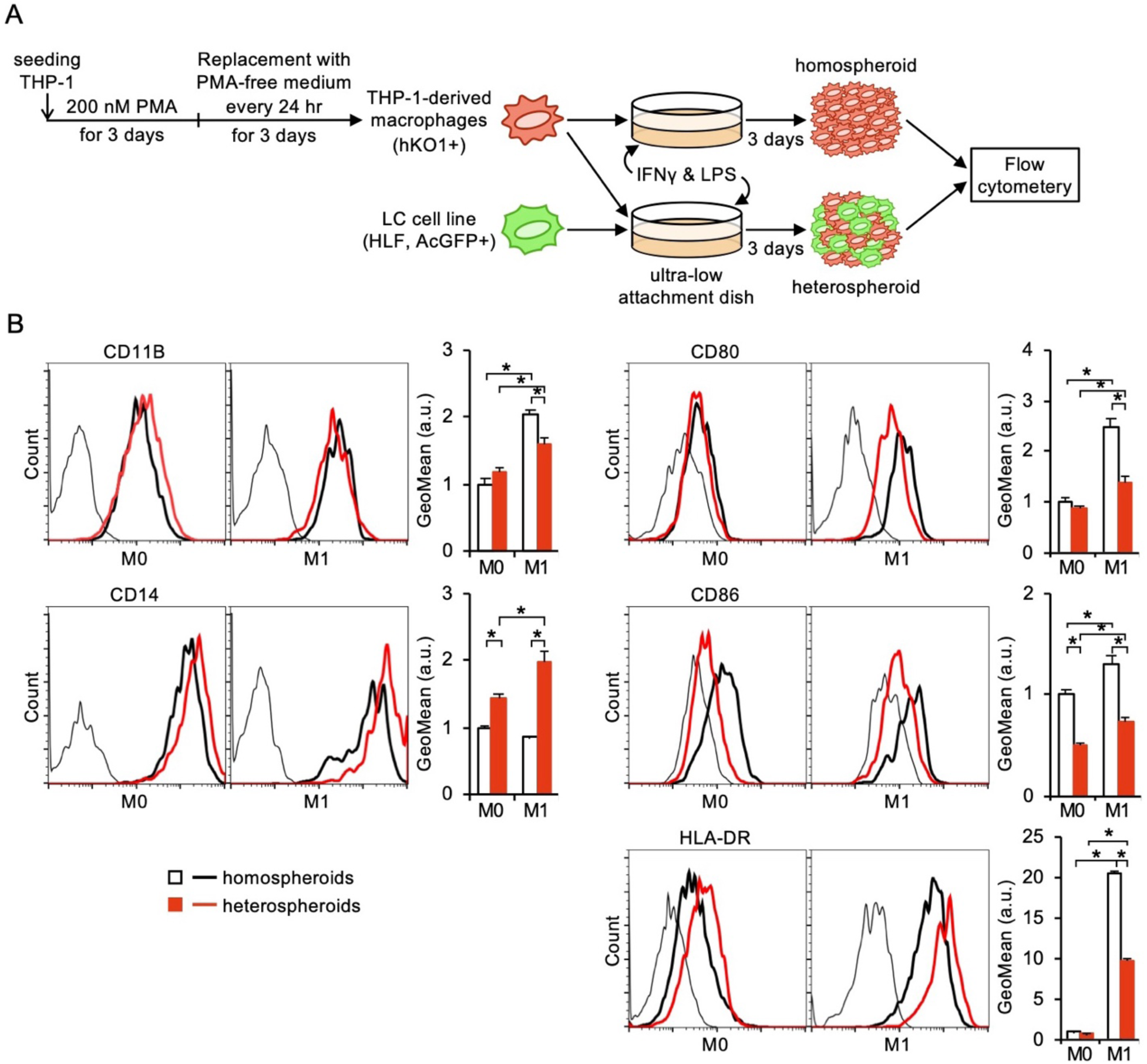
Suppression of M1 polarization in heterospheroids. (A) Preparation of homospheroids consisting of macrophages, and heterospheroids comprised of macrophages and liver cancer (LC) cells. (B) Macrophage marker expression on the macrophage surface. The spheroids were incubated in the absence (M0) or presence (M1) of IFNγ and LPS for 3 days. *, *P* < 0.05, Tukey’s test (*n* = 3).

### Decreased macrophage infiltration in the presence of IFNγ and LPS

Fluorescent microscopy revealed that the number of heterospheroids was increased by M1 inducers compared with untreated controls (M0), whereas a significant decrease was observed in the area of the heterospheroids, the occupancy of macrophages in the heterospheroids (area ratio of hKO1 to AcGFP and hKO1), and the mean fluorescent intensity (MFI) of macrophages in a heterospheroid (hKO1 intensity per heterospheroid area) (Figure 2A). In contrast, IL4 and IL13 (M2 inducers) slightly increased the occupancy and MFI.

**Figure 2.**
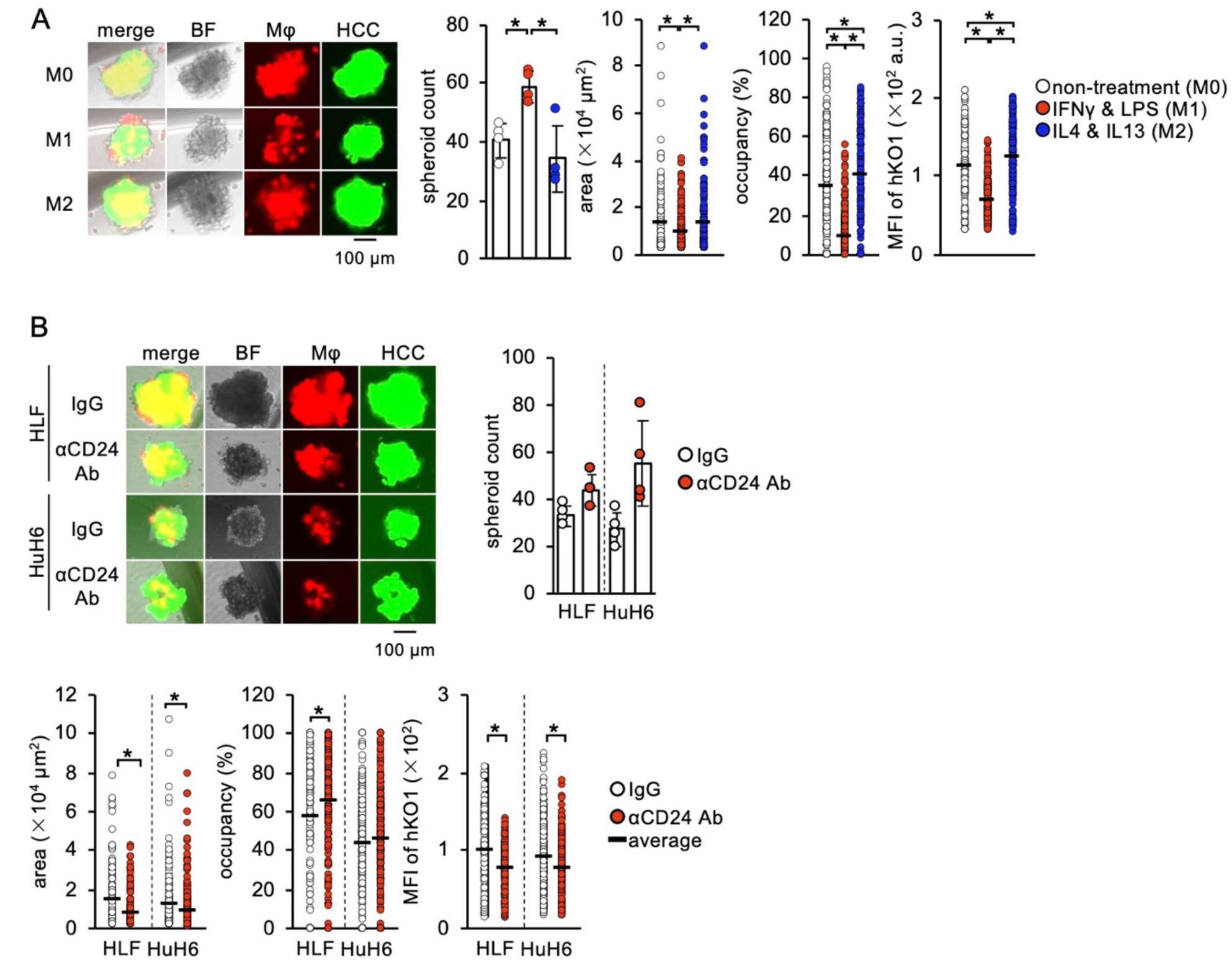
Imaging analysis of heterospheroids. (A) Representative images, count, area, occupancy, and mean fluorescence intensity (MFI) of hKO1 of heterospheroids. Heterospheroids were incubated in the absence (M0) or presence of IFNγ and LPS (M1), or IL4 and IL13 (M2) for 6 days. *, *P* < 0.05, Tukey’s test (*n* = 4 for count). (B) Representative images, count, area, occupancy, and mean fluorescence intensity (MFI) of hKO1 of the heterospheroids. LC cells were incubated with IgG or anti-CD24 antibody (αCD24 Ab) before preparing the heterospheroids. *, *P* < 0.05, Student’s *t*-test (*n* = 4 for count).

CD24 is an immune checkpoint molecule that suppresses the M1 polarization of TAMs [22]. We examined the effects of CD24 blockade on heterospheroids using two different LC cell lines, HLF and HuH6. The number of heterospheroids and macrophage occupancy tended to be increased by CD24 neutralization in both cell lines, whereas the area and MFI were significantly decreased (Figure 2B). These results suggest that M1 polarization reduces the size of heterospheroids and suppresses THP-1-derived macrophage infiltration.

### Screening for TAM activators using heterospheroids

Using the MFI of hKO1 in a heterospheroid as an indicator, we performed high-content imaging analysis to screen TAM activators using a library containing 1,134 FDA-approved drugs (Figure 3A). We found that 34 compounds decreased MFI, whereas 14 increased it without obvious toxicity (Table S2). Moreover, the second screen identified 8 of the 34 compounds that induced HLA-DR expression on macrophages in heterospheroids (Table S3); however, only alprostadil was able to reproduce this result with HLA-DR and CD80 (Figure 3B, S1). Using heterospheroids consisting of macrophages with HLF or HuH6, other PGE1-derivatives, lubiprostone, and misoprostol, exhibited less activity compared with alprostadil (Figure 3C, D). Therefore, we identified alprostadil as a TAM activator using heterospheroids.

**Figure 3.**
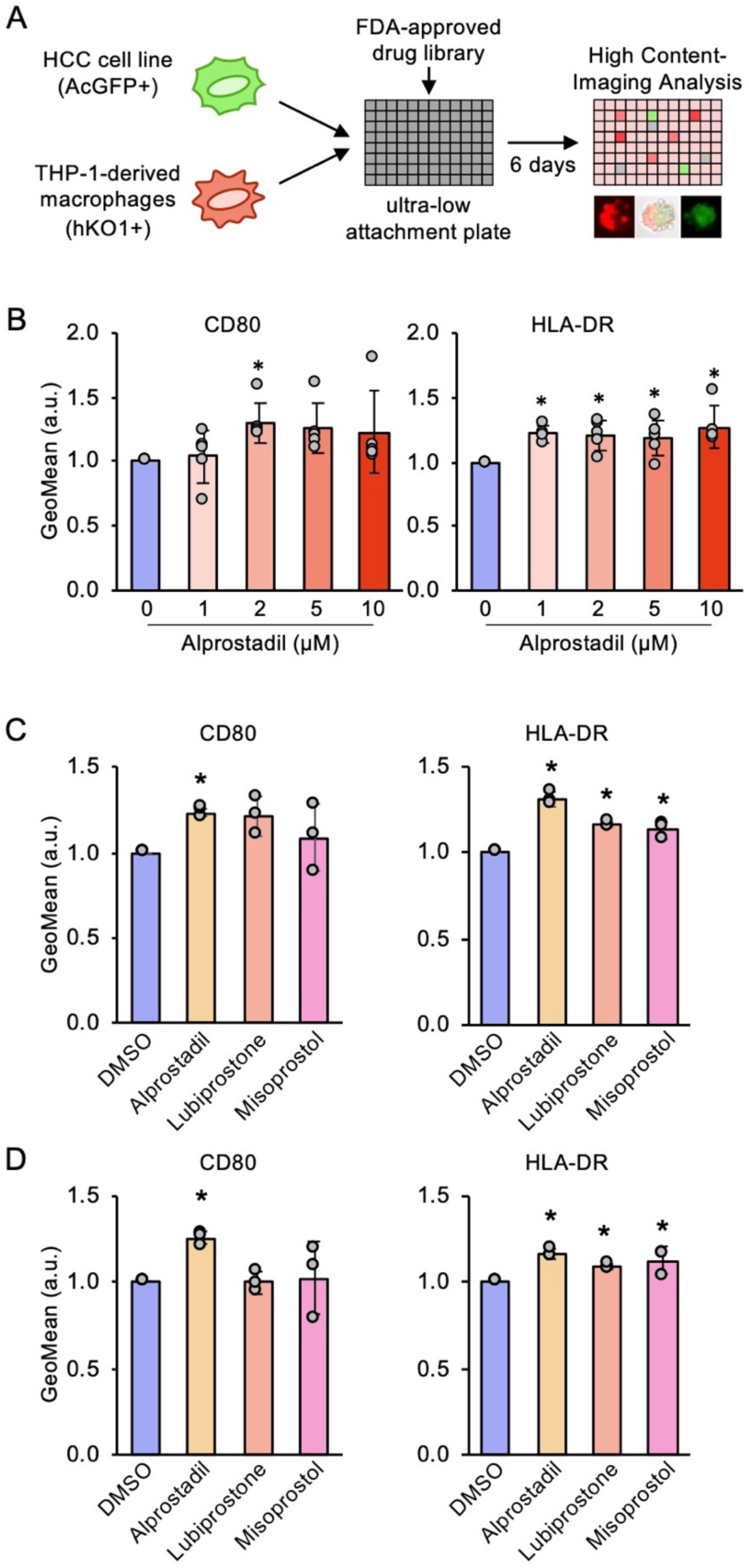
Upregulation of M1 markers by alprostadil. (A) Schematic representation of high-content imaging screening for TAM activators using heterospheroids. (B) CD80 and HLA-DR expression on THP-1-derived macrophages in heterospheroids with HLF cells. Heterospheroids were incubated with the indicated concentrations of alprostadil for 3 days. *, *P* < 0.05 vs. 0 µM, Dunnett’s test (*n* = 5). (C, D) CD80 and HLA-DR expression on THP-1-derived macrophages in heterospheroids with HLF (C) or HuH6 (D) cells. Heterospheroids were incubated with 2 µM of alprostadil, lubiprostone, or misoprostol for 3 days. *, *P* < 0.05 vs. DMSO, Dunnett’s test (*n* = 3).

### Identification of the retinoid X receptor (RXR) antagonist, HX531, as a TAM activator

After the second screening, we observed that several nuclear receptor agonists suppressed HLA-DR expression (Table S3). Consistently, treatment with all-*trans*-retinoic acid (ATRA), a potent retinoic acid receptor (RAR) agonist, suppressed M1 polarization (Figure S2). Therefore, we examined RAR and RXR antagonists. Although AR-7 (RARα antagonist), LE135 (RARα/β antagonist), and CD2665 (RARβ/γ antagonist) decreased HLA-DR expression, HX531 (RXR antagonist) significantly upregulated its expression (Figure 4A). However, other RXR antagonists (UVI3003 and PA452) and a glucocorticoid receptor antagonist, relacorilant, showed no inhibitory effects on the expression of M1 markers (Figure 4B). HX531-induced M1 marker expression was also confirmed in heterospheroids using HuH6 cells (Figure 4C).

**Figure 4.**
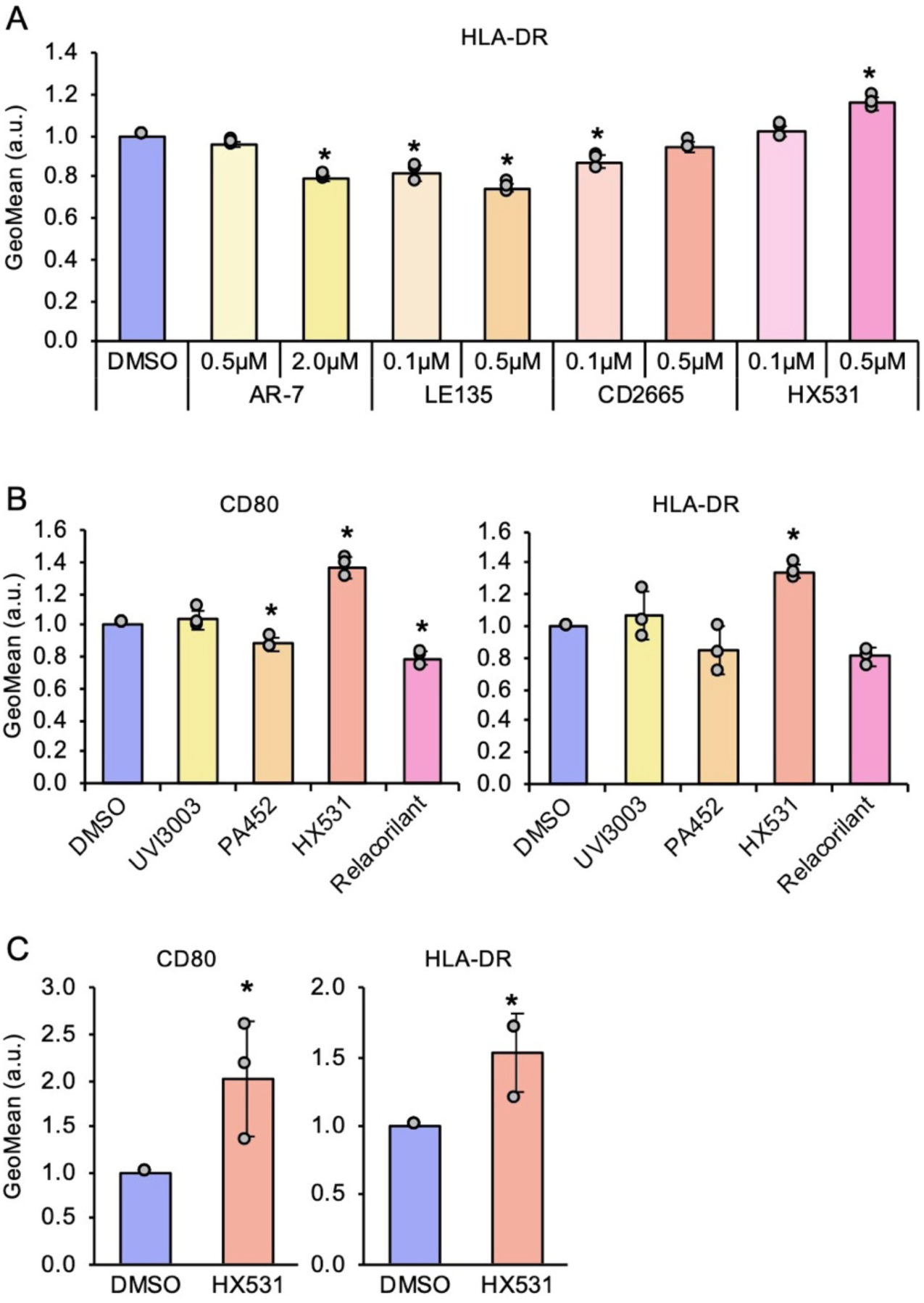
M1 marker upregulation by HX531. (A) HLA-DR expression on THP-1-derived macrophages in heterospheroids with HLF cells. Heterospheroids were incubated with the indicated concentrations of AR-7 (RARα antagonist), LE135 (RARα/β antagonist), CD2665 (RARβ/γ antagonist), or HX531 (RXR antagonist) for 3 days. *, *P* < 0.05 vs. DMSO, Dunnett’s test (*n* = 3). (B) CD80 and HLA-DR expression on THP-1-derived macrophages in heterospheroids with HLF cells. Heterospheroids were incubated with 1 µM of UVI3003 (RXR antagonist), PA452 (RXR antagonist), HX531, or relacorilant (glucocorticoid receptor antagonist) for 3 days. *, *P* < 0.05 vs. DMSO, Dunnett’s test (*n* = 3). (C) CD80 and HLA-DR expression on THP-1-derived macrophages in heterospheroids with HuH6 cells. Heterospheroids were incubated with DMSO or 1 µM HX531 for 3 days. *, *P* < 0.05 vs. DMSO, Student’s *t*-test (*n* = 3).

We further applied the heterospheroid culture to primary macrophages and examined the potential of alprostadil and HX531 as TAM activators. HLA-DR expression was significantly increased by HX531, but not by alprostadil, whereas the M2 marker, TREM2, was significantly downregulated by alprostadil, but not by HX531 (Figure 5A, B). These results suggest that although alprostadil and HX531 act as TAM activators in primary human macrophages, their actions are mediated by distinct mechanisms.

**Figure 5.**
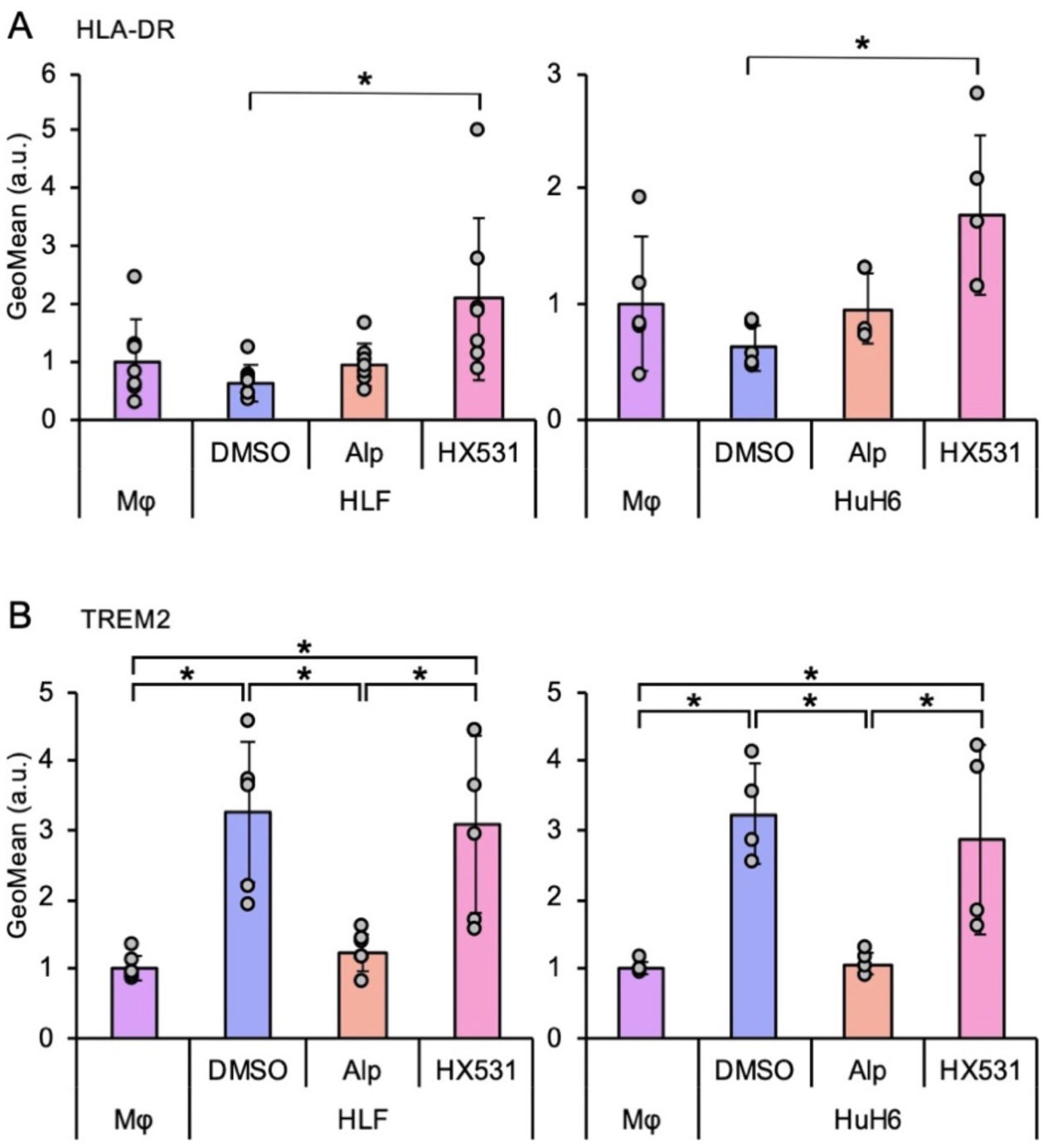
Upregulation of M1 markers by HX531. (A, B) Expression of HLA-DR (A) and TREM2 (B) on primary macrophages in homospheroids (Mφ) or heterospheroids with HLF or HuH6 cells. Heterospheroids were incubated with DMSO, 2 µM alprostadil (Alp), or 1 µM HX531. *, *P* < 0.05 vs. DMSO, Tukey’s test (*n* = 4–7).

### Transcriptome analysis of heterospheroids following HX531 treatment

Transcriptome analysis was performed to gain more insight into the mechanism of action of HX531. Treatment with HX531 for 2 days resulted in a significant difference in HLA-DR expression in heterospheroids consisting of HLF cells (Figure S3A). After a 2-day treatment with HX531, macrophages and HLF cells sorted from the heterospheroids were subjected to bulk RNA-seq. The number of differentially expressed genes (DEGs) was much larger in the macrophages compared with that in HLF cells (Figure 6 A–C, Table S4), suggesting that HX531 preferentially targets macrophages over LC cells. Pathway analyses using Gene Ontology Biological Process and Reactome revealed that the immune and interferon pathways in LC cells and cell proliferation pathways in macrophages were significantly enriched following HX531 treatment (Figure 6D, E, S4A, B, Table S5, S6). Reverse transcription-quantitative polymerase chain reaction (RT-qPCR) indicated that HX531 treatment upregulated CD80 and HLA-DRA mRNAs in macrophages (Figure S3B). In contrast, the expression of M2 markers, such as CXCR2, IL1RN, PD-L2, and TREM2, was significantly decreased in macrophages treated with HX531 (Figure S3B). PD-L2 was also downregulated in HLF cells (Figure S3B). Flow cytometry revealed that the cell surface expression of CXCR2 and TREM2 on THP-1-derived macrophages was decreased following treatment with alprostadil or HX531 (Figure S3C). iRegulon and Enrichr analyses identified several transcription factors (FOS and IRF in HLF cells; CEBPB, E2F1/4, FOXM1, and SIN3A in macrophages) (Figure S4C, Table S7, S8). The RNA-seq data indicated that the expression of E2F1, FOXM1, and KIF20A, the FOXM1-targeted gene in macrophages [23], was significantly decreased in macrophages (Table S4). RT-qPCR confirmed the downregulation of these genes in macrophages treated with HX531 (Figure S4D).

**Figure 6.**
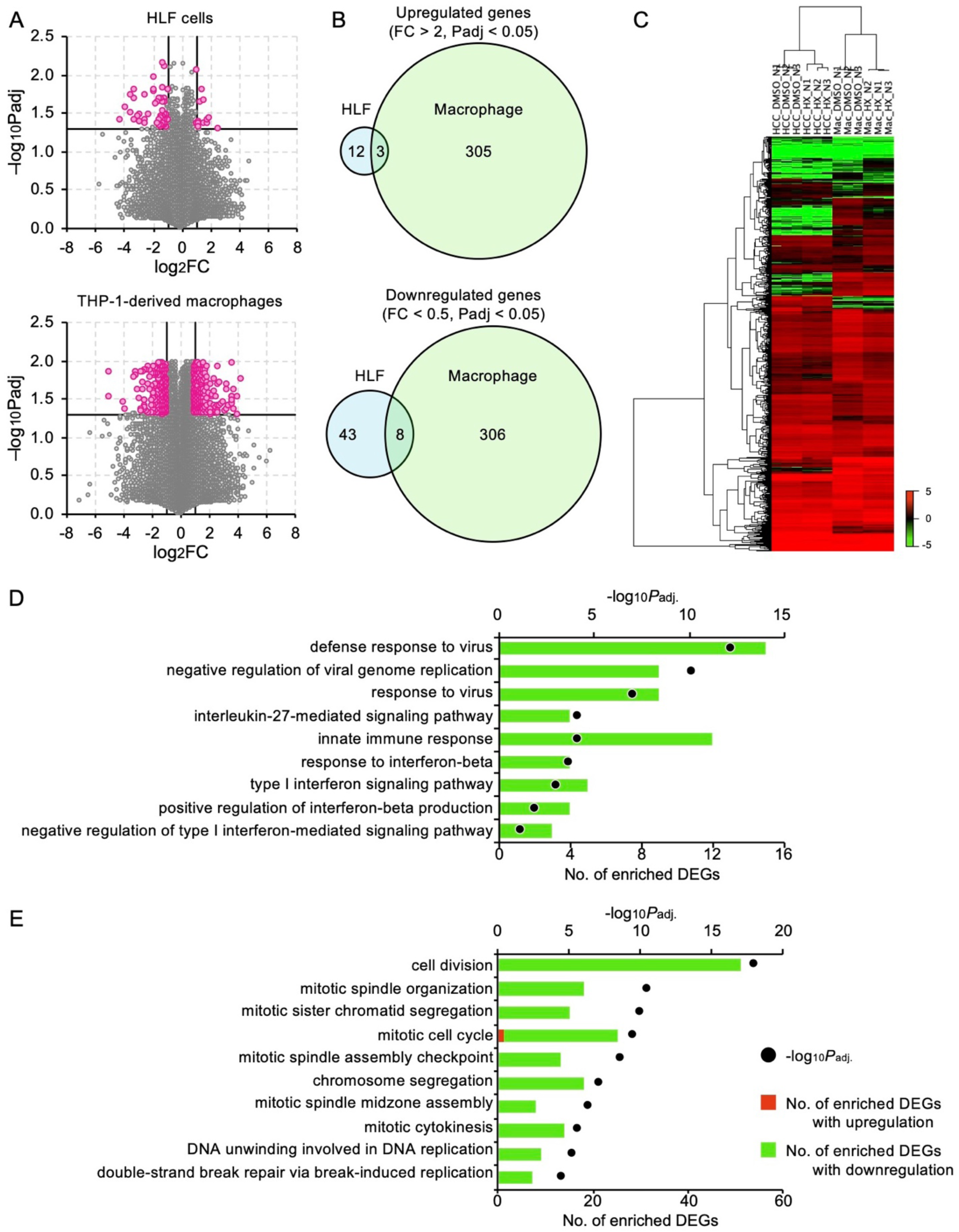
Transcriptome analysis of heterospheroids treated with HX531. THP-1-derived macrophages and HLF cells in heterospheroids treated with DMSO or 1 µM HX531 for 2 days were subjected to RNA-seq analysis (*n* = 3). (A) Volcano plots. Red dots indicate differentially expressed genes (DEGs) with |log_2_FC| > 1 and *P*_adj_ > 0.05. FC, fold change; *P*_adj_, adjusted *P* value. (B, C) Venn diagrams (B) and hierarchical cluster analysis (C) of DEGs. (D, E) GoTerm Biological Process analyses of DEGs in HLF cells (D) and macrophages (E).

### *In vivo* efficacy of alprostadil

The therapeutic efficacy of alprostadil was examined in C57BL/6 mice subcutaneously transplanted with syngeneic Hepa1-6 cells. Treatment with alprostadil (6 ng/g body weight, daily) significantly suppressed tumor growth (Figure 7A, S5A, B). In the tumor tissues, the frequencies of total and M1-polarized TAMs were not altered by alprostadil treatment (Figure 7B); however, Mhc-II expression on TAMs was significantly increased, whereas the frequency of Cd206^+^ TAMs was decreased (Figure 7B). Moreover, the frequencies of infiltrated CD45^+^ lymphocytes, particularly CD8^+^ T cells, were significantly increased, whereas those of Pd-1^+^Cd8^+^ and Cd4^+^ T cells were unchanged (Figure 7C). Treatment with anti-mouse Pd-1 antibody (100 µg/mouse, at days 0, 3, 6, 9, and 12) (Figure 7D) markedly reduced tumor volume, and when combined with alprostadil, the tumors were more rapidly eradicated compared with that in the anti-Pd-1 antibody alone (Figure 7E, S5C). This enhancement was also observed in a reduced total dose of the anti-Pd-1 antibody (days 3 and 6) (FigureS5DEF). These results indicate that alprostadil reprograms TAM polarity and enhances the efficacy of ICI therapy, which may contribute to a reduction in the frequency of ICI administration and, thus, the prevention of immune-related adverse events that are associated with ICI treatment.

**Figure 7.**
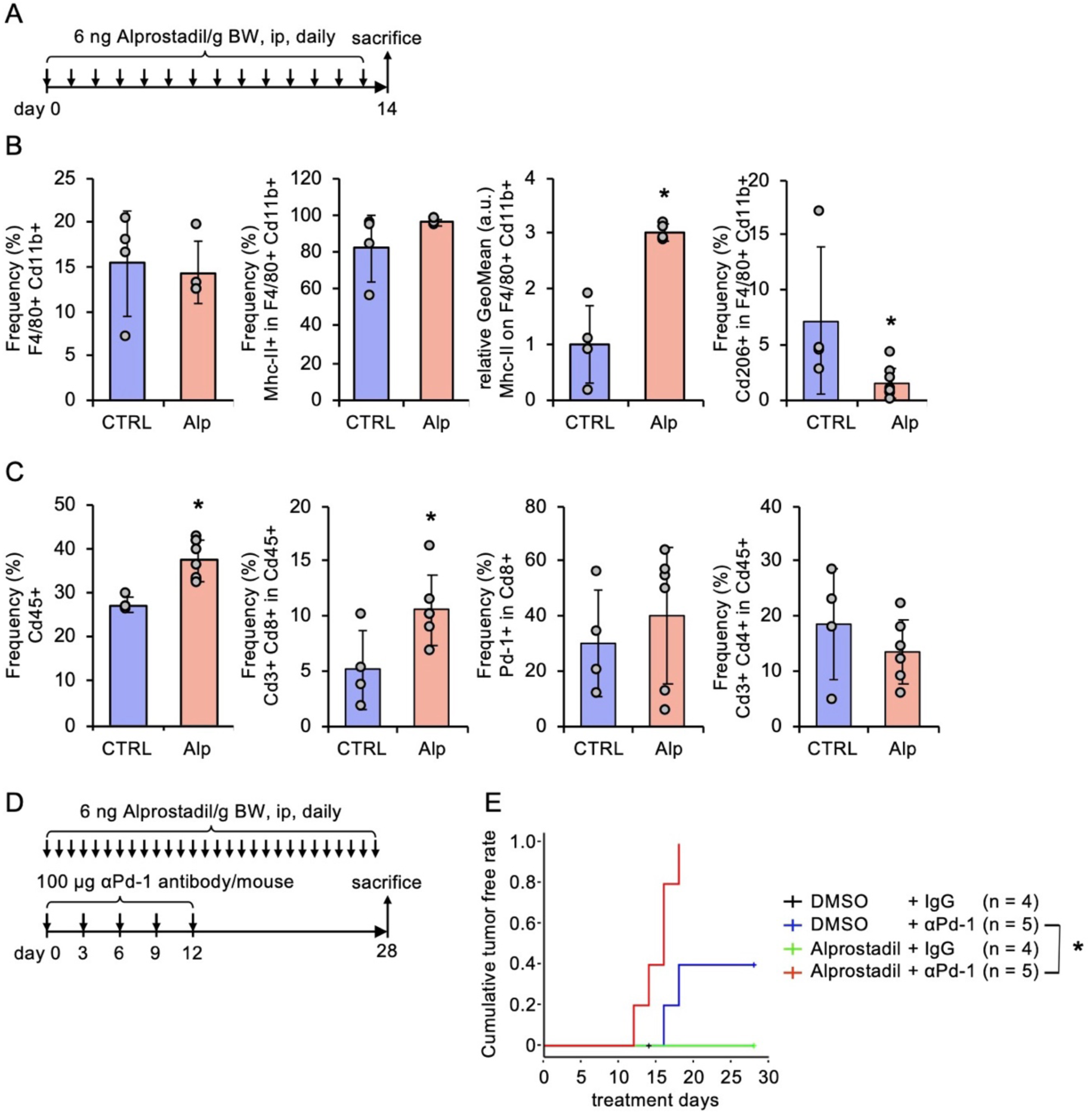
In vivo efficacy of alprostadil. (A) Treatment schedule of alprostadil. (B, C) Flow cytometry analysis of TAM (B) and lymphocytes (C) following DMSO (CTRL) or alprostadil (Alp) administration. *, *P* < 0.05 vs. CTRL, Student’s *t*-test (*n* = 4–8). (D) Treatment schedule of the combination of alprostadil and anti-Pd-1 antibody. (E) Cumulative tumor-free rate. *, *P* < 0.05, log-rank test.

## Discussion

Several mechanisms have been implicated in the polarization of TAMs. Methylenetetrahydrofolate dehydrogenase 2, which is a key enzyme in one-carbon metabolism, activates the AKT signaling pathway in macrophages, independent of its metabolic activity, to promote M2 polarization, whereas its inhibition increases M1-like macrophages and suppresses tumor growth [12]. Downregulation of xanthine oxidoreductase in TAMs promotes M2 polarization by increasing the production of the immunosuppressive metabolites adenosine and kynurenic acid [13]. A high concentration of potassium in the TME promotes the metabolic reprogramming of TAMs, resulting in M2 polarization [17]. In contrast, D-lactate produced by the gut microbiota repolarizes M2-like TAMs to the M1 phenotype through the PI3K/AKT pathway [15]. Therefore, pathways and molecules involved in M1/M2 polarization are potential targets for the development of TAM activators; however, the mechanisms underlying macrophage polarization remain unclear.

Zhou et al. established an efficient system using Il1β-reporter mouse BMDMs and performed a high-throughput screening of drugs that induce Il1β expression in BMDMs previously polarized to the M2 phenotype by IL4 [16]. They identified carfilzomib as a TAM activator and showed that it enhanced the recruitment of macrophages and CD8^+^ T cells into tumor tissues, and synergistically enhanced the efficacy of an ICI in mice with lung cancer [16]. This elegant screening system relies on the specific reporter mouse strain. Ideally, it should take the immunosuppressive effects of the TME into account. The heterospheroid culture consisting of cancer cells and tumor-associated stromal cells, has several advantages over monoculture. Cancer cells shape an immunosuppressive TME using humoral factors and direct cellular interaction, both of which play important roles in heterospheroids formation. Compared with regular cell culture, cancer cells in a 3D culture recapitulate the various features observed *in vivo*, such as cancer stemness, therapeutic resistance, and metabolic and metastatic potential. Guiet et al. showed that macrophages infiltrated into cancer cell spheroids enhanced the metastatic potential of cancer cells and promoted their release from the spheroids [24]. Herter et al. produced heterospheroids consisting of cancer cells and fibroblasts and incubated them with peripheral blood mononuclear cells [19]. They found that the heterospheroid culture enabled the assessment of the therapeutic effects of immunotherapy agents *in vitro* [19]. Moreover, Tevis et al. confirmed that heterospheroids consisting of cancer cells and macrophages promote the immunosuppressive cytokine production of macrophages and chemoresistantce of cancer cells [20]. In the present study, we demonstrated that the M1 polarization of macrophages in heterospheroids was significantly decreased compared with that in homospheroids, suggesting that an immunosuppressive TME was recapitulated *in vitro*.

It has been suggested that the widely used M2 markers, such as CD163 and CD206, are difficult to be induced in THP-1-derived macrophages by M2 inducers [25–27]. Consistently, we observed no change in the surface expression of CD163 and CD206 in the presence of IL4 and IL13. Instead, they were upregulated by IFNγ and LPS (Data not shown). Other M2 markers, including CXCR2 and TREM2, were significantly downregulated by alprostadil and HX531 (FigureS3C). CXCR2 is a receptor for CXCL1 and IL8, which exacerbates malignant phenotypes in various cancers, in part, by recruiting suppressive immune cells to the TME [28]. Currently, CXCR1/2 antagonists are undergoing clinical trials in patients with metastatic melanoma [29]. TREM2^+^ macrophages and microglial cells suppress the IFNγ-mediated anti-tumor immune response in glioblastoma and are associated with a poor prognosis [30]. TREM2^+^ macrophages may also represent a poor prognostic factor following local ablation therapy of hepatocellular carcinoma [31]. Therefore, the heterospheroid culture of THP-1-derived macrophages may be used to identify novel compounds targeting CXCR2 and TREM2, and to study their pathological functions and underlying mechanisms.

Alprostadil, also known as PGE1, exerts various physiological activities through the EP1, EP2, EP3, EP4, and IP receptors [32]. PGE2 is one of the most abundant prostaglandins in the human body. It has been shown to facilitate tumor progression by suppressing CD8^+^ T cell effector function through the EP2 and EP4 receptors on CD8^+^ T cells [33]. Moreover, EP4 antagonists are currently undergoing clinical trials as cancer treatments [34]. Therefore, alprostadil-induced EP4 activation may compromise antitumor immunity; however, our in vivo study demonstrated that alprostadil significantly promotes antitumor immunity and enhances anti-Pd-1 antibody efficacy. Because PGE2 binds to the EP1, EP2, EP3, and EP4 receptors, but not to the IP receptor [32,35], alprostadil may exert its effect through the IP receptor. In a future study, we will examine the mechanism of action of alprostadil to identify the receptor responsible for its therapeutic effects. Moreover, because we determined the antitumor effects of alprostadil only in a subcutaneous transplant model, a more precise study should be conducted in an orthotopic model or more clinically relevant models.

In the present study, we identified HX531 as a TAM activator based on the results of the second screening. Oyarce et al. previously reported HX531-induced macrophage reprogramming [36], which supports the results of our study. M2 polarization is accompanied by metabolic reprogramming orchestrated by several nuclear receptors, such as PPARγ and LXR, whose heterodimer partner is RXR [36,37]. A greater number of DEGs were identified in macrophages compared with LC cells (Figure 6B), suggesting that HX531 may target metabolic reprogramming in macrophages. However, our transcriptome analysis of macrophages revealed the enrichment of cell proliferation pathways, rather than metabolic pathways (Figure 6D, E, S4A, B, Tables S5, S6). Of note, almost all of these pathways were associated with downregulated genes, indicating that HX531 suppresses macrophage proliferation. Among the recurrent transcription factors (Figure S4C, Tables S7, S8), the expression of E2F1 and FOXM1 was significantly decreased in macrophages (Figure S4D, Tables S4). E2F1 is a well-known cell cycle regulator that promotes the transition from the G1 to S phase. Moreover, E2F1 induces IL1RN to promote the M2 polarization of THP-1-derived macrophages [38]. Therefore, HX531-induced E2F1 downregulation may inhibit cell proliferation and the downregulation of IL1RN (Figure S3B). FOXM1 promotes the M2 polarization of TAM through KIF20A [23], which was also significantly downregulated in HX531-treated macrophages (Figure S4D, Table S4). Interestingly, several studies have identified the *FOXM1* gene as a transcriptional target of E2F1 [39–41], suggesting that E2F1 is the primary target of HX531. However, because the interaction between HX531 and E2F1 has not been examined, further studies are needed to elucidate the underlying mechanism.

## Conclusions

In summary, we constructed a screening system using heterospheroids and identified alprostadil and HX531 as TAM activators. We also demonstrated that heterospheroids can provide mechanistic insight into the activity of TAM activators; for example, through transcriptome analysis. Heterospheroids provide a novel platform for developing tumor immune therapeutics and investigating oncoimmunological events through the interaction between macrophages and cancer cells (Figure S6).

## Acknowledgments

This study was partly performed at the Research Initiative Center, Tottori University. This research was partially supported by the Research Support Project for Life Science and Drug Discovery (Basis for Supporting Innovative Drug Discovery and Life Science Research (BINDS)) from AMED under Grant Number JP22ama121054.

## Data Availability

The RNA-seq data are publicly available in the DDBJ Sequence Read Archive at PRJDB35411. The other data generated in this study are available from the corresponding author upon reasonable request.

## Ethics Statement

Registry and Registration No. of the study are the isolation and culture of human peripheral blood monocytes, 22B016 (approved by the Ethical Committee of Tottori University). Informed consent was obtained from all the subjects. The animal study was approved by the Tottori University Animal Use Committee (23-Y-48).

## Author Contributions

H.T. conceptualized the project, performed most of the experiments, analyzed the data, and wrote the manuscript. M.O. performed the experiments and analyzed the data. T.H., J.Y., and M.K. contributed to the collection of blood samples and the curation of clinical information. Y.F. and D.N. supervised the project, and reviewed and edited the manuscript. All the authors have read and approved the final manuscript.

## Supplementary Description

**Supplementary Table 1. Key research resources**

**Supplementary Table 2. Summary of the first screening**

**Supplementary Table 3. Summary of the second screening**

**Supplementary Table 4. DEGs in LC cells and macrophages after HX531 treatment**

**Supplementary Table 5. Pathways predicted by GoTerm Biological Process**

**Supplementary Table 6. Pathways predicted by Reactome**

**Supplementary Table 7. Transcription factors predicted by iRegulon**

**Supplementary Table 8. Transcription factor predicted by Enrichr**

**Supplementary Figure 1. Screening of TAM activators from an FDA-approved drug library.** CD80 and HLA-DR expressions on THP-1-derived macrophages in heterospheroids with HLF cells. The heterospheroids were incubated with the indicated concentrations of each drug for 3 days. No significant difference was observed vs. 0 µM, Dunnett’s test (*n* = 3).

**Supplementary Figure 2. Downregulation of M1 markers by all-trans-retinoic acid (ATRA).** CD80 and HLA-DR expressions on THP-1-derived macrophages in heterospheroids with HLF or HuH6 cells. The heterospheroids were incubated with 10 µM of ATRA for 3 days. *, *P* < 0.05 vs. DMSO, Student’s *t*-test (*n* = 3).

**Supplementary Figure 3. Transcriptome analysis of heterospheroids treated with HX531.** (A) HLA-DR expressions on THP-1-derived macrophages in heterospheroids with HLF or HuH6 cells. The heterospheroids were incubated with DMSO or 2 µM of ATRA for the indicated days. *, *P* < 0.05 vs. DMSO, Student’s *t*-test (*n* = 3). (B) Validation of RNA-seq analysis. MRNA were recovered from THP-1-derived macrophages and HLF cells in heterospheroids treated with DMSO or 1 µM HX531 for 2 days, and subjected to RT-qPCR. *, *P* < 0.05 vs. DMSO, Student’s *t*-test (*n* = 3). (C) Cell surface CXCR2 and TREM2 expressions on THP-1-derived macrophages in heterospheroids with HLF or HuH6 cells. The heterospheroids were incubated with DMSO, 2 µM alprostadil (Alp), or 1 µM HX531 for 3 days. *, *P* < 0.05 vs. DMSO, Dunnett’s test (*n* = 4).

**Supplementary Figure 4. Pathway analysis of heterospheroids treated with HX531.** (A,B) Reactome analyses of DEGs in HLF cells (A) and macrophages (B). (C) iRegulon and Enrichr analyses of DEGs in HLF cells and macrophages. (D) Validation of iRegulon and Enrichr analyses. MRNA were recovered from THP-1-derived macrophages and HLF cells in heterospheroids treated with DMSO or 1 µM HX531 for 2 days, and subjected to RT-qPCR. *, *P* < 0.05 vs. DMSO, Student’s *t*-test (*n* = 3).

**Supplementary Figure 5. In vivo efficacy of alprostadil.** (A) Tumor volume changes in mice treated with DMSO or alprostadil, as shown in Figure 7A. (B) Representative images and weights of tumor tissues after alprostadil (Alp) treatment. *, *P* < 0.05 vs. DMSO, Student’s *t*-test (*n* = 5). (C) Tumor volume changes in each mouse treated with the combination of alprostadil and anti-Pd-1 antibody, as shown in Figure 7D. (D) Treatment schedule of combination of alprostadil and a reduced total dose of anti-Pd-1 antibody. (E,F) Tumor volume changes (E) and cumulative tumor free rate (F) of mice treated with the combination of alprostadil and anti-Pd-1 antibody, as shown in Figure S5D. *, *P* < 0.05, log-rank test.

**Supplementary Figure 6. Heterospheroids offer a new platform for oncoimmunological research and development.**

