## Supplementary for "High-throughput heterospheroid-based screening identifies drugs that reprogram tumor-associated macrophages"

**Supplementary Table 1. Key research resources**

| Reagents / Resource | Source | RRID | Authentication |
|---|---|---|---|
| **Cell Lines** | | | |
| HLF | Japanese Collection of Research Bioresources Cell Bank (August 2018) | JCRB0405 | STR (July, 2024) |
| HuH6 | Japanese Collection of Research Bioresources Cell Bank (August 2018) | JCRB0401 | STR (July, 2024) |
| HLF expressing AcGFP | Our group | N/A | STR (July, 2024) |
| HuH6 expressing AcGFP | Our group | N/A | STR (July, 2024) |
| THP-1 | RIKEN Bioresource Center (April 2021) | RCB1189 | STR (July, 2024) |
| THP-1 expressing hKO1 | Our group | N/A | STR (July, 2024) |
| Hepa 1-6 | RIKEN Bioresource Center (June 2019) | RCB1638 | STR (July, 2024) |
| **Mouse** | | | |
| C57BL/6J (male, 6 week old) | CLEA Japan | N/A | |
| **Chemicals and reagents** | | | |
| 96-well Ultra Low Attachment Multiple Well Pate | Corning | 3474 | |
| Accumax | Nacalai Tesque | 17087-54 | |
| Accutase | Nacalai Tesque | 12679-54 | |
| Albumin, Bovine Serum, General Grade, pH7.0 | Nacalai Tesque | 01860-65 | |
| All-trans retinoic acid | Santa Cruz Biotechnology | sc-200898 | |
| Alprostadil | Selleck | S1508 | |
| AR-7 | Cayman Chemical | 22946 | |
| Carbamazepine | Santa Cruz Biotechnology | SC-202518 | |
| CD2665 | Cayman Chemical | 16031 | |
| CellTracker™ CM-DiI Dye | Thermo Fisher Scientific | C7000 | |
| Chlorocresol | Tokyo Chemical Industry | C0150 | |
| Chloroxine | Sigma-Aldrich | D64600 | |
| Chlortetracycline HCl | Sigma-Aldrich | C4881-5G | |
| Dipyridamole | Sigma-Aldrich | D9766 | |
| DMEM | Shimadzu Diagnostics | 05919 | |
| EGF, Human, Recombinant, Animal Free | Peprotech | AF-100-15 | |
| Emtricitabine | Tokyo Chemical Industry | E1007 | |
| FDA-approved drug library | Center for Supporting Drug Discovery and Life Science Research, Osaka University | N/A | |
| FGF-basic(154a.a.), Human, Recombinant, Animal Free | Peprotech | AF-100-18B | |
| G418 | Nacalai Tesque | 08973-14 | |
| HX531 | Cayman Chemical | 20762 | |
| IFN-gamma, Human, Recombinant, Animal Free | Peprotech | AF-300-02 | |
| IL-13, Human, Recombinant, Animal Free | Peprotech | AF-200-13 | |
| IL-4, Human, Recombinant, Animal Free | Peprotech | AF-200-04 | |
| LE135 | Cayman Chemical | 14415 | |
| Lipopolysaccharides (LPS), from Escherichia coli O111:B4 | Sigma-Aldrich | L2630 | |
| Lubiprostone | Selleck | S1675 | |

| | | |
|---|---|---|
| M-CSF, Human, Recombinant, Animal Free | Peprotech | AF-300-25 |
| Misoprostol | Selleck | S4415 |
| Monocytes Spin Medium | pluriSelect | 60-00095-10 |
| NEBNext Poly(A) mRNA Magnetic Isolation Module | New England Biolabs | E7490 |
| NEBNext Ultra II Directional RNA Library Prep Kit for Illumina | New England Biolabs | E7760 |
| PA452 | MedChemExpress | HY-108522 |
| pAcGFP-C1 | TaKaRa Bio | 632470 |
| Phorbol 12-myristate 13-acetate (PMA) | Nacalai Tesque | 27547-14 |
| PrimeSurface shale 35 | Sumitomo Bakelite | MS-90350 |
| Relacorilant | Selleck | E1091 |
| ReverTra Ace | TOYOBO | TRT-101 |
| RPMI1640 | Shimadzu Diagnostics | 05918 |
| Sepasol-RNA I Super G | Nacalai Tesque | 09379-55 |
| THUNDERBIRD SYBR qPCR Mix | TOYOBO | QPS-201 |
| Trometamol | Sigma-Aldrich | T1503-1KG |
| UVI3003 | MedChemExpress | HY-107500 |

| **Antibodies** | **Source** | |
|---|---|---|
| *Neutralization* | | |
| Mouse IgG1κ (clone: P3.6.2.8.1) | Thermo Fisher Scientific | 14-4714-85 |
| CD24 (clone: SN3) | Thermo Fisher Scientific | MA511828 |
| Clear Back (Human Fc receptor blocking reagent) | MBL | MTG-001 |
| *Mouse treatment* | | |
| Rat IgG2aκ (clone: 2A3) | Selleck | A2123 |
| Pd-1 (clone: RMP1-14) | Selleck | A2122 |
| *Flowcytometry—human* | | |
| CD11b (clone: M1/70)—APC | BioLegend | 101212 |
| CD14 (clone: 63D3)—Alexa Fluor 647 | BioLegend | 367128 |
| CD80 (clone: 2D10.4)—APC | Thermo Fisher Scientific | 17-0809-42 |
| CD86 (clone: IT2.2)—Alexa Fluor 647 | Thermo Fisher Scientific | 51-0869-42 |
| CXCR2 (clone: 5E8/CXCR2)—Alexa Fluor 647 | BioLegend | 320714 |
| HLA-DR (clone: L243)—APC | BioLegend | 321116 |
| TREM2 (clone: 237920R)—Alexa Fluor 647 | R&D Systems | FAB17291RR |
| *Flowcytometry—mouse* | | |
| Cd45 (clone: 30-F11)—FITC | BioLegend | 103108 |
| Cd11b (clone: M1/70)—PerCP | BioLegend | 101230 |
| F4/80 (clone: BM8)—PE | BioLegend | 123110 |
| I-A/I-E (clone: M5/114.15.2)—APC | BioLegend | 107614 |
| Cd206 (clone: C068C2)—APC | BioLegend | 141708 |
| Cd3ε (clone: 145-2C11)—PerCP | BioLegend | 100326 |
| Cd4 (clone: GK1.5)—PE | BioLegend | 100408 |
| Cd8a (clone: 53-6.7)—PE | BioLegend | 100708 |
| Pd-1 (clone: RMP1-30)—Alexa Fluor 647 | BioLegend | 109118 |

| Primers | Sequence | Reference sequence |
|---|---|---|
| 36B4, antisense | TCAGCATGTTCAGCAGTGTG | NM_007475.5 |
| 36B4, sense | TGGGCATCACCACGAAAATC | |
| CD80, antisense | AGCGTTGCCACTTCTTTCAC | NM_005191.4 |
| CD80, sense | TCACCATCCAAGTGTCCATACC | |
| CXCR2, antisense | TGGGCTTTTCACCTGTAGGAC | NM_001557.4 |
| CXCR2, sense | ACCTTGAGGCACAGTGAAGAC | |
| E2F1, antisense | AAAACATCGATCGGGCCTTG | NM_005225.3 |
| E2F1, sense | ACCTTCGTAGCATTGCAGAC | |
| FOXM1, antisense | AACATGTCGTGCAGGGAAAG | NM_202002.3 |
| FOXM1, sense | ATTCGCCATCAACAGCACTG | |
| HLA-DRA, antisense | TGGTGGGGTGAACTTGTCTATG | NM_019111.5 |
| HLA-DRA, sense | TCATGACAAAGCGCTCCAAC | |
| IL1RA, antisense | GCTTGTCCTGCTTTCTGTTCTC | NM_173841.3 |
| IL1RN, sense | TCCTGTGTCAAGTCTGGTGATG | |
| KIF20A, antisense | GCCTGTGAAGAAACCTTGGAAC | NM_005733.3 |
| KIF20A, sense | TTGCTGCCCTTCGTCAAAAC | |
| PDL2, antisense | ATGTGAAGCAGCCAAGTTGG | NM_025239.4 |
| PDL2, sense | TGGCAGAAACTTCAGCTGTG | |
| TREM2, antisense | AGTGCTTCATGGAGTCATAGGG | NM_018965.4 |
| TREM2, sense | CGGCTGCTCATCTTACTCTTTG | |

**Supplementary Table 2. Summary of the first screening**

| Compounds | MFI of hKO1[1),2)] n = 1 | n = 2 | mean | toxicity[3)] |
|---|---|---|---|---|
| Afatinib (BIBW2992) | 1.32 | 1.19 | 1.26 | toxic |
| Bortezomib (PS-341) | 0.00 | 0.59 | 0.30 | toxic |
| Everolimus (RAD001) | 0.78 | 0.69 | 0.73 | |
| Dasatinib | 1.53 | 1.02 | 1.27 | toxic |
| Erlotinib HCl (OSI-744) | 1.44 | 1.26 | 1.35 | |
| Gefitinib (ZD1839) | 1.47 | 1.20 | 1.33 | |
| Imatinib Mesylate (STI571) | 0.86 | 1.07 | 0.96 | |
| Lapatinib (GW-572016) Ditosylate | 1.00 | 0.96 | 0.98 | |
| Lenalidomide (CC-5013) | 1.01 | 0.99 | 1.00 | |
| Nilotinib (AMN-107) | 0.99 | 1.00 | 0.99 | |
| Rapamycin (Sirolimus) | 0.94 | 0.79 | 0.86 | |
| Sorafenib Tosylate | 0.44 | 0.90 | 0.67 | |
| Sunitinib Malate | 0.61 | 0.89 | 0.75 | |
| Temsirolimus (CCI-779, NSC 683864) | 0.69 | 1.07 | 0.88 | |
| Vandetanib (ZD6474) | 1.09 | 1.05 | 1.07 | |
| Vorinostat (SAHA, MK0683) | 1.33 | 1.21 | 1.27 | toxic |
| Masitinib (AB1010) | 1.20 | 0.93 | 1.07 | |
| Crizotinib (PF-02341066) | 0.96 | 1.01 | 0.98 | |
| Vismodegib (GDC-0449) | 0.95 | 0.96 | 0.96 | |
| Cabozantinib (XL184, BMS-907351) | 0.82 | 0.94 | 0.88 | |
| Abiraterone (CB-7598) | 0.81 | 0.81 | 0.81 | |
| Pemetrexed | 1.10 | 1.11 | 1.10 | |
| Malotilate | 0.98 | 0.91 | 0.94 | |
| Ivacaftor (VX-770) | 1.17 | 0.65 | 0.91 | |
| Docetaxel | 1.29 | 1.18 | 1.24 | |
| Gemcitabine HCl (Gemzar) | 1.26 | 1.24 | 1.25 | |
| Paclitaxel | 1.35 | 1.08 | 1.22 | |
| Capecitabine | 0.97 | 1.03 | 1.00 | |
| Cisplatin | 0.99 | 1.11 | 1.05 | |
| Valproic acid sodium salt (Sodium valproate) | 0.97 | 1.10 | 1.03 | |
| Ritonavir | 0.99 | 0.97 | 0.98 | |
| Axitinib | 0.78 | 0.87 | 0.83 | |
| Aprepitant | 1.00 | 1.07 | 1.03 | |
| Regorafenib (BAY 73-4506) | 0.70 | 0.53 | 0.61 | toxic |
| Fulvestrant | 0.98 | 1.00 | 0.99 | |
| Thalidomide | 1.02 | 0.80 | 0.91 | |
| Exemestane | 0.96 | 1.05 | 1.01 | |
| Finasteride | 1.11 | 0.98 | 1.04 | |
| Irinotecan | 1.22 | 1.10 | 1.16 | |
| Cladribine | 1.04 | 1.11 | 1.08 | |
| Dutasteride | 0.95 | 1.28 | 1.12 | |
| Melatonin | 1.01 | 0.97 | 0.99 | |
| Bisoprolol fumarate | 0.98 | 1.11 | 1.04 | |
| Doxorubicin (Adriamycin) | 1.76 | 0.86 | 1.31 | toxic |
| Fluorouracil (5-Fluoracil, 5-FU) | 1.10 | 1.22 | 1.16 | |
| Methothrexate | 1.34 | 1.17 | 1.26 | |
| Imiquimod | 1.04 | 1.04 | 1.04 | |
| Bendamustine HCl | 1.02 | 1.10 | 1.06 | |
| Nelarabine | 1.04 | 1.16 | 1.10 | |
| Bleomycin Sulfate | 1.04 | 1.13 | 1.08 | |
| Clafen (Cyclophosphamide) | 0.97 | 0.95 | 0.96 | |
| Clofarabine | 1.57 | 1.17 | 1.37 | toxic |
| Dacarbazine | 1.01 | 0.90 | 0.95 | |
| Dexrazoxane HCl (ICRF-187, ADR-529) | 1.17 | 0.94 | 1.05 | |
| Epirubicin HCl | 1.67 | 0.87 | 1.27 | toxic |
| Decitabine | 1.05 | 0.86 | 0.96 | |
| Etoposide | 1.10 | 1.12 | 1.11 | |
| Raloxifene HCl | 0.89 | 1.13 | 1.01 | |

| | | | | |
|---|---|---|---|---|
| Idarubicin HCl | 1.38 | 0.94 | 1.16 | |
| Fludarabine Phosphate | 2.08 | 1.00 | 1.54 | toxic |
| 2-Methoxyestradiol (2-MeOE2) | 1.10 | 1.01 | 1.05 | |
| Pazopanib HCl (GW786034 HCl) | 1.06 | 0.92 | 0.99 | |
| Leucovorin Calcium | 1.06 | 1.02 | 1.04 | |
| Temozolomide | 1.01 | 0.95 | 0.98 | |
| Vincristine | 0.58 | 0.59 | 0.59 | toxic |
| Agomelatine | 1.07 | 1.28 | 1.17 | |
| Leflunomide | 0.86 | 0.96 | 0.91 | |
| Vinblastine | 0.81 | 0.70 | 0.75 | toxic |
| Carboplatin | 0.94 | 1.00 | 0.97 | |
| Topotecan HCl | 1.56 | 0.85 | 1.21 | |
| Nepafenac | 1.07 | 1.09 | 1.08 | |
| Rufinamide | 1.09 | 0.97 | 1.03 | |
| Posaconazole | 0.96 | 0.97 | 0.96 | |
| Prasugrel | 1.12 | 0.99 | 1.06 | |
| Ramelteon | 1.07 | 1.11 | 1.09 | |
| Cinacalcet HCl | 1.00 | 1.13 | 1.07 | |
| Celecoxib | 1.02 | 1.07 | 1.05 | |
| Vemurafenib (PLX4032, RG7204) | 1.14 | 1.18 | 1.16 | |
| Acarbose | 1.12 | 1.04 | 1.08 | |
| Adapalene | 0.84 | 0.77 | 0.81 | |
| Amisulpride | 1.11 | 1.04 | 1.08 | |
| Aniracetam | 1.20 | 0.86 | 1.03 | |
| Artemisinin | 1.12 | 0.78 | 0.95 | |
| Asenapine | 1.09 | 1.07 | 1.08 | |
| Benazepril HCl | 1.24 | 1.05 | 1.15 | |
| Biperiden HCl | 1.15 | 0.66 | 0.90 | |
| Budesonide | 1.33 | 0.91 | 1.12 | |
| Bumetanide | 1.26 | 0.99 | 1.13 | |
| Camptothecin | 1.70 | 1.11 | 1.40 | toxic |
| Carmofur | 1.65 | 1.18 | 1.42 | |
| Cilnidipine | 1.09 | 0.94 | 1.02 | |
| Cilostazol | 1.15 | 1.05 | 1.10 | |
| Floxuridine | 1.33 | 1.21 | 1.27 | |
| FT-207 (NSC 148958) | 1.12 | 0.95 | 1.03 | |
| Ifosfamide | 1.13 | 1.13 | 1.13 | |
| Megestrol Acetate | 1.20 | 1.03 | 1.11 | |
| Mercaptopurine (6-MP) | 1.18 | 0.91 | 1.04 | |
| Pamidronate Disodium | 1.14 | 0.97 | 1.05 | |
| Streptozotocin (STZ) | 1.10 | 1.11 | 1.11 | |
| Zoledronic Acid | 1.17 | 1.05 | 1.11 | |
| Doxazosin Mesylate | 1.02 | 1.07 | 1.05 | |
| Edaravone | 1.15 | 0.93 | 1.04 | |
| Ellagic acid | 1.11 | 0.95 | 1.03 | |
| Etodolac | 1.15 | 1.04 | 1.10 | |
| Etomidate | 1.16 | 1.03 | 1.10 | |
| Felbamate | 1.19 | 0.95 | 1.07 | |
| Fluconazole | 1.02 | 0.99 | 1.01 | |
| Flumazenil | 1.21 | 1.02 | 1.11 | |
| Fluoxetine HCl | 1.15 | 0.95 | 1.05 | |
| Fluvoxamine maleate | 1.12 | 0.98 | 1.05 | |
| Gatifloxacin | 0.99 | 0.89 | 0.94 | |
| Genistein | 1.09 | 1.17 | 1.13 | |
| Glimepiride | 1.17 | 1.11 | 1.14 | |
| Granisetron HCl | 1.15 | 0.86 | 1.01 | |
| Ivermectin | 1.10 | 0.87 | 0.98 | |
| Ketoconazole | 1.13 | 0.99 | 1.06 | |
| Lansoprazole | 1.09 | 1.02 | 1.05 | |
| Levetiracetam | 1.10 | 1.10 | 1.10 | |
| Lidocaine | 1.15 | 1.19 | 1.17 | |
| Loratadine | 1.09 | 1.18 | 1.13 | |

| | | | |
|---|---|---|---|
| Acitretin | 0.69 | 0.55 | 0.62 |
| Biapenem | 1.11 | 1.03 | 1.07 |
| Cefoselis Sulfate | 1.06 | 0.89 | 0.97 |
| Daptomycin | 1.14 | 0.96 | 1.05 |
| Doripenem Hydrate | 1.14 | 1.09 | 1.12 |
| Dorzolamide HCL | 1.10 | 1.09 | 1.10 |
| Gestodene | 1.14 | 1.06 | 1.10 |
| Drospirenone | 1.22 | 0.96 | 1.09 |
| Altretamine | 1.17 | 1.12 | 1.14 |
| Isotretinoin | 0.84 | 0.73 | 0.79 |
| Meropenem | 0.86 | 0.96 | 0.91 |
| Mianserin HCl | 1.13 | 0.90 | 1.01 |
| Minoxidil | 1.19 | 0.82 | 1.00 |
| Mizoribine | 1.14 | 0.94 | 1.04 |
| Mosapride Citrate | 1.14 | 0.95 | 1.04 |
| Nafamostat Mesylate | 1.29 | 0.97 | 1.13 |
| Omeprazole | 1.15 | 0.91 | 1.03 |
| Ondansetron HCl | 1.13 | 1.11 | 1.12 |
| Oxcarbazepine | 1.14 | 0.88 | 1.01 |
| Pizotifen Malate | 1.23 | 1.11 | 1.17 |
| Rocuronium Bromide | 1.03 | 1.01 | 1.02 |
| Stavudine (d4T) | 1.16 | 0.97 | 1.06 |
| Teicoplanin | 1.15 | 0.93 | 1.04 |
| Tenofovir Disoproxil Fumarate | 1.05 | 1.04 | 1.04 |
| Tenofovir | 1.11 | 0.98 | 1.05 |
| Tigecycline | 1.14 | 1.07 | 1.11 |
| Entecavir Hydrate | 1.22 | 1.02 | 1.12 |
| Vecuronium Bromide | 1.18 | 1.05 | 1.12 |
| Bimatoprost | 1.14 | 1.03 | 1.08 |
| Linezolid | 1.15 | 1.07 | 1.11 |
| Clopidogrel | 1.15 | 1.10 | 1.13 |
| Prazosin HCl | 1.14 | 0.92 | 1.03 |
| Ranolazine 2HCl | 1.23 | 1.06 | 1.14 |
| Repaglinide | 1.14 | 1.10 | 1.12 |
| Risedronate sodium | 1.20 | 1.11 | 1.16 |
| Rolipram | 1.20 | 1.03 | 1.12 |
| Sildenafil Citrate | 1.18 | 1.01 | 1.09 |
| Sumatriptan Succinate | 1.15 | 1.00 | 1.07 |
| Tianeptine sodium | 1.15 | 0.94 | 1.05 |
| Tizanidine HCl | 1.21 | 1.08 | 1.14 |
| Tranilast | 1.07 | 1.07 | 1.07 |
| Varenicline Tartrate | 1.04 | 1.06 | 1.05 |
| Venlafaxine | 0.95 | 1.14 | 1.05 |
| Voriconazole | 0.89 | 0.95 | 0.92 |
| Zileuton | 1.08 | 1.04 | 1.06 |
| Ziprasidone HCl | 1.01 | 1.02 | 1.01 |
| Zonisamide | 1.03 | 1.05 | 1.04 |
| Atazanavir Sulfate | 1.03 | 1.15 | 1.09 |
| Ofloxacin (Floxin) | 0.99 | 1.00 | 1.00 |
| Marbofloxacin | 1.05 | 1.12 | 1.08 |
| Calcitriol | 0.78 | 0.71 | 0.74 |
| Doxercalciferol | 0.79 | 0.98 | 0.89 |
| Alfacalcidol | 1.06 | 1.04 | 1.05 |
| Calcifediol | 0.65 | 0.88 | 0.76 |
| Iloperidone | 0.90 | 0.89 | 0.89 |
| Naratriptan | 0.92 | 1.07 | 1.00 |
| Ponatinib (AP24534) | 0.79 | 0.84 | 0.81 |
| Fludarabine (Fludara) | 1.49 | 0.99 | 1.24 |
| Pralatrexate | 1.29 | 1.16 | 1.23 |
| Cefaclor (Ceclor) | 1.07 | 1.07 | 1.07 |
| Mycophenolate Mofetil | 0.91 | 1.01 | 0.96 |
| Cephalexin | 0.95 | 0.97 | 0.96 |

| | | | |
|---|---|---|---|
| Dyphylline | 0.89 | 1.05 | 0.97 |
| Aztreonam | 1.00 | 1.13 | 1.06 |
| Perindopril Erbumine (Aceon) | 1.06 | 0.98 | 1.02 |
| Irbesartan | 0.92 | 0.89 | 0.90 |
| Alprostadil | 0.69 | 0.83 | 0.76 |
| Norfloxacin (Norxacin) | 0.99 | 1.07 | 1.03 |
| Tadalafil | 0.96 | 1.02 | 0.99 |
| Cyclosporine | 1.00 | 1.05 | 1.02 |
| Natamycin | 0.81 | 1.10 | 0.96 |
| Ibuprofen Lysine (NeoProfen) | 0.90 | 1.02 | 0.96 |
| Vinorelbine (Navelbine) | 0.85 | 0.77 | 0.81 |
| Telaprevir (VX-950) | 0.89 | 0.91 | 0.90 |
| Cetirizine DiHCl | 1.01 | 0.91 | 0.96 |
| Dexamethasone (DHAP) | 0.84 | 0.97 | 0.91 |
| Nebivolol | 0.98 | 0.90 | 0.94 |
| Pimobendan | 0.78 | 0.90 | 0.84 |
| Pomalidomide | 0.90 | 1.09 | 1.00 |
| Tazarotene | 0.67 | 0.81 | 0.74 |
| Candesartan | 0.95 | 0.94 | 0.95 |
| Ubenimex (Bestatin) | 0.82 | 0.95 | 0.89 |
| Apixaban | 0.96 | 1.01 | 0.98 |
| Reserpine | 0.90 | 0.99 | 0.94 |
| Furosemide | 0.93 | 0.98 | 0.95 |
| Olmesartan Medoxomil | 0.99 | 0.97 | 0.98 |
| Cefdinir | 0.96 | 0.96 | 0.96 |
| Clotrimazole | 0.94 | 1.14 | 1.04 |
| Rizatriptan Benzoate | 0.92 | 1.06 | 0.99 |
| Pyridostigmine Bromide | 0.95 | 1.12 | 1.03 |
| Metolazone | 0.84 | 1.02 | 0.93 |
| Cefoperazone | 1.04 | 0.94 | 0.99 |
| Silodosin | 1.02 | 0.93 | 0.97 |
| Riluzole | 0.92 | 1.05 | 0.98 |
| Risperidone | 0.90 | 0.84 | 0.87 |
| Sulfapyridine | 0.99 | 0.94 | 0.96 |
| Sulfameter | 0.99 | 1.04 | 1.01 |
| Prilocaine | 0.91 | 1.02 | 0.96 |
| Darunavir Ethanolate | 1.00 | 1.06 | 1.03 |
| Prednisone | 0.87 | 1.05 | 0.96 |
| Alendronate (Fosamax) | 0.90 | 1.03 | 0.97 |
| Ethinyl Estradiol | 0.97 | 0.96 | 0.96 |
| Naproxen | 1.03 | 0.97 | 1.00 |
| Nitazoxanide | 1.09 | 1.15 | 1.12 |
| Triamcinolone Acetonide | 1.20 | 0.96 | 1.08 |
| Orlistat | 1.02 | 0.98 | 1.00 |
| Allopurinol | 0.95 | 0.98 | 0.97 |
| Zafirlukast | 1.05 | 0.89 | 0.97 |
| Erythromycin | 0.89 | 1.10 | 0.99 |
| Amphotericin B | 0.93 | 1.03 | 0.98 |
| Ibuprofen | 1.12 | 0.99 | 1.05 |
| Amprenavir | 0.89 | 0.97 | 0.93 |
| Albendazole | 1.07 | 1.11 | 1.09 |
| Chlorothiazide | 0.99 | 0.98 | 0.99 |
| Methyldopa (Aldomet) | 1.03 | 0.87 | 0.95 |
| Ursodiol | 1.00 | 1.14 | 1.07 |
| Nitrofural | 0.96 | 0.97 | 0.96 |
| Ketoprofen | 1.00 | 1.24 | 1.12 |
| Ketorolac | 1.05 | 1.13 | 1.09 |
| Adenosine | 1.07 | 1.18 | 1.12 |
| Zolmitriptan | 1.07 | 0.90 | 0.99 |
| Telbivudine | 1.03 | 0.95 | 0.99 |
| Monobenzone | 1.00 | 1.10 | 1.05 |
| Tretinoin | 0.81 | 0.90 | 0.85 |

| | | | | |
|---|---|---|---|---|
| Phenylbutazone | 0.98 | 1.04 | 1.01 | |
| Ezetimibe | 1.01 | 0.93 | 0.97 | |
| Enalaprilat Dihydrate | 1.04 | 1.08 | 1.06 | |
| Dofetilide | 1.13 | 1.09 | 1.11 | |
| Isradipine | 1.18 | 1.02 | 1.10 | |
| Estrone | 1.01 | 1.04 | 1.03 | |
| Trichlormethiazide | 1.06 | 0.97 | 1.02 | |
| Loteprednol etabonate | 1.02 | 0.94 | 0.98 | |
| （6-） ε- ? Aminocaproic acid | 0.97 | 1.03 | 1.00 | |
| Aminoglutethimide | 1.01 | 0.98 | 1.00 | |
| Aminophylline | 0.96 | 1.00 | 0.98 | |
| Amorolfine HCl | 0.96 | 0.97 | 0.96 | |
| Chloramphenicol | 1.06 | 1.02 | 1.04 | |
| Flurbiprofen | 0.98 | 0.99 | 0.98 | |
| Disulfiram | 0.95 | 1.02 | 0.99 | |
| Mesalamine | 0.97 | 1.10 | 1.04 | |
| Sulfanilamide | 1.03 | 0.93 | 0.98 | |
| Betamethasone Dipropionate (Diprolene) | 1.11 | 1.00 | 1.06 | |
| Meprednisone | 0.96 | 0.92 | 0.94 | |
| Betamethasone valerate (Betnovate) | 1.00 | 1.07 | 1.04 | |
| Praziquantel | 0.98 | 1.02 | 1.00 | |
| Busulfan | 1.04 | 1.00 | 1.02 | |
| Carbamazepine | 0.80 | 0.79 | 0.80 | |
| Hydrocortisone | 1.03 | 0.89 | 0.96 | |
| Torsemide | 1.12 | 0.98 | 1.05 | |
| Desonide | 1.15 | 0.97 | 1.06 | |
| Divalproex Sodium | 0.99 | 0.93 | 0.96 | |
| Emtricitabine | 0.80 | 0.80 | 0.80 | |
| Progesterone | 0.89 | 0.98 | 0.94 | |
| Lamivudine | 1.02 | 0.94 | 0.98 | |
| Eplerenone | 0.97 | 0.92 | 0.94 | |
| Hydrochlorothiazide | 1.02 | 0.99 | 1.00 | |
| Estradiol | 1.03 | 0.98 | 1.00 | |
| Deferasirox | 0.98 | 1.00 | 0.99 | |
| Piroxicam | 1.06 | 1.03 | 1.04 | |
| Gemcitabine | 1.57 | 1.16 | 1.37 | toxic |
| Glyburide | 0.95 | 1.04 | 0.99 | |
| Adefovir Dipivoxil | 0.77 | 0.83 | 0.80 | |
| Zalcitabine | 0.86 | 0.93 | 0.89 | |
| Azathioprine | 0.89 | 0.98 | 0.93 | |
| Indomethacin | 0.97 | 1.00 | 0.99 | |
| Paliperidone (Invega) | 0.98 | 0.81 | 0.89 | |
| Terbinafine | 0.97 | 1.00 | 0.99 | |
| Levodopa (Sinemet) | 0.85 | 1.02 | 0.93 | |
| Levonorgestrel | 0.99 | 0.94 | 0.96 | |
| Gemfibrozil | 0.98 | 1.07 | 1.03 | |
| Mitotane | 0.94 | 1.04 | 0.99 | |
| Methylprednisolone | 1.01 | 0.90 | 0.96 | |
| Meloxicam | 1.07 | 0.92 | 0.99 | |
| Mesna | 0.94 | 0.87 | 0.90 | |
| Methocarbamol | 0.90 | 1.00 | 0.95 | |
| Prednisolone | 1.02 | 0.99 | 1.00 | |
| Telmisartan | 1.03 | 0.97 | 1.00 | |
| Thiabendazole | 1.05 | 0.94 | 1.00 | |
| Guaifenesin | 1.09 | 0.97 | 1.03 | |
| Rifabutin | 0.99 | 1.08 | 1.03 | |
| Esomeprazole magnesium (Nexium) | 0.92 | 0.93 | 0.92 | |
| Nicotinic Acid | 0.97 | 0.96 | 0.97 | |
| Nimodipine | 1.03 | 0.84 | 0.93 | |
| Nisoldipine | 1.02 | 1.07 | 1.04 | |
| L-Glutamine | 0.95 | 0.93 | 0.94 | |
| Gadodiamide | 0.87 | 0.90 | 0.89 | |

| | | | |
|---|---|---|---|
| Oxybutynin (Ditropan) | 1.06 | 0.96 | 1.01 |
| Enoxacin | 1.11 | 1.03 | 1.07 |
| Pitavastatin Calcium | 1.22 | 0.76 | 0.99 |
| Rifapentine | 1.08 | 0.86 | 0.97 |
| Pyrazinamide | 0.99 | 0.95 | 0.97 |
| Quetiapine Fumarate | 0.98 | 0.92 | 0.95 |
| Rifampin | 1.01 | 1.06 | 1.04 |
| Beta Carotene | 0.95 | 1.03 | 0.99 |
| Cefditoren Pivoxil | 0.96 | 1.02 | 0.99 |
| Sulfadiazine | 0.98 | 0.98 | 0.98 |
| Chlorprothixene | 0.94 | 0.90 | 0.92 |
| Oxytetracycline (Terramycin) | 1.03 | 0.97 | 1.00 |
| Thioguanine | 0.98 | 1.09 | 1.03 |
| Toremifene Citrate | 1.10 | 1.02 | 1.06 |
| Trifluridine | 0.85 | 0.87 | 0.86 |
| Azacitidine | 0.86 | 0.92 | 0.89 |
| Vidarabine | 0.97 | 1.00 | 0.98 |
| Verteporfin (Visudyne) | 0.84 | 1.03 | 0.93 |
| Teniposide | 1.20 | 1.15 | 1.17 |
| Rifaximin | 0.80 | 0.89 | 0.85 |
| Losartan Potassium (DuP 753) | 0.95 | 0.89 | 0.92 |
| Simvastatin | 1.03 | 1.14 | 1.08 |
| Ramipril | 0.92 | 0.97 | 0.95 |
| Fenofibrate | 0.87 | 1.05 | 0.96 |
| Ranitidine | 0.96 | 0.94 | 0.95 |
| Acadesine | 1.01 | 0.84 | 0.93 |
| Acetylcholine Chloride | 0.79 | 0.92 | 0.85 |
| Acipimox | 0.82 | 0.99 | 0.90 |
| Aciclovir | 0.81 | 0.84 | 0.82 |
| Nifedipine | 0.81 | 0.89 | 0.85 |
| Amiloride HCl | 0.89 | 0.88 | 0.88 |
| Amlodipine besylate (Norvasc) | 0.90 | 0.96 | 0.93 |
| Chlorpheniramine Maleate | 0.79 | 0.92 | 0.85 |
| Clofibrate (Atromid-S) | 1.06 | 0.97 | 1.02 |
| Erdosteine | 0.92 | 0.93 | 0.93 |
| Betaxolol hydrochloride (Betoptic) | 0.92 | 0.99 | 0.96 |
| Proparacaine HCl | 0.83 | 0.86 | 0.84 |
| Pranlukast | 0.93 | 0.87 | 0.90 |
| Oxfendazole | 0.75 | 0.98 | 0.87 |
| Carvedilol | 0.85 | 0.89 | 0.87 |
| Atracurium Besylate | 1.02 | 0.99 | 1.00 |
| Butoconazole nitrate | 0.92 | 1.04 | 0.98 |
| Azithromycin | 0.87 | 1.07 | 0.97 |
| Albendazole Oxide | 0.98 | 0.93 | 0.96 |
| Chloroxine | 0.84 | 0.67 | 0.75 |
| Lomustine | 0.77 | 0.95 | 0.86 |
| Chenodeoxycholic Acid | 0.85 | 1.03 | 0.94 |
| Cimetidine | 1.00 | 0.91 | 0.95 |
| Clemastine Fumarate | 0.86 | 0.91 | 0.88 |
| Curcumin | 0.78 | 0.85 | 0.82 |
| Daidzein | 0.96 | 1.02 | 0.99 |
| Oxibendazole | 1.01 | 0.95 | 0.98 |
| Penicillamine (Cuprimine) | 0.92 | 0.98 | 0.95 |
| Bifonazole | 0.84 | 0.99 | 0.92 |
| Metoprolol Tartrate | 1.00 | 0.96 | 0.98 |
| Etidronate (Didronel) | 0.84 | 0.98 | 0.91 |
| Diethylstilbestrol | 0.88 | 0.99 | 0.93 |
| Diltiazem HCl | 0.93 | 0.97 | 0.95 |
| Diphenhydramine HCl | 0.92 | 0.91 | 0.91 |
| Dapoxetine HCl | 0.87 | 0.88 | 0.87 |
| Risedronic acid (Actonel) | 1.00 | 0.84 | 0.92 |
| Tranexamic Acid | 1.08 | 0.90 | 0.99 |

| | | | |
|---|---|---|---|
| Valaciclovir HCl | 0.76 | 1.02 | 0.89 |
| Ganciclovir | 0.75 | 0.97 | 0.86 |
| Protionamide | 0.94 | 1.02 | 0.98 |
| Idoxuridine | 0.86 | 0.95 | 0.90 |
| Sparfloxacin | 0.78 | 0.86 | 0.82 |
| Felodipine | 0.78 | 0.90 | 0.84 |
| Deflazacort | 0.70 | 1.01 | 0.85 |
| Nizatidine | 0.90 | 0.94 | 0.92 |
| Carbidopa | 0.80 | 0.88 | 0.84 |
| Valsartan | 0.89 | 0.97 | 0.93 |
| Dipyridamole | 0.65 | 0.66 | 0.66 |
| Hydroxyurea | 0.92 | 1.01 | 0.97 |
| Tropisetron | 0.88 | 0.98 | 0.93 |
| Nicotinamide (Vitamin B3) | 0.89 | 0.93 | 0.91 |
| Talc | 1.04 | 1.00 | 1.02 |
| Gabapentin Hydrochloride | 0.90 | 0.98 | 0.94 |
| Diclofenac Sodium | 0.95 | 0.94 | 0.95 |
| Avobenzone | 0.79 | 0.88 | 0.83 |
| Amlodipine | 0.91 | 1.08 | 1.00 |
| Metronidazole | 0.81 | 0.85 | 0.83 |
| Flutamide | 0.86 | 1.00 | 0.93 |
| Fluvastatin Sodium | 1.07 | 0.87 | 0.97 |
| Disodium Cromoglycate | 0.85 | 0.92 | 0.88 |
| Tropicamide | 0.87 | 0.99 | 0.93 |
| Pregnenolone | 0.81 | 0.80 | 0.81 |
| Sulfamethoxazole | 0.92 | 0.86 | 0.89 |
| Sulfisoxazole | 0.97 | 0.94 | 0.95 |
| Crystal Violet | 1.00 | 1.02 | 1.01 |
| Haloperidol | 0.89 | 0.96 | 0.93 |
| Phenindione | 0.90 | 0.96 | 0.93 |
| Alibendol | 0.85 | 1.08 | 0.96 |
| Irsogladine | 1.02 | 1.20 | 1.11 |
| Nystatin (Fungicidin) | 0.95 | 0.93 | 0.94 |
| Isoniazid | 0.90 | 0.99 | 0.94 |
| Resveratrol | 0.92 | 1.00 | 0.96 |
| Levofloxacin | 0.89 | 0.98 | 0.94 |
| Enalapril Maleate | 0.88 | 1.03 | 0.95 |
| Menadione | 0.76 | 0.80 | 0.78 |
| Metformin hydrochloride (Glucophage) | 0.87 | 1.00 | 0.94 |
| Methoxsalen | 0.87 | 1.02 | 0.95 |
| Miconazole Nitrate | 0.88 | 0.98 | 0.93 |
| Sulfamethizole | 0.94 | 1.04 | 0.99 |
| Tolfenamic Acid | 0.84 | 1.00 | 0.92 |
| Pranoprofen | 0.93 | 1.04 | 0.99 |
| Sulphadimethoxine | 0.93 | 0.84 | 0.89 |
| Rimantadine | 0.76 | 1.02 | 0.89 |
| Primidone | 0.99 | 0.77 | 0.88 |
| Nefiracetam | 0.89 | 1.03 | 0.96 |
| Nicorandil | 1.02 | 1.05 | 1.03 |
| Tamoxifen Citrate | 0.92 | 1.11 | 1.02 |
| Meglumine | 0.94 | 0.98 | 0.96 |
| Aripiprazole | 0.85 | 1.04 | 0.95 |
| Methscopolamine | 0.84 | 1.02 | 0.93 |
| Amiodarone HCl | 0.91 | 1.00 | 0.96 |
| Adenine | 1.00 | 0.85 | 0.92 |
| Adenine sulfate | 0.89 | 1.00 | 0.94 |
| Adenine HCl | 0.90 | 0.98 | 0.94 |
| Ticlopidine HCl | 0.92 | 1.10 | 1.01 |
| ATP (Adenosine-Triphosphate) | 0.88 | 1.05 | 0.97 |
| Meclizine 2HCl | 0.75 | 0.97 | 0.86 |
| Mometasone furoate | 1.06 | 0.93 | 0.99 |
| Propylthiouracil | 0.99 | 1.08 | 1.03 |

| | | | | |
|---|---|---|---|---|
| Lacidipine | 0.87 | 0.97 | 0.92 | |
| Procarbazine HCl (Matulane) | 0.85 | 1.02 | 0.94 | |
| Ondansetron (Zofran) | 0.79 | 1.05 | 0.92 | |
| Liranaftate | 0.87 | 0.99 | 0.93 | |
| D-Cycloserine | 0.96 | 1.07 | 1.02 | |
| Sodium butyrate | 0.91 | 1.11 | 1.01 | |
| Sodium orthovanadate | 0.91 | 0.98 | 0.95 | |
| Elvitegravir (GS-9137, JTK-303) | 1.00 | 1.00 | 1.00 | |
| Maraviroc | 0.96 | 1.05 | 1.01 | |
| Raltegravir (MK-0518) | 0.93 | 1.06 | 1.00 | |
| Sulindac | 0.78 | 0.82 | 0.80 | |
| Taurine | 0.81 | 0.93 | 0.87 | |
| Pramipexole dihydrochloride monohydrate | 0.82 | 0.89 | 0.85 | |
| Suplatast Tosylate | 0.86 | 0.89 | 0.87 | |
| Mirtazapine | 0.88 | 1.03 | 0.95 | |
| Benidipine HCl | 0.90 | 0.99 | 0.95 | |
| Formoterol Hemifumarate | 0.84 | 1.03 | 0.94 | |
| Chlormezanone | 0.95 | 1.01 | 0.98 | |
| Ketotifen Fumarate | 0.88 | 0.98 | 0.93 | |
| Urapidil HCl | 0.95 | 1.03 | 0.99 | |
| Ciprofloxacin (Cipro) | 0.84 | 0.86 | 0.85 | |
| Lopinavir | 0.96 | 0.94 | 0.95 | |
| Uridine | 0.80 | 0.79 | 0.80 | |
| Flunarizine 2HCl | 0.88 | 0.89 | 0.89 | |
| Fenticonazole Nitrate | 0.92 | 0.90 | 0.91 | |
| Rebamipide | 0.82 | 1.06 | 0.94 | |
| Epalrestat | 0.88 | 1.02 | 0.95 | |
| Aspartame | 0.89 | 1.21 | 1.05 | |
| Candesartan Cilexetil | 0.77 | 1.23 | 1.00 | |
| Phentolamine Mesylate | 0.97 | 0.98 | 0.98 | |
| Dyclonine HCl | 0.87 | 0.92 | 0.90 | |
| Memantine HCl | 0.85 | 0.96 | 0.90 | |
| Cyproheptadine HCl | 0.84 | 1.04 | 0.94 | |
| Doxifluridine | 0.91 | 1.02 | 0.96 | |
| Pioglitazone HCl | 0.81 | 0.93 | 0.87 | |
| Lornoxicam | 0.92 | 1.07 | 1.00 | |
| Clindamycin phosphate | 0.84 | 1.13 | 0.98 | |
| Alfuzosin HCl | 0.89 | 1.18 | 1.03 | |
| Captopril | 0.90 | 0.95 | 0.92 | |
| Oxytetracycline dihydrate | 0.81 | 1.01 | 0.91 | |
| Orphenadrine Citrate | 0.87 | 0.99 | 0.93 | |
| Gimeracil | 0.79 | 0.99 | 0.89 | |
| Cyclophosphamide monohydrate | 0.83 | 1.04 | 0.94 | |
| Tolnaftate | 1.01 | 1.06 | 1.04 | |
| Terazosin HCl | 0.96 | 1.08 | 1.02 | |
| Bromhexine HCl | 0.82 | 0.98 | 0.90 | |
| Lovastatin | 0.88 | 0.95 | 0.91 | |
| Tiopronin | 0.98 | 0.97 | 0.97 | |
| Balofloxacin | 0.88 | 1.06 | 0.97 | |
| Lafutidine | 1.12 | 1.18 | 1.15 | |
| Ozagrel HCl | 1.03 | 1.06 | 1.05 | |
| Argatroban | 1.04 | 0.87 | 0.96 | |
| Mitiglinide calcium | 1.03 | 1.05 | 1.04 | |
| Mecarbinate | 0.96 | 0.94 | 0.95 | |
| Rosiglitazone HCl | 1.03 | 0.99 | 1.01 | |
| Lisinopril (Zestril) | 1.05 | 0.94 | 0.99 | |
| Atorvastatin Calcium | 1.39 | 1.16 | 1.28 | toxic |
| Famotidine | 1.05 | 1.01 | 1.03 | |
| Moexipril HCl | 1.15 | 0.95 | 1.05 | |
| Clevidipine Butyrate | 1.08 | 1.04 | 1.06 | |
| Adiphenine HCl | 1.02 | 0.86 | 0.94 | |
| Duloxetine HCl | 1.09 | 0.96 | 1.02 | |

| | | | |
|---|---|---|---|
| Trimebutine | 1.03 | 1.02 | 1.02 |
| Ivabradine HCl | 0.96 | 0.96 | 0.96 |
| Rivastigmine Tartrate | 1.07 | 0.95 | 1.01 |
| Topiramate | 1.11 | 1.06 | 1.09 |
| Dexmedetomidine HCl (Precedex) | 1.06 | 0.98 | 1.02 |
| Betaxolol | 1.14 | 0.91 | 1.03 |
| Detomidine HCl | 0.93 | 0.84 | 0.89 |
| Fosinopril sodium (Monopril) | 1.10 | 1.04 | 1.07 |
| Ambrisentan | 1.10 | 0.94 | 1.02 |
| Bexarotene | 0.79 | 0.72 | 0.76 |
| Temocapril HCl | 1.11 | 0.99 | 1.05 |
| Gabexate Mesylate | 0.99 | 0.89 | 0.94 |
| Rasagiline Mesylate | 0.97 | 1.10 | 1.04 |
| Naltrexone HCl | 1.15 | 0.99 | 1.07 |
| Levosulpiride | 1.04 | 0.93 | 0.99 |
| Azasetron HCl (Y-25130) | 1.01 | 0.97 | 0.99 |
| Mizolastine (Mizollen) | 0.94 | 0.98 | 0.96 |
| Flunixin Meglumin | 1.02 | 0.95 | 0.99 |
| Vinpocetine (Cavinton) | 0.98 | 0.90 | 0.94 |
| Lapatinib | 1.00 | 0.95 | 0.98 |
| Blonanserin (Lonasen) | 1.16 | 1.00 | 1.08 |
| Cisatracurium Besylate | 1.09 | 1.01 | 1.05 |
| Dronedarone HCl | 0.91 | 0.96 | 0.94 |
| Conivaptan HCl | 1.15 | 0.96 | 1.06 |
| Ibutilide Fumarate | 0.93 | 1.04 | 0.98 |
| Probucol | 1.16 | 0.97 | 1.07 |
| Arbidol HCl | 1.09 | 1.04 | 1.07 |
| Licofelone | 0.90 | 0.97 | 0.93 |
| Xylose | 1.13 | 0.99 | 1.06 |
| Mestranol | 1.00 | 0.87 | 0.93 |
| Naftopidil | 1.06 | 0.88 | 0.97 |
| S- (+)-Rolipram | 1.09 | 0.92 | 1.01 |
| Bazedoxifene HCl | 1.06 | 0.92 | 0.99 |
| Fudosteine | 0.99 | 1.04 | 1.02 |
| Atropine | 0.86 | 1.07 | 0.97 |
| Roflumilast | 1.07 | 1.16 | 1.12 |
| Gabapentin (Neurontin) | 1.02 | 1.01 | 1.02 |
| Sitafloxacin Hydrate | 1.06 | 0.97 | 1.01 |
| Tebipenem Pivoxil | 1.07 | 0.97 | 1.02 |
| Rosuvastatin Calcium | 1.01 | 1.02 | 1.02 |
| Dichlorphenamide | 1.05 | 0.92 | 0.98 |
| BIBR 953 (Dabigatran etexilate, Pradaxa) | 1.03 | 0.97 | 1.00 |
| Aliskiren Hemifumarate | 0.97 | 1.02 | 1.00 |
| OSI-420 | 1.21 | 1.18 | 1.20 |
| DAPT (GSI-IX) | 1.03 | 0.93 | 0.98 |
| Irinotecan HCl Trihydrate | 1.06 | 1.05 | 1.06 |
| Betamethasone | 0.83 | 0.90 | 0.87 |
| TAME | 1.10 | 0.92 | 1.01 |
| Esomeprazole Sodium | 1.05 | 0.98 | 1.02 |
| Fesoterodine Fumarate | 0.97 | 0.97 | 0.97 |
| Abiraterone Acetate | 1.00 | 0.86 | 0.93 |
| Artemether | 1.02 | 0.85 | 0.94 |
| DL-Carnitine HCl | 0.97 | 0.92 | 0.94 |
| Nalidixic acid | 1.01 | 0.94 | 0.97 |
| Ammonium Glycyrrhizinate | 1.08 | 1.00 | 1.04 |
| D-Mannitol | 1.12 | 1.06 | 1.09 |
| L-carnitine | 1.09 | 1.05 | 1.07 |
| Sorbitol | 1.09 | 0.98 | 1.04 |
| 10-Deacetylbaccatin-III | 1.10 | 0.98 | 1.04 |
| Paeoniflorin | 1.02 | 0.99 | 1.00 |
| Geniposide | 1.06 | 0.41 | 0.73 |
| Genipin | 1.01 | 1.01 | 1.01 |

| | | | |
|---|---|---|---|
| Geniposidic acid | 1.01 | 1.24 | 1.12 |
| Tolbutamide | 1.03 | 1.00 | 1.02 |
| Moxifloxacin HCl | 0.99 | 1.07 | 1.03 |
| Amantadine HCl | 1.11 | 0.94 | 1.02 |
| Amfebutamone HCl | 1.03 | 0.90 | 0.97 |
| Benserazide HCl | 1.12 | 1.00 | 1.06 |
| Bethanechol chloride | 0.94 | 0.93 | 0.93 |
| Chlorpromazine HCl | 1.00 | 0.91 | 0.95 |
| Clindamycin HCl | 1.08 | 1.04 | 1.06 |
| Clonidine HCl | 0.94 | 1.01 | 0.97 |
| Clozapine | 1.03 | 0.99 | 1.01 |
| Pramipexole | 1.05 | 0.91 | 0.98 |
| Domperidone | 1.03 | 1.01 | 1.02 |
| Donepezil HCl (Aricept) | 1.04 | 0.98 | 1.01 |
| Estriol | 1.10 | 0.92 | 1.01 |
| Famciclovir | 0.97 | 1.00 | 0.99 |
| Fleroxacin (Quinodis) | 1.07 | 0.98 | 1.03 |
| Fluocinolone Acetonide | 1.03 | 1.15 | 1.09 |
| Gallamine Triethiodide | 1.09 | 0.96 | 1.02 |
| Imatinib (STI571) | 1.01 | 0.84 | 0.92 |
| Itraconazole | 0.95 | 0.99 | 0.97 |
| Lincomycin HCl | 1.00 | 1.03 | 1.02 |
| Loperamide HCl | 0.90 | 1.00 | 0.95 |
| Manidipine | 1.18 | 1.01 | 1.10 |
| Cidofovir (Vistide) | 1.12 | 0.99 | 1.05 |
| Mitoxantrone HCl | 1.14 | 1.02 | 1.08 |
| Mycophenolic acid | 1.01 | 1.01 | 1.01 |
| Nateglinide | 1.01 | 1.03 | 1.02 |
| Neostigmine bromide (Prostigmin) | 1.10 | 0.94 | 1.02 |
| Nitrendipine | 1.04 | 1.04 | 1.04 |
| Novobiocin Sodium | 0.98 | 0.85 | 0.92 |
| Olanzapine | 0.89 | 0.91 | 0.90 |
| Olopatadine HCl | 0.96 | 0.98 | 0.97 |
| Oxymetazoline HCl | 1.05 | 0.98 | 1.02 |
| Ozagrel | 1.00 | 0.92 | 0.96 |
| Pancuronium dibromide | 1.08 | 0.95 | 1.02 |
| Phenoxybenzamine HCl | 1.08 | 0.87 | 0.98 |
| Propafenone HCl | 0.97 | 0.95 | 0.96 |
| Quinine HCl Dihydrate | 1.05 | 0.92 | 0.99 |
| Racecadotril | 0.87 | 1.11 | 0.99 |
| Ribavirin | 1.00 | 1.04 | 1.02 |
| Rosiglitazone maleate | 0.92 | 0.93 | 0.92 |
| Roxithromycin | 0.88 | 0.94 | 0.91 |
| Salbutamol sulfate (Albuterol) | 0.98 | 1.00 | 0.99 |
| Scopolamine HBr | 1.05 | 0.95 | 1.00 |
| Sotalol | 1.01 | 0.83 | 0.92 |
| Sulfadoxine (Sulphadoxine) | 0.96 | 0.99 | 0.98 |
| Tenoxicam | 0.97 | 0.91 | 0.94 |
| Tobramycin | 0.93 | 0.91 | 0.92 |
| Vardenafil HCl Trihydrate | 1.07 | 0.88 | 0.98 |
| Xylazine HCl | 0.82 | 0.99 | 0.90 |
| Maprotiline HCl | 0.93 | 0.98 | 0.96 |
| Naphazoline HCl | 1.05 | 0.99 | 1.02 |
| Epinephrine Bitartrate | 1.02 | 0.81 | 0.91 |
| L-Adrenaline | 0.84 | 0.95 | 0.89 |
| DL-Adrenaline | 0.95 | 1.00 | 0.98 |
| Phenytoin | 1.01 | 0.91 | 0.96 |
| Methacycline HCl | 0.93 | 0.84 | 0.88 |
| Ciclopirox | 0.93 | 0.86 | 0.90 |
| Dopamine HCl | 0.93 | 0.75 | 0.84 |
| Ritodrine HCl | 0.92 | 0.95 | 0.93 |
| Isoconazole nitrate | 1.04 | 0.98 | 1.01 |

| | | | |
|---|---|---|---|
| Econazole nitrate | 1.02 | 0.85 | 0.94 |
| Miconazole | 0.94 | 0.92 | 0.93 |
| Secnidazole | 1.15 | 0.91 | 1.03 |
| Acetanilide | 0.94 | 0.95 | 0.94 |
| Riboflavin (Vitamin B2) | 0.79 | 0.73 | 0.76 |
| Clomipramine HCl | 1.00 | 0.85 | 0.93 |
| Phenformin HCl | 0.99 | 0.88 | 0.94 |
| Ceftiofur HCl | 1.15 | 0.98 | 1.07 |
| Cefprozil hydrate (Cefzil) | 0.92 | 0.90 | 0.91 |
| Scopine | 1.00 | 0.96 | 0.98 |
| Tiotropium Bromide hydrate | 1.13 | 0.85 | 0.99 |
| Trospium chloride | 0.96 | 0.92 | 0.94 |
| Tolterodine tartrate | 1.13 | 0.89 | 1.01 |
| Sulbactam sodium | 1.06 | 1.00 | 1.03 |
| 5-Aminolevulinic acid HCl | 1.02 | 0.99 | 1.00 |
| Clarithromycin | 1.04 | 0.91 | 0.97 |
| Rosiglitazone | 0.80 | 0.78 | 0.79 |
| Terbinafine HCl | 1.08 | 0.99 | 1.03 |
| Cortisone acetate | 0.97 | 1.07 | 1.02 |
| Amiloride hydrochloride dihydrate | 1.04 | 0.85 | 0.94 |
| Clomifene citrate | 1.08 | 0.90 | 0.99 |
| Hydralazine HCl | 0.90 | 0.91 | 0.91 |
| Oxacillin sodium monohydrate | 0.91 | 0.99 | 0.95 |
| Cloxacillin Sodium | 0.96 | 1.02 | 0.99 |
| Isoprenaline HCl | 0.91 | 0.87 | 0.89 |
| Medroxyprogesterone acetate | 0.92 | 0.85 | 0.88 |
| Neomycin sulfate | 1.00 | 1.01 | 1.01 |
| Phenylephrine HCl | 0.88 | 0.92 | 0.90 |
| Prednisolone acetate (Omnipred) | 1.11 | 0.87 | 0.99 |
| Streptomycin sulfate | 0.85 | 0.84 | 0.85 |
| Tetracaine HCl | 0.95 | 0.90 | 0.92 |
| Tetracycline HCl | 0.93 | 0.83 | 0.88 |
| Vancomycin HCl (Vancocin) | 0.97 | 0.87 | 0.92 |
| Xylometazoline HCl | 0.98 | 0.86 | 0.92 |
| Zidovudine | 0.95 | 0.82 | 0.88 |
| Quinapril HCl | 0.97 | 0.81 | 0.89 |
| Trazodone HCl | 0.95 | 0.84 | 0.90 |
| Thiamphenicol | 0.92 | 0.77 | 0.85 |
| Clobetasol propionate | 0.99 | 0.76 | 0.88 |
| Brompheniramine hydrogen maleate | 0.99 | 0.78 | 0.89 |
| Dimethyl Fumarate | 1.23 | 0.87 | 1.05 |
| Calcium levofolinate (Calcium Folinate) | 1.26 | 1.25 | 1.26 |
| Miglitol | 1.17 | 0.89 | 1.03 |
| Pioglitazone | 1.12 | 0.87 | 1.00 |
| Pramiracetam | 0.98 | 0.79 | 0.88 |
| Clindamycin palmitate HCl | 0.99 | 0.84 | 0.91 |
| Oseltamivir phosphate | 0.90 | 0.82 | 0.86 |
| L-Thyroxine | 1.05 | 0.85 | 0.95 |
| Gliclazide | 0.91 | 0.78 | 0.84 |
| Acemetacin | 0.91 | 0.79 | 0.85 |
| Tioxolone | 1.00 | 0.77 | 0.89 |
| Dehydroepiandrosterone (DHEA) | 1.00 | 0.85 | 0.92 |
| Idebenone | 0.80 | 0.97 | 0.88 |
| Mifepristone | 0.99 | 0.88 | 0.94 |
| Fluocinonide | 0.87 | 0.80 | 0.84 |
| Lonidamine | 0.90 | 0.97 | 0.93 |
| Ethisterone | 0.95 | 0.76 | 0.85 |
| Clorsulon | 0.86 | 0.95 | 0.91 |
| Arecoline | 0.87 | 0.95 | 0.91 |
| Noradrenaline bitartrate monohydrate | 0.90 | 0.83 | 0.86 |
| Ibrutinib (PCI-32765) | 1.38 | 1.10 | 1.24 |
| Peramivir Trihydrate | 0.85 | 0.85 | 0.85 |

| | | | | |
|---|---|---|---|---|
| Methimazole | 1.07 | 0.78 | 0.92 | |
| Dabrafenib (GSK2118436) | 0.97 | 0.80 | 0.89 | |
| Carfilzomib (PR-171) | 0.60 | 0.48 | 0.54 | toxic |
| Cobicistat (GS-9350) | 0.90 | 0.88 | 0.89 | |
| Hygromycin B | 0.87 | 0.58 | 0.73 | |
| Carbazochrome sodium sulfonate (AC-17) | 0.95 | 0.73 | 0.84 | |
| Rivaroxaban | 1.04 | 0.93 | 0.99 | |
| Paroxetine HCl | 0.79 | 0.92 | 0.86 | |
| Zanamivir | 0.91 | 0.85 | 0.88 | |
| Zaltoprofen | 1.02 | 0.85 | 0.93 | |
| Pazopanib | 0.85 | 0.89 | 0.87 | |
| Amoxicillin | 1.20 | 0.99 | 1.09 | |
| Niflumic acid | 0.88 | 0.84 | 0.86 | |
| Ciclopirox ethanolamine | 0.92 | 0.76 | 0.84 | |
| Rimonabant | 0.85 | 0.92 | 0.88 | |
| Cabazitaxel | 1.16 | 0.98 | 1.07 | |
| Bufexamac | 0.98 | 0.86 | 0.92 | |
| Lamotrigine | 0.89 | 0.95 | 0.92 | |
| PMSF | 0.87 | 0.92 | 0.90 | |
| Fenoprofen calcium hydrate | 1.01 | 0.92 | 0.97 | |
| Niclosamide (Niclocide) | 0.70 | 0.52 | 0.61 | toxic |
| Linagliptin | 0.97 | 0.94 | 0.95 | |
| Sulfasalazine | 1.02 | 0.91 | 0.96 | |
| Daunorubicin HCl | 1.02 | 0.96 | 0.99 | |
| Pravastatin sodium | 0.96 | 0.98 | 0.97 | |
| Bepotastine Besilate | 0.82 | 0.86 | 0.84 | |
| Fosaprepitant dimeglumine salt | 0.80 | 0.83 | 0.81 | |
| Droxidopa (L-DOPS) | 0.90 | 0.82 | 0.86 | |
| Rofecoxib | 0.98 | 0.88 | 0.93 | |
| Lurasidone HCl | 1.05 | 0.78 | 0.92 | |
| Cinepazide maleate | 1.11 | 0.89 | 1.00 | |
| Azilsartan | 1.01 | 0.91 | 0.96 | |
| Solifenacin succinate | 0.88 | 0.88 | 0.88 | |
| Palonosetron HCl | 0.93 | 0.88 | 0.91 | |
| Azelnidipine | 1.12 | 0.97 | 1.04 | |
| Alverine Citrate | 1.01 | 0.95 | 0.98 | |
| Besifloxacin HCl (Besivance) | 0.89 | 0.84 | 0.86 | |
| Azilsartan Medoxomil | 0.88 | 0.87 | 0.87 | |
| Danofloxacin Mesylate | 0.83 | 0.92 | 0.87 | |
| Enrofloxacin | 0.96 | 0.94 | 0.95 | |
| Medetomidine HCl | 1.03 | 0.88 | 0.95 | |
| Diclofenac Potassium | 1.27 | 0.97 | 1.12 | |
| Amikacin sulfate | 0.92 | 0.94 | 0.93 | |
| Naloxone HCl | 0.97 | 0.88 | 0.93 | |
| (R)-baclofen | 0.98 | 1.01 | 1.00 | |
| Caspofungin Acetate | 1.00 | 0.92 | 0.96 | |
| Dexmedetomidine | 0.97 | 1.05 | 1.01 | |
| Beclomethasone dipropionate | 1.26 | 1.24 | 1.25 | |
| Atovaquone | 0.84 | 0.92 | 0.88 | |
| Acetylcysteine | 0.97 | 0.92 | 0.95 | |
| Ulipristal | 1.06 | 1.03 | 1.04 | |
| Indacaterol Maleate | 1.10 | 0.81 | 0.95 | |
| Creatinine | 1.00 | 0.93 | 0.96 | |
| Moguisteine | 1.02 | 0.89 | 0.95 | |
| Nadifloxacin | 1.01 | 0.83 | 0.92 | |
| Pidotimod | 0.81 | 1.06 | 0.93 | |
| Pyridoxine HCl | 0.85 | 0.92 | 0.88 | |
| Vitamin C | 0.91 | 0.83 | 0.87 | |
| Sulfathiazole | 1.00 | 0.82 | 0.91 | |
| Oxybutynin chloride | 0.85 | 0.97 | 0.91 | |
| Ornidazole | 1.00 | 0.94 | 0.97 | |
| Docosanol (Abreva) | 0.96 | 0.99 | 0.98 | |

| | | | | |
|---|---|---|---|---|
| Trimethoprim | 1.00 | 0.80 | 0.90 | |
| Biotin (Vitamin B7) | 0.91 | 0.95 | 0.93 | |
| Sulfamerazine | 1.01 | 0.97 | 0.99 | |
| Sulfamethazine | 1.06 | 0.86 | 0.96 | |
| Sodium salicylate | 0.97 | 1.01 | 0.99 | |
| Methylthiouracil | 0.89 | 0.94 | 0.91 | |
| Methenamine | 0.91 | 0.90 | 0.91 | |
| Milnacipran HCl | 0.92 | 0.94 | 0.93 | |
| Darifenacin HBr | 0.99 | 0.91 | 0.95 | |
| Tripelennamine HCl | 1.06 | 0.89 | 0.98 | |
| Ibandronate sodium | 0.89 | 0.82 | 0.85 | |
| Estradiol valerate | 0.98 | 1.03 | 1.01 | |
| Articaine HCl | 1.02 | 0.84 | 0.93 | |
| Gliquidone | 0.99 | 0.83 | 0.91 | |
| Butenafine HCl | 0.97 | 0.90 | 0.93 | |
| Mepivacaine HCl | 0.81 | 0.91 | 0.86 | |
| Naftifine HCl | 0.83 | 0.85 | 0.84 | |
| Ethynodiol diacetate | 0.94 | 0.99 | 0.97 | |
| Sertaconazole nitrate | 1.00 | 0.90 | 0.95 | |
| Tylosin tartrate | 0.97 | 0.90 | 0.94 | |
| Abacavir sulfate | 0.84 | 0.92 | 0.88 | |
| Altrenogest | 0.93 | 0.88 | 0.90 | |
| Ampicillin sodium | 0.84 | 0.87 | 0.86 | |
| Anagrelide HCl | 1.02 | 0.85 | 0.94 | |
| Antipyrine | 0.81 | 0.98 | 0.90 | |
| L-Arginine HCl | 0.88 | 0.94 | 0.91 | |
| Atomoxetine HCl | 0.92 | 0.87 | 0.89 | |
| Betahistine 2HCl | 0.86 | 0.88 | 0.87 | |
| Brinzolamide | 0.88 | 0.91 | 0.90 | |
| Carbenicillin disodium | 1.05 | 0.85 | 0.95 | |
| Amitriptyline HCl | 1.09 | 0.91 | 1.00 | |
| Adrenalone HCl | 0.97 | 0.95 | 0.96 | |
| Azatadine dimaleate | 0.99 | 0.79 | 0.89 | |
| (+,-)-Octopamine HCl | 0.79 | 0.91 | 0.85 | |
| Ropinirole HCl | 0.79 | 0.80 | 0.79 | |
| Azlocillin sodium salt | 0.96 | 0.93 | 0.94 | |
| Azacyclonol | 0.88 | 0.91 | 0.89 | |
| Reboxetine mesylate | 1.00 | 0.84 | 0.92 | |
| Triflusal | 0.94 | 0.88 | 0.91 | |
| Trifluoperazine 2HCl | 0.76 | 0.83 | 0.79 | |
| Meptazinol HCl | 0.89 | 0.89 | 0.89 | |
| Fexofenadine HCl | 1.07 | 1.06 | 1.07 | |
| Amidopyrine | 1.11 | 0.84 | 0.97 | |
| Thiamine HCl (Vitamin B1) | 1.01 | 0.79 | 0.90 | |
| Moclobemide (Ro 111163) | 0.93 | 0.87 | 0.90 | |
| Pergolide mesylate | 1.18 | 0.97 | 1.07 | |
| Lithocholic acid | 1.02 | 0.94 | 0.98 | |
| Ethambutol HCl | 0.92 | 0.87 | 0.89 | |
| Doxycycline HCl | 0.90 | 1.05 | 0.98 | |
| Clinafloxacin (PD127391) | 1.03 | 0.86 | 0.95 | |
| Pemirolast (BMY 26517) potassium | 0.93 | 1.04 | 0.98 | |
| Mirabegron | 1.03 | 0.88 | 0.95 | |
| Acebutolol HCl | 0.98 | 0.96 | 0.97 | |
| Ampiroxicam | 0.99 | 0.85 | 0.92 | |
| Desloratadine | 0.92 | 0.90 | 0.91 | |
| Sodium Monofluorophosphate | 0.81 | 0.94 | 0.88 | |
| Hyoscyamine | 0.95 | 0.94 | 0.94 | |
| Cyclamic acid | 0.91 | 0.80 | 0.86 | |
| Ouabain | 0.59 | 0.47 | 0.53 | toxic |
| Allylthiourea | 1.08 | 0.87 | 0.98 | |
| Sodium Picosulfate | 0.96 | 0.96 | 0.96 | |
| Tolcapone | 0.93 | 0.77 | 0.85 | |

| | | | |
|---|---|---|---|
| Probenecid | 0.99 | 0.88 | 0.94 |
| Procaine HCl | 0.89 | 0.98 | 0.93 |
| Homatropine Bromide | 0.96 | 0.96 | 0.96 |
| Hydroxyzine 2HCl | 0.90 | 0.96 | 0.93 |
| Flavoxate HCl | 1.06 | 1.00 | 1.03 |
| Colistin Sulfate | 0.97 | 1.06 | 1.01 |
| Nevirapine | 1.13 | 0.94 | 1.04 |
| Aclidinium Bromide | 1.02 | 1.08 | 1.05 |
| Diphemanil Methylsulfate | 0.94 | 0.98 | 0.96 |
| Vitamin D2 | 0.96 | 0.99 | 0.98 |
| Doxapram HCl | 0.89 | 0.81 | 0.85 |
| Dibucaine HCl | 1.07 | 0.88 | 0.98 |
| Methazolamide | 0.94 | 0.85 | 0.90 |
| Norethindrone | 0.99 | 0.92 | 0.96 |
| Olsalazine Sodium | 1.05 | 0.96 | 1.01 |
| Nafcillin Sodium | 1.04 | 0.97 | 1.00 |
| Tetrahydrozoline HCl | 1.06 | 0.95 | 1.00 |
| Toltrazuril | 1.03 | 0.94 | 0.99 |
| Bisacodyl | 0.99 | 1.09 | 1.04 |
| Carbimazole | 1.05 | 0.90 | 0.98 |
| Valdecoxib | 1.02 | 0.98 | 1.00 |
| Valganciclovir HCl | 0.85 | 0.99 | 0.92 |
| Nabumetone | 0.96 | 0.89 | 0.92 |
| Netilmicin Sulfate | 0.99 | 0.77 | 0.88 |
| Sertraline HCl | 1.04 | 0.95 | 0.99 |
| Spironolactone | 0.98 | 0.98 | 0.98 |
| Retapamulin | 1.01 | 0.95 | 0.98 |
| Methyclothiazide | 0.96 | 1.07 | 1.01 |
| Cytarabine | 1.01 | 0.89 | 0.95 |
| Erythromycin Ethylsuccinate | 0.89 | 0.85 | 0.87 |
| Ronidazole | 0.98 | 0.97 | 0.98 |
| Vitamin D3 | 0.84 | 0.89 | 0.87 |
| Glipizide | 0.96 | 0.83 | 0.89 |
| Guanabenz Acetate | 0.91 | 0.97 | 0.94 |
| Dequalinium chloride | 0.81 | 0.76 | 0.78 |
| Deferiprone | 1.04 | 0.97 | 1.01 |
| Tinidazole | 1.07 | 1.00 | 1.04 |
| Hexamethonium bromide | 0.99 | 0.93 | 0.96 |
| Decamethonium Bromide | 0.93 | 0.95 | 0.94 |
| Sodium 4-Aminosalicylate | 1.02 | 0.97 | 1.00 |
| Flucytosine | 0.89 | 1.02 | 0.95 |
| Ipratropium Bromide | 0.82 | 0.88 | 0.85 |
| Propranolol HCl | 0.90 | 0.84 | 0.87 |
| Mequinol | 1.04 | 0.96 | 1.00 |
| Mefenamic Acid | 0.75 | 0.93 | 0.84 |
| Ticagrelor | 0.95 | 1.01 | 0.98 |
| Triamterene | 0.83 | 0.99 | 0.91 |
| Sulfacetamide Sodium | 1.01 | 0.89 | 0.95 |
| Lomerizine HCl | 1.00 | 0.95 | 0.98 |
| Levobetaxolol HCl | 0.82 | 0.98 | 0.90 |
| Loxapine Succinate | 0.93 | 0.85 | 0.89 |
| Oxymetholone | 0.88 | 0.95 | 0.92 |
| Flumethasone | 0.79 | 0.79 | 0.79 |
| Halobetasol Propionate | 0.83 | 0.66 | 0.74 |
| Fenspiride HCl | 1.03 | 1.10 | 1.06 |
| Indapamide | 1.07 | 1.01 | 1.04 |
| Pramoxine HCl | 1.11 | 0.97 | 1.04 |
| Bismuth Subcitrate Potassium | 1.11 | 0.95 | 1.03 |
| Difluprednate | 1.05 | 1.00 | 1.03 |
| Droperidol | 1.07 | 1.00 | 1.03 |
| Dydrogesterone | 1.00 | 0.96 | 0.98 |
| Dexlansoprazole | 0.92 | 0.83 | 0.88 |

| | | | |
|---|---|---|---|
| Esmolol HCl | 0.81 | 0.76 | 0.78 |
| Voglibose | 1.01 | 0.90 | 0.95 |
| Eprosartan Mesylate | 1.00 | 0.98 | 0.99 |
| Diminazene Aceturate | 0.99 | 0.95 | 0.97 |
| Suprofen | 0.91 | 0.89 | 0.90 |
| Ethionamide | 0.84 | 0.94 | 0.89 |
| Clodronate Disodium | 0.96 | 0.97 | 0.97 |
| Dicloxacillin Sodium | 0.92 | 0.92 | 0.92 |
| Triclabendazole | 0.91 | 0.86 | 0.88 |
| Didanosine | 1.07 | 0.93 | 1.00 |
| Isovaleramide | 0.97 | 0.94 | 0.96 |
| Histamine Phosphate | 0.95 | 0.94 | 0.95 |
| Sulconazole Nitrate | 1.10 | 0.92 | 1.01 |
| Tilmicosin | 1.14 | 0.92 | 1.03 |
| Troxipide | 1.07 | 1.00 | 1.03 |
| Ranolazine | 1.02 | 0.87 | 0.95 |
| Fenoprofen Calcium | 0.88 | 1.05 | 0.97 |
| Clorprenaline HCl | 0.90 | 0.94 | 0.92 |
| Carprofen | 0.91 | 0.94 | 0.93 |
| Eprazinone 2HCl | 0.95 | 0.94 | 0.95 |
| Dropropizine | 1.05 | 0.93 | 0.99 |
| Amprolium HCl | 0.92 | 0.91 | 0.91 |
| Decoquinate | 0.98 | 1.08 | 1.03 |
| Bacitracin | 0.96 | 1.05 | 1.00 |
| Azithromycin Dihydrate | 1.04 | 0.93 | 0.99 |
| Ampicillin Trihydrate | 0.86 | 0.97 | 0.92 |
| Orbifloxacin | 0.83 | 1.02 | 0.92 |
| Penfluridol | 0.91 | 0.95 | 0.93 |
| Ethamsylate | 0.85 | 0.82 | 0.84 |
| D-Phenylalanine | 1.05 | 0.92 | 0.98 |
| Flubendazole | 0.74 | 0.95 | 0.84 |
| Chlorzoxazone | 0.73 | 0.94 | 0.83 |
| Chlortetracycline HCl | 0.79 | 0.80 | 0.79 |
| Chloroquine Phosphate | 0.88 | 1.03 | 0.96 |
| Roxatidine Acetate HCl | 0.95 | 0.99 | 0.97 |
| Bezafibrate | 0.98 | 1.09 | 1.04 |
| Benzoic Acid | 0.83 | 1.00 | 0.92 |
| Benzethonium Chloride | 0.81 | 0.87 | 0.84 |
| Doxofylline | 0.99 | 0.93 | 0.96 |
| Benzydamine HCl | 0.80 | 1.01 | 0.91 |
| Chlorpropamide | 0.93 | 0.94 | 0.94 |
| Cyromazine | 0.89 | 0.95 | 0.92 |
| Pefloxacin Mesylate | 0.83 | 0.92 | 0.87 |
| Coumarin | 0.92 | 1.01 | 0.97 |
| Choline Chloride | 0.87 | 0.97 | 0.92 |
| Cetylpyridinium Chloride | 0.99 | 0.94 | 0.96 |
| Sodium Gluconate | 0.85 | 0.85 | 0.85 |
| Sulfaguanidine | 0.89 | 0.93 | 0.91 |
| Trometamol | 0.78 | 0.79 | 0.78 |
| Uracil | 0.82 | 0.97 | 0.89 |
| Climbazole | 0.87 | 1.05 | 0.96 |
| Mezlocillin Sodium | 0.84 | 0.91 | 0.88 |
| Nefopam HCl | 0.87 | 1.02 | 0.95 |
| Nicardipine HCl | 0.95 | 0.87 | 0.91 |
| Nifuroxazide | 0.85 | 0.87 | 0.86 |
| Paromomycin Sulfate | 1.04 | 0.92 | 0.98 |
| Tiratricol | 0.88 | 1.11 | 0.99 |
| Domiphen Bromide | 0.98 | 0.94 | 0.96 |
| Salicylanilide | 0.87 | 0.96 | 0.91 |
| Sasapyrine | 0.87 | 1.08 | 0.97 |
| Cyclandelate | 0.86 | 1.10 | 0.98 |
| Cinchophen | 0.86 | 1.02 | 0.94 |

| | | | | |
|---|---|---|---|---|
| Betamipron | 0.88 | 0.94 | 0.91 | |
| Chlorquinaldol | 0.92 | 0.87 | 0.89 | |
| Azaguanine-8 | 0.90 | 1.00 | 0.95 | |
| Broxyquinoline | 0.90 | 0.85 | 0.88 | |
| Bemegride | 1.06 | 0.85 | 0.95 | |
| Aminothiazole | 0.93 | 0.96 | 0.95 | |
| Antazoline HCl | 0.82 | 1.00 | 0.91 | |
| Tolperisone HCl | 0.90 | 0.92 | 0.91 | |
| Florfenicol | 0.79 | 1.02 | 0.91 | |
| Furaltadone HCl | 0.87 | 0.91 | 0.89 | |
| Isosorbide | 0.93 | 1.01 | 0.97 | |
| Dibenzothiophene | 0.85 | 0.97 | 0.91 | |
| Cysteamine HCl | 0.97 | 0.89 | 0.93 | |
| Clofibric Acid | 0.92 | 1.02 | 0.97 | |
| Chlorocresol | 0.72 | 0.87 | 0.79 | |
| Benzocaine | 0.87 | 1.01 | 0.94 | |
| Montelukast Sodium | 0.86 | 0.93 | 0.90 | |
| Dirithromycin | 1.05 | 0.93 | 0.99 | |
| Sucralose | 1.08 | 0.89 | 0.99 | |
| Valnemulin HCl | 0.98 | 1.01 | 1.00 | |
| Liothyronine Sodium | 0.82 | 0.90 | 0.86 | |
| Amoxapine | 0.97 | 0.91 | 0.94 | |
| Azaperone | 0.95 | 1.06 | 1.00 | |
| Benzbromarone | 0.69 | 0.75 | 0.72 | toxic |
| Mevastatin | 0.94 | 0.95 | 0.95 | |
| Mexiletine HCl | 1.07 | 1.05 | 1.06 | |
| Minocycline HCl | 0.83 | 1.04 | 0.94 | |
| Fidaxomicin | 0.96 | 0.91 | 0.93 | |
| Fluorometholone Acetate | 0.94 | 0.84 | 0.89 | |
| Oxybuprocaine HCl | 0.91 | 0.95 | 0.93 | |
| Oxaprozin | 0.93 | 1.15 | 1.04 | |
| Pilocarpine HCl | 0.93 | 1.02 | 0.98 | |
| Nithiamide | 0.81 | 1.02 | 0.92 | |
| Zoxazolamine | 1.17 | 1.00 | 1.09 | |
| Phenazopyridine HCl | 0.95 | 0.81 | 0.88 | |
| Triamcinolone | 1.05 | 0.92 | 0.99 | |
| Primaquine Diphosphate | 1.17 | 0.84 | 1.01 | |
| Cepharanthine | 1.08 | 1.12 | 1.10 | |
| Bergapten | 1.02 | 1.01 | 1.02 | |
| Doxylamine Succinate | 1.16 | 1.07 | 1.12 | |
| Cetrimonium Bromide (CTAB) | 1.03 | 0.86 | 0.95 | |
| Deoxycorticosterone acetate | 1.16 | 1.02 | 1.09 | |
| Serotonin HCl | 1.15 | 0.96 | 1.05 | |
| Sodium ascorbate | 1.14 | 0.94 | 1.04 | |
| Potassium Iodide | 0.99 | 0.95 | 0.97 | |
| Alexidine HCl | 0.38 | 0.41 | 0.40 | toxic |
| 9-Aminoacridine | 0.89 | 1.08 | 0.98 | |
| Anisindione | 0.93 | 0.97 | 0.95 | |
| Anisotropine Methylbromide | 0.95 | 0.84 | 0.89 | |
| Auranofin | 0.83 | 0.70 | 0.77 | toxic |
| Benzthiazide | 1.20 | 0.90 | 1.05 | |
| Bromocriptine Mesylate | 0.87 | 1.06 | 0.97 | |
| Butacaine | 1.28 | 1.26 | 1.27 | |
| Calcium Gluceptate | 1.22 | 1.00 | 1.11 | |
| Ceftazidime Pentahydrate | 1.14 | 0.88 | 1.01 | |
| Clinafloxacin HCl | 1.28 | 0.89 | 1.09 | |
| Clopamide | 1.08 | 0.96 | 1.02 | |
| Clorgyline HCl | 1.12 | 0.96 | 1.04 | |
| Colistimethate Sodium | 0.93 | 0.91 | 0.92 | |
| Dibenzepine HCl | 1.09 | 0.98 | 1.04 | |
| Dimaprit 2HCl | 0.99 | 0.89 | 0.94 | |
| Diphenylpyraline HCl | 1.11 | 0.90 | 1.01 | |

| | | | | |
|---|---|---|---|---|
| Disopyramide Phosphate | 1.14 | 0.94 | 1.04 | |
| Emetine | 0.51 | 0.53 | 0.52 | toxic |
| Ethoxzolamide | 1.13 | 1.00 | 1.06 | |
| Famprofazone | 1.08 | 0.92 | 1.00 | |
| Guanethidine Sulfate | 1.06 | 0.89 | 0.98 | |
| Hemicholinium Bromide | 1.13 | 0.90 | 1.02 | |
| Isoetharine Mesylate | 0.78 | 0.77 | 0.77 | |
| Meclocycline Sulfosalicylate | 1.02 | 0.96 | 0.99 | |
| Medrysone | 1.24 | 1.03 | 1.14 | |
| Mepiroxol | 0.91 | 0.95 | 0.93 | |
| Mesoridazine Besylate | 1.02 | 1.02 | 1.02 | |
| Metaproterenol Sulfate | 1.36 | 1.15 | 1.25 | toxic |
| Methapyrilene HCl | 1.01 | 1.00 | 1.01 | |
| Methoxamine HCl | 1.09 | 0.96 | 1.03 | |
| Meticrane | 1.04 | 0.92 | 0.98 | |
| Moxalactam Disodium | 1.05 | 0.91 | 0.98 | |
| Nalmefene HCl | 1.05 | 0.97 | 1.01 | |
| Nialamide | 1.24 | 1.00 | 1.12 | |
| Oxethazaine | 1.06 | 1.00 | 1.03 | |
| Oxprenolol HCl | 1.10 | 1.02 | 1.06 | |
| Pentoxifylline | 1.11 | 1.09 | 1.10 | |
| Physostigmine Salicylate | 1.09 | 0.94 | 1.02 | |
| Piromidic Acid | 1.04 | 0.97 | 1.01 | |
| Procyclidine HCl | 1.22 | 0.91 | 1.07 | |
| R-(-)-Apomorphine HCl Hemihydrate | 0.75 | 0.69 | 0.72 | |
| Ractopamine HCl | 1.30 | 1.27 | 1.28 | |
| Rolitetracycline | 1.00 | 0.91 | 0.95 | |
| Terfenadine | 1.13 | 1.07 | 1.10 | |
| Thiostrepton | 1.05 | 0.94 | 1.00 | |
| Thonzonium Bromide | 1.32 | 1.20 | 1.26 | |
| Tolazamide | 1.22 | 0.99 | 1.11 | |
| Tacrine HCl | 1.19 | 1.05 | 1.12 | |
| Carbachol | 1.06 | 0.96 | 1.01 | |
| Tolmetin Sodium | 1.04 | 0.95 | 0.99 | |
| Cinoxacin | 1.02 | 0.98 | 1.00 | |
| Glafenine HCl | 0.93 | 1.03 | 0.98 | |
| Noscapine HCl | 1.04 | 1.05 | 1.04 | |
| Phenothrin | 1.20 | 0.88 | 1.04 | |
| Phthalylsulfacetamide | 1.25 | 1.04 | 1.14 | |
| Pinacidil | 1.19 | 1.05 | 1.12 | |
| Suxibuzone | 1.05 | 1.02 | 1.04 | |
| Carbenoxolone Sodium | 0.93 | 1.06 | 1.00 | |
| Tioconazole | 1.07 | 0.92 | 0.99 | |
| Pridinol Methanesulfonate | 1.12 | 0.94 | 1.03 | |
| Triflupromazine HCl | 1.11 | 0.95 | 1.03 | |
| Dicyclomine HCl | 1.04 | 1.00 | 1.02 | |
| Thioridazine HCl | 1.29 | 1.28 | 1.28 | |
| Mepenzolate Bromide | 0.93 | 0.96 | 0.95 | |
| Aceclidine HCl | 1.06 | 0.96 | 1.01 | |
| Imipramine HCl | 1.12 | 0.95 | 1.03 | |
| Methylhydantoin-5-(D) | 1.36 | 1.22 | 1.29 | |
| Metrizamide | 1.28 | 1.28 | 1.28 | |
| Proadifen HCl | 1.18 | 0.98 | 1.08 | |
| Pyrilamine Maleate | 0.92 | 0.98 | 0.95 | |
| Bekanamycin | 1.09 | 0.95 | 1.02 | |
| Difloxacin HCl | 1.00 | 0.99 | 0.99 | |
| Fosfomycin Tromethamine | 1.08 | 1.01 | 1.05 | |
| Azelastine HCl | 1.11 | 1.12 | 1.12 | |
| Bendroflumethiazide | 1.15 | 0.94 | 1.05 | |
| Bentiromide | 1.04 | 1.08 | 1.06 | |
| Bephenium Hydroxynaphthoate | 0.96 | 0.95 | 0.95 | |
| Guanidine HCl | 1.00 | 0.98 | 0.99 | |

| | | | |
|---|---|---|---|
| Potassium Canrenoate | 0.93 | 0.94 | 0.94 |
| Cephapirin Sodium | 1.07 | 0.95 | 1.01 |
| Clofoctol | 0.93 | 1.06 | 1.00 |
| Dichlorisone Acetate | 1.13 | 1.06 | 1.09 |
| Digoxigenin | 1.12 | 0.92 | 1.02 |
| Diperodon HCl | 1.12 | 1.03 | 1.08 |
| Isoxicam | 0.91 | 1.03 | 0.97 |
| Lithium Citrate | 0.88 | 0.97 | 0.93 |
| Lofexidine HCl | 1.03 | 1.02 | 1.03 |
| Nifenazone | 0.92 | 0.94 | 0.93 |
| Sulbactam | 0.96 | 1.01 | 0.99 |
| Pasiniazid | 0.98 | 1.06 | 1.02 |
| Picrotoxinin | 1.03 | 0.98 | 1.00 |
| Pindolol | 1.03 | 0.97 | 1.00 |
| Prochlorperazine Dimaleate | 1.13 | 0.96 | 1.04 |
| Procodazole | 0.88 | 0.97 | 0.93 |
| Quipazine Maleate | 1.09 | 1.10 | 1.09 |
| Diclofenac Diethylamine | 1.25 | 1.05 | 1.15 |
| Phosphatidylcholine | 0.97 | 1.09 | 1.03 |
| Spiramycin | 1.27 | 1.27 | 1.27 |
| Sodium 4-aminohippurate Hydrate | 1.02 | 0.94 | 0.98 |
| Misoprostol | 0.70 | 0.85 | 0.78 |
| Trimipramine Maleate | 0.81 | 0.95 | 0.88 |
| Eltrombopag | 1.17 | 0.97 | 1.07 |
| Tacrolimus (FK506) | 0.97 | 1.01 | 0.99 |
| Pimecrolimus | 1.00 | 1.06 | 1.03 |
| Plerixafor (AMD3100) | 0.81 | 0.95 | 0.88 |
| 2-Thiouracil | 0.87 | 1.09 | 0.98 |
| Pentamidine | 0.90 | 1.05 | 0.97 |
| Clofazimine | 1.01 | 1.18 | 1.09 |
| Sarafloxacin HCl | 1.06 | 1.03 | 1.04 |
| Nimesulide | 1.08 | 0.98 | 1.03 |
| Almotriptan Malate | 0.84 | 1.00 | 0.92 |
| Cephalomannine | 1.13 | 1.06 | 1.09 |
| D-Pantothenic acid | 1.06 | 0.98 | 1.02 |
| Amoxicillin Sodium | 1.12 | 1.03 | 1.08 |
| Clindamycin | 1.00 | 1.10 | 1.05 |
| Dexamethasone acetate | 1.01 | 0.97 | 0.99 |
| Avanafil | 1.07 | 1.09 | 1.08 |
| Deoxyarbutin | 0.95 | 0.99 | 0.97 |
| Fluticasone propionate | 0.89 | 0.95 | 0.92 |
| Cytidine | 1.08 | 0.96 | 1.02 |
| Imidapril HCl | 1.02 | 0.89 | 0.95 |
| Bupivacaine HCl | 1.00 | 0.92 | 0.96 |
| Spectinomycin HCl | 0.99 | 1.02 | 1.00 |
| Phenacetin | 0.95 | 1.06 | 1.01 |
| Aspirin | 0.93 | 1.03 | 0.98 |
| Benztropine mesylate | 1.00 | 0.99 | 0.99 |
| Bismuth Subsalicylate | 1.07 | 1.05 | 1.06 |
| Amfenac Sodium Monohydrate | 1.14 | 1.10 | 1.12 |
| Pyrimethamine | 1.05 | 1.04 | 1.04 |
| Moxonidine | 1.00 | 1.04 | 1.02 |
| Dextrose | 1.05 | 1.04 | 1.05 |
| Fenbendazole | 1.11 | 0.89 | 1.00 |
| Phenytoin sodium | 1.00 | 0.97 | 0.99 |
| Tolvaptan | 1.12 | 1.01 | 1.06 |
| Bindarit | 0.89 | 0.98 | 0.94 |
| Flumequine | 1.13 | 0.99 | 1.06 |
| Pheniramine Maleate | 0.99 | 0.98 | 0.99 |
| Penicillin G Sodium | 1.12 | 1.08 | 1.10 |
| Ginkgolide A | 0.97 | 0.89 | 0.93 |
| Cilazapril Monohydrate | 0.84 | 0.92 | 0.88 |

| | | | |
|---|---|---|---|
| Dabigatran Etexilate | 1.09 | 0.99 | 1.04 |
| Moroxydine HCl | 0.94 | 1.05 | 0.99 |
| Lomefloxacin HCl | 0.87 | 1.10 | 0.98 |
| Buflomedil HCl | 1.03 | 0.95 | 0.99 |
| Otilonium Bromide | 1.06 | 0.98 | 1.02 |
| Catharanthine | 0.94 | 0.94 | 0.94 |
| Ropivacaine HCl | 1.21 | 1.00 | 1.11 |
| 1-Hexadecanol | 0.95 | 1.11 | 1.03 |
| Penciclovir | 0.88 | 1.06 | 0.97 |
| Pipenzolate Bromide | 0.88 | 1.12 | 1.00 |
| Ethacridine lactate monohydrate | 0.77 | 0.93 | 0.85 |
| Pimozide | 0.92 | 1.02 | 0.97 |
| Chromocarb | 1.06 | 0.96 | 1.01 |
| Hydrastinine HCl | 0.88 | 1.02 | 0.95 |
| Piperacillin Sodium | 0.87 | 0.91 | 0.89 |
| Nomifensine Maleate | 0.83 | 0.86 | 0.85 |
| Benfotiamine | 1.13 | 0.99 | 1.06 |
| Camylofin Chlorhydrate | 0.92 | 0.90 | 0.91 |
| Carbadox | 1.15 | 1.07 | 1.11 |
| Oxeladin Citrate | 1.12 | 0.95 | 1.04 |
| Physostigmine Hemisulfate Salt | 1.06 | 1.20 | 1.13 |
| Metaraminol Bitartrate | 0.87 | 0.99 | 0.93 |

1) Two independent experiments were performed.
2) Red indicates more than 1.25 while green indicates less than 0.80.
3) Toxicity was detemined by microscopic observation.

**Supplementary Table 3. Summary of the second screening**

| Compounds | Targets/Remarks | Relative GeoMean of HLA-DR[1),2)] | | |
|---|---|---|---|---|
| | | n = 1 | n = 2 | mean |
| Chloroxine | Quinoline derivative | 1.56 | 1.59 | 1.57 |
| Dipyridamole | Phosphodiesterase V inhibitor | 1.57 | 1.47 | 1.52 |
| Alprostadil | Prostaglandin E1 | 1.27 | 1.30 | 1.28 |
| Emtricitabine | HIV reverse transcriptase inhibitor | 1.32 | 1.24 | 1.28 |
| Chlortetracycline HCl | Tetracycline analogue | 1.29 | 1.26 | 1.27 |
| Carbamazepine | Na+ channel blockers | 1.33 | 1.22 | 1.27 |
| Trometamol | pH buffering agent, also known as Tris | 1.26 | 1.28 | 1.27 |
| Chlorocresol | Cresol derivative | 1.26 | 1.25 | 1.26 |
| Isoetharine Mesylate | β2 adrenergic receptor agonist | 1.08 | 1.07 | 1.08 |
| Esmolol HCl | β1 adrenergic receptor antagonist | 1.02 | 1.08 | 1.05 |
| Uridine | Pyrimidine synthesis promotion | 0.98 | 1.02 | 1.00 |
| Isotretinoin | Retinoic acid receptor agonist | 1.02 | 0.93 | 0.98 |
| Geniposide | Iridoid glycoside from gardenia | 0.79 | 1.13 | 0.96 |
| Adefovir Dipivoxil | HBV reverse transcriptase inhibitor | 1.06 | 0.84 | 0.95 |
| Hygromycin B | Aminoglycoside | 0.86 | 1.01 | 0.94 |
| Riboflavin | Vitamin B2 | 0.73 | 1.13 | 0.93 |
| Misoprostol | Prostaglandin E1 | 1.08 | 0.78 | 0.93 |
| Menadione | Vitamin K | 0.87 | 0.94 | 0.91 |
| Tazarotene | Retinoic acid receptor agonist | 0.86 | 0.92 | 0.89 |
| Bexarotene | Retinoid X receptor agonist | 0.80 | 0.95 | 0.88 |
| Acitretin | Retinoic acid receptor agonist | 0.79 | 0.80 | 0.79 |
| Sunitinib Malate | Multi-kinase inhibitor | 0.85 | 0.70 | 0.77 |
| R-(-)-Apomorphine HCl Hemihydrate | Dopamine receptor agonist | 0.70 | 0.82 | 0.76 |
| Sulindac | Cyclooxidase inhibitor | 0.77 | 0.74 | 0.76 |
| Ropinirole HCl | Dopamine D2 receptor agonist | 0.71 | 0.75 | 0.73 |
| Rosiglitazone | Peroxisome proliferator-activated receptor γ agonist | 0.69 | 0.76 | 0.73 |
| Everolimus (RAD001) | Mechanistic target of rapamycin inhibitor | 0.66 | 0.76 | 0.71 |
| Trifluoperazine 2HCl | Dopamine D2 receptor antagonist | 0.53 | 0.80 | 0.67 |
| Calcitriol | Vitamin D receptor agonist | 0.56 | 0.62 | 0.59 |
| Calcifediol | Vitamin D receptor agonist | 0.50 | 0.67 | 0.59 |
| Dequalinium chloride | Quinoline derivative, quaternary ammonium | 0.27 | 0.75 | 0.51 |
| Sorafenib Tosylate | Multi-kinase inhibitor | 0.24 | 0.44 | 0.34 |
| Halobetasol Propionate | Glucocorticoid receptor agonist | 0.20 | 0.22 | 0.21 |
| Flumethasone | Glucocorticoid receptor agonist | 0.08 | 0.23 | 0.16 |

1) Two independent experiments were performed.

2) Red indicates more than 1.25 while blue indicates less than 0.80.

**Supplementary Table 4. DEGs in LC cells and macrophages after HX531 treatment**

| Gene | HLF cells | | THP-1-derived macrophages | | UP/DOWN |
|---|---|---|---|---|---|
| | fold changes[1),2)] | adjusted P values | fold changes[1),2)] | adjusted P values | |
| AARS1 | 1.59 | 0.022 | **2.73** | **0.019** | Mac UP |
| ABCA10 | **2.87** | **0.043** | 1.50 | 0.676 | LC UP |
| ABCC3 | 1.70 | 0.194 | **2.40** | **0.038** | Mac UP |
| ABLIM3 | **2.39** | **0.023** | 2.41 | 0.059 | LC UP |
| ACKR3 | 0.84 | 0.797 | **0.49** | **0.032** | Mac DOWN |
| ACSL1 | 2.31 | 0.136 | **2.23** | **0.037** | Mac UP |
| ADM2 | 1.61 | 0.088 | **13.06** | **0.043** | Mac UP |
| ADORA3 | 0.42 | 0.646 | **0.32** | **0.043** | Mac DOWN |
| AKAP5 | 0.82 | 0.510 | **2.37** | **0.038** | Mac UP |
| ALDH1L2 | 3.78 | 0.395 | **13.30** | **0.024** | Mac UP |
| ALPK2 | 0.75 | 0.243 | **2.41** | **0.050** | Mac UP |
| ALS2CL | 0.88 | 0.540 | **2.09** | **0.011** | Mac UP |
| ANKRD37 | **0.42** | **0.008** | 0.44 | 0.225 | LC DOWN |
| ANLN | 0.79 | 0.120 | **0.30** | **0.029** | Mac DOWN |
| APBB1 | 0.80 | 0.269 | **0.47** | **0.048** | Mac DOWN |
| APOC1 | 0.44 | 0.299 | **0.26** | **0.040** | Mac DOWN |
| APOC2 | 0.46 | 0.260 | **0.28** | **0.027** | Mac DOWN |
| APOC4-APOC2 | 0.76 | 0.707 | **0.42** | **0.038** | Mac DOWN |
| AQP1 | 0.47 | 0.381 | **0.27** | **0.027** | Mac DOWN |
| ARAP2 | 1.36 | 0.238 | **3.09** | **0.044** | Mac UP |
| ARHGAP11A | 0.87 | 0.711 | **0.46** | **0.016** | Mac DOWN |
| ARHGAP18 | 0.94 | 0.736 | **0.43** | **0.037** | Mac DOWN |
| ARHGDIB | 0.52 | 0.187 | **0.38** | **0.011** | Mac DOWN |
| ARHGEF35 | 1.01 | 0.920 | **2.36** | **0.047** | Mac UP |
| ARHGEF5 | 0.95 | 0.671 | **2.23** | **0.021** | Mac UP |
| ARVCF | 0.88 | 0.611 | **2.86** | **0.037** | Mac UP |
| ASF1B | 0.34 | 0.160 | **0.40** | **0.012** | Mac DOWN |
| ASNS | **2.69** | **0.016** | **15.27** | **0.030** | Common UP |
| ATAD2 | 0.83 | 0.130 | **0.44** | **0.043** | Mac DOWN |
| ATF4 | 1.41 | 0.057 | **2.45** | **0.011** | Mac UP |
| ATF5 | 1.05 | 0.718 | **3.22** | **0.013** | Mac UP |
| ATP8B4 | 0.25 | 0.173 | **0.28** | **0.040** | Mac DOWN |
| AURKA | 0.79 | 0.240 | **0.49** | **0.011** | Mac DOWN |
| AURKB | 0.81 | 0.537 | **0.31** | **0.016** | Mac DOWN |
| BBS2 | 1.14 | 0.059 | **2.04** | **0.015** | Mac UP |
| BCL2L1 | **0.41** | **0.007** | 0.64 | 0.116 | LC DOWN |
| BEND6 | 1.28 | 0.264 | **2.23** | **0.025** | Mac UP |
| BEST1 | 1.41 | 0.395 | **2.22** | **0.018** | Mac UP |
| BIRC5 | 0.79 | 0.222 | **0.39** | **0.018** | Mac DOWN |
| BIRC7 | 0.75 | 0.611 | **2.42** | **0.038** | Mac UP |
| BLM | 0.78 | 0.632 | **0.42** | **0.046** | Mac DOWN |
| BRPF3 | 1.22 | 0.056 | **2.01** | **0.033** | Mac UP |
| BST2 | 0.08 | 0.102 | **0.43** | **0.025** | Mac DOWN |
| BUB1B | 0.66 | 0.436 | **0.36** | **0.032** | Mac DOWN |
| BVES | 1.51 | 0.400 | **2.27** | **0.015** | Mac UP |
| C1QTNF6 | 0.69 | 0.136 | **3.56** | **0.017** | Mac UP |
| C1RL-AS1 | 1.83 | 0.824 | **3.18** | **0.024** | Mac UP |
| CACNA1F | 2.04 | 0.629 | **2.19** | **0.042** | Mac UP |
| CACNA1I | **0.07** | **0.026** | 0.70 | 0.647 | LC DOWN |
| CAPN5 | 1.11 | 0.531 | **2.58** | **0.018** | Mac UP |
| CARF | 1.37 | 0.229 | **2.61** | **0.035** | Mac UP |
| CARS1 | 1.26 | 0.088 | **2.66** | **0.015** | Mac UP |
| CBS | 1.36 | 0.089 | **3.23** | **0.029** | Mac UP |
| CCDC112 | 1.42 | 0.187 | **2.10** | **0.034** | Mac UP |
| CCDC15-DT | 1.17 | 0.785 | **10.01** | **0.040** | Mac UP |
| CCDC154 | 0.71 | 0.764 | **2.85** | **0.026** | Mac UP |
| CCDC24 | 1.14 | 0.336 | **2.13** | **0.020** | Mac UP |

| | | | | | |
|---|---|---|---|---|---|
| CCL20 | 0.78 | 0.190 | **0.42** | **0.012** | Mac DOWN |
| CCL4 | 0.63 | 0.500 | **0.41** | **0.038** | Mac DOWN |
| CCL7 | 0.25 | 0.387 | **0.31** | **0.037** | Mac DOWN |
| CCNA2 | 0.91 | 0.706 | **0.41** | **0.015** | Mac DOWN |
| CCNB1 | 0.90 | 0.612 | **0.38** | **0.034** | Mac DOWN |
| CCNB2 | 0.96 | 0.830 | **0.42** | **0.042** | Mac DOWN |
| CCNE2 | 0.57 | 0.143 | **0.26** | **0.048** | Mac DOWN |
| CCNF | 0.97 | 0.735 | **0.42** | **0.016** | Mac DOWN |
| CCPG1 | 1.38 | 0.254 | **2.19** | **0.015** | Mac UP |
| CCSER2 | 1.39 | 0.128 | **2.22** | **0.021** | Mac UP |
| CD14 | 0.64 | 0.400 | **0.47** | **0.045** | Mac DOWN |
| CD163 | 3.10 | 0.322 | **0.38** | **0.014** | Mac DOWN |
| CD36 | 0.50 | 0.231 | **0.47** | **0.040** | Mac DOWN |
| CD74 | **0.38** | **0.036** | 0.69 | 0.193 | LC DOWN |
| CD86 | 0.65 | 0.728 | **0.34** | **0.014** | Mac DOWN |
| CDC20 | 0.90 | 0.743 | **0.49** | **0.037** | Mac DOWN |
| CDC25A | 0.71 | 0.327 | **0.38** | **0.030** | Mac DOWN |
| CDC25C | 0.68 | 0.347 | **0.35** | **0.027** | Mac DOWN |
| CDC45 | 0.34 | 0.082 | **0.41** | **0.041** | Mac DOWN |
| CDC7 | 1.06 | 0.672 | **0.40** | **0.018** | Mac DOWN |
| CDCA3 | 0.77 | 0.406 | **0.36** | **0.046** | Mac DOWN |
| CDCA5 | 0.81 | 0.278 | **0.41** | **0.018** | Mac DOWN |
| CDCA8 | 0.98 | 0.913 | **0.40** | **0.018** | Mac DOWN |
| CDK1 | 0.78 | 0.138 | **0.30** | **0.012** | Mac DOWN |
| CDKN2D | 0.66 | 0.122 | **0.49** | **0.025** | Mac DOWN |
| CDKN3 | 0.70 | 0.332 | **0.37** | **0.035** | Mac DOWN |
| CDT1 | 0.67 | 0.059 | **0.44** | **0.016** | Mac DOWN |
| CEBPG | 1.61 | 0.062 | **2.32** | **0.021** | Mac UP |
| CELF2 | 0.45 | 0.466 | **0.48** | **0.047** | Mac DOWN |
| CENPA | 1.30 | 0.445 | **0.41** | **0.025** | Mac DOWN |
| CENPE | 0.89 | 0.672 | **0.49** | **0.049** | Mac DOWN |
| CENPQ | 1.18 | 0.591 | **0.46** | **0.016** | Mac DOWN |
| CENPW | 0.99 | 0.990 | **0.39** | **0.050** | Mac DOWN |
| CFAP77 | | | **720.77**[3)] | **0.034** | Mac UP |
| CHAC1 | 1.60 | 0.101 | **16.46** | **0.050** | Mac UP |
| CHRNA5 | 0.74 | 0.203 | **0.33** | **0.020** | Mac DOWN |
| CKMT2-AS1 | 1.37 | 0.297 | **2.02** | **0.026** | Mac UP |
| CKS2 | 1.04 | 0.780 | **0.44** | **0.020** | Mac DOWN |
| CLCN5 | 1.83 | 0.066 | **2.14** | **0.021** | Mac UP |
| CLGN | 4.26 | 0.302 | **9.05** | **0.044** | Mac UP |
| CLIP4 | 1.43 | 0.118 | **2.21** | **0.039** | Mac UP |
| CLSPN | 0.67 | 0.095 | **0.37** | **0.029** | Mac DOWN |
| CMKLR1 | 0.00 | 0.615 | **0.37** | **0.038** | Mac DOWN |
| CMPK2 | **0.09** | **0.018** | 0.10 | 0.099 | LC DOWN |
| COPB2-DT | 1.57 | 0.484 | **2.40** | **0.024** | Mac UP |
| CORO7 | 0.72 | 0.089 | **0.48** | **0.037** | Mac DOWN |
| CPEB1-AS1 | 0.66 | 0.646 | **3.04** | **0.027** | Mac UP |
| CSRP1-AS1 | 1.15 | 0.732 | **2.67** | **0.042** | Mac UP |
| CST1 | 0.26 | 0.209 | **0.03** | **0.030** | Mac DOWN |
| CTNNA3 | 1.74 | 0.837 | **5.21** | **0.040** | Mac UP |
| CXCL10 | **0.10** | **0.021** | 0.18 | 0.246 | LC DOWN |
| CXCL11 | **0.14** | **0.032** | 0.05 | 0.662 | LC DOWN |
| CYP19A1 | 0.90 | 0.741 | **0.45** | **0.045** | Mac DOWN |
| CYP27A1 | 1.08 | 0.565 | **2.93** | **0.034** | Mac UP |
| CYP4V2 | 1.55 | 0.142 | **3.01** | **0.039** | Mac UP |
| D2HGDH | 1.03 | 0.427 | **2.00** | **0.024** | Mac UP |
| DCAF10 | 1.66 | 0.032 | **2.76** | **0.013** | Mac UP |
| DENND2B | 1.02 | 0.766 | **0.40** | **0.042** | Mac DOWN |
| DENND2D | 0.58 | 0.517 | **2.01** | **0.012** | Mac UP |
| DEPDC1B | 0.74 | 0.562 | **0.41** | **0.027** | Mac DOWN |

| | | | | | |
|---|---|---|---|---|---|
| DEPDC7 | 1.01 | 0.974 | 0.37 | 0.050 | Mac DOWN |
| DEPP1 | 0.38 | 0.150 | 0.49 | 0.044 | Mac DOWN |
| DET1 | 1.20 | 0.663 | 3.19 | 0.034 | Mac UP |
| DHRS11 | 1.09 | 0.712 | 2.89 | 0.019 | Mac UP |
| DHRS12 | 1.01 | 0.970 | 2.03 | 0.041 | Mac UP |
| DIAPH3 | 0.67 | 0.320 | 0.40 | 0.042 | Mac DOWN |
| DLGAP1-AS2 | 2.23 | 0.068 | 2.24 | 0.026 | Mac UP |
| DLGAP5 | 0.73 | 0.547 | 0.31 | 0.015 | Mac DOWN |
| DMGDH | 2.00 | 0.728 | 9.61 | 0.024 | Mac UP |
| DNAI4 | 2.18 | 0.094 | 2.02 | 0.025 | Mac UP |
| DSCC1 | 0.90 | 0.769 | 0.42 | 0.032 | Mac DOWN |
| DTL | 0.59 | 0.053 | 0.35 | 0.018 | Mac DOWN |
| DTNB | 1.26 | 0.240 | 2.14 | 0.034 | Mac UP |
| E2F1 | 0.47 | 0.033 | 0.55 | 0.015 | LC DOWN |
| E2F2 | 0.69 | 0.398 | 0.22 | 0.012 | Mac DOWN |
| ECM1 | 0.64 | 0.071 | 0.47 | 0.048 | Mac DOWN |
| ELOVL6 | 1.54 | 0.102 | 2.46 | 0.040 | Mac UP |
| ENSG00000203644 | 1.33 | 0.229 | 2.16 | 0.031 | Mac UP |
| ENSG00000224404 | 2.64 | 0.573 | 0.00 | 0.049 | Mac DOWN |
| ENSG00000224810 | | | 1354.08[3)] | 0.029 | Mac UP |
| ENSG00000228703 | 6.39 | 0.514 | 0.00 | 0.049 | Mac DOWN |
| ENSG00000230612 | 4.84 | 0.710 | 0.00 | 0.049 | Mac DOWN |
| ENSG00000232811 | 0.53 | 0.833 | 0.36 | 0.041 | Mac DOWN |
| ENSG00000233478 | | | 348.23[3)] | 0.033 | Mac UP |
| ENSG00000236352 | 1.38 | 0.853 | 4.44 | 0.037 | Mac UP |
| ENSG00000238178 | | | 0.18 | 0.033 | Mac DOWN |
| ENSG00000253821 | 0.00 | 0.651 | 0.41 | 0.047 | Mac DOWN |
| ENSG00000254587 | | | 345.28[3)] | 0.033 | Mac UP |
| ENSG00000255491 | 0.24 | 0.059 | 0.44 | 0.036 | Mac DOWN |
| ENSG00000256568 | 0.00 | 0.596 | 0.27 | 0.030 | Mac DOWN |
| ENSG00000258561 | 1.07 | 0.960 | 5.05 | 0.018 | Mac UP |
| ENSG00000259066 | 4442.23[3)] | 0.769 | 9510.67[3)] | 0.038 | Mac UP |
| ENSG00000259661 | 0.67 | 0.755 | 11.92 | 0.030 | Mac UP |
| ENSG00000260331 | 0.66 | 0.790 | 1891.24[3)] | 0.033 | Mac UP |
| ENSG00000261172 | | | 2.69 | 0.048 | Mac UP |
| ENSG00000261625 | | | 0.00 | 0.049 | Mac DOWN |
| ENSG00000266709 | 0.57 | 0.138 | 0.24 | 0.013 | Mac DOWN |
| ENSG00000267385 | 0.79 | 0.852 | 0.36 | 0.044 | Mac DOWN |
| ENSG00000267765 | 2.04 | 0.693 | 2154.28[3)] | 0.033 | Mac UP |
| ENSG00000270207 | | | 5.40 | 0.049 | Mac UP |
| ENSG00000271698 | 26.84 | 0.138 | 11.89 | 0.035 | Mac UP |
| ENSG00000272405 | 1.10 | 0.709 | 2.67 | 0.033 | Mac UP |
| ENSG00000272449 | 0.00 | 0.630 | 0.37 | 0.032 | Mac DOWN |
| ENSG00000272668 | 1.09 | 0.403 | 2.46 | 0.032 | Mac UP |
| ENSG00000273192 | 1.10 | 0.756 | 3.91 | 0.045 | Mac UP |
| ENSG00000273230 | 0.88 | 0.550 | 2.64 | 0.034 | Mac UP |
| ENSG00000274767 | 1.00 | 1.001 | 2.43 | 0.022 | Mac UP |
| ENSG00000274922 | 1.12 | 0.781 | 2.09 | 0.023 | Mac UP |
| ENSG00000274929 | 1.43 | 0.907 | 0.26 | 0.044 | Mac DOWN |
| ENSG00000275389 | 1.06 | 0.948 | 12.21 | 0.029 | Mac UP |
| ENSG00000276075 | 0.97 | 0.960 | 18.00 | 0.018 | Mac UP |
| ENSG00000278730 | 1.28 | 0.073 | 2.19 | 0.036 | Mac UP |
| ENSG00000279406 | 0.12 | 0.105 | 0.40 | 0.021 | Mac DOWN |
| ENSG00000280160 | 0.14 | 0.041 | 0.24 | 0.109 | LC DOWN |
| ENSG00000286153 | 4.02 | 0.228 | 2.13 | 0.030 | Mac UP |
| ENSG00000286448 | 1.76 | 0.658 | 3.51 | 0.032 | Mac UP |
| ENSG00000286471 | 3.09 | 0.230 | 2.49 | 0.042 | Mac UP |
| ENSG00000287069 | 1.32 | 0.392 | 2.31 | 0.018 | Mac UP |
| ENSG00000287300 | 1.13 | 0.903 | 606.00[3)] | 0.033 | Mac UP |
| ENSG00000287566 | | | 0.00 | 0.049 | Mac DOWN |

| | | | | | |
|---|---|---|---|---|---|
| ENSG00000287733 | 0.79 | 0.705 | 0.42 | 0.018 | Mac DOWN |
| ENSG00000288085 | 2.08 | 0.686 | 0.00 | 0.049 | Mac DOWN |
| ENSG00000288973 | 1.30 | 0.265 | 2.28 | 0.028 | Mac UP |
| ENSG00000289009 | 0.36 | 0.021 | 0.78 | 0.282 | LC DOWN |
| ENSG00000289046 | 0.81 | 0.666 | 0.15 | 0.047 | Mac DOWN |
| ENSG00000289142 | 0.94 | 0.980 | 0.40 | 0.048 | Mac DOWN |
| ENSG00000289316 | 0.60 | 0.724 | 0.18 | 0.043 | Mac DOWN |
| ENSG00000289507 | 0.70 | 0.773 | 0.47 | 0.030 | Mac DOWN |
| ENSG00000289754 | 1.25 | 0.785 | 3.13 | 0.036 | Mac UP |
| ENSG00000289915 | 1.64 | 0.640 | 10.22 | 0.040 | Mac UP |
| ENSG00000290105 | 1.06 | 0.912 | 0.44 | 0.036 | Mac DOWN |
| ENSG00000290383 | 0.80 | 0.759 | 2.39 | 0.026 | Mac UP |
| ENSG00000290394 | 0.93 | 0.760 | 2.43 | 0.040 | Mac UP |
| ENSG00000290396 | 1.05 | 0.833 | 2.00 | 0.037 | Mac UP |
| ENSG00000290679 | | | 0.17 | 0.042 | Mac DOWN |
| ENSG00000290796 | 1.09 | 0.664 | 2.39 | 0.024 | Mac UP |
| ENSG00000290890 | 1.27 | 0.518 | 2.42 | 0.037 | Mac UP |
| ENSG00000291044 | 1.53 | 0.217 | 0.27 | 0.041 | Mac DOWN |
| ENSG00000291207 | 1.35 | 0.136 | 2.07 | 0.026 | Mac UP |
| ENSG00000291270 | 1.17 | 0.656 | 2.03 | 0.034 | Mac UP |
| ENSG00000291283 | | | 4.28 | 0.050 | Mac UP |
| EPAS1 | 1.69 | 0.054 | 2.48 | 0.037 | Mac UP |
| EPRS1 | 1.47 | 0.086 | 2.19 | 0.046 | Mac UP |
| EPSTI1 | 0.17 | 0.039 | 0.12 | 0.050 | LC DOWN |
| ERCC6L | 0.52 | 0.252 | 0.39 | 0.049 | Mac DOWN |
| ERMAP | 1.28 | 0.235 | 2.15 | 0.014 | Mac UP |
| ESCO2 | 0.47 | 0.157 | 0.31 | 0.021 | Mac DOWN |
| ESPL1 | 0.72 | 0.307 | 0.41 | 0.036 | Mac DOWN |
| ETV7 | 0.27 | 0.354 | 0.49 | 0.040 | Mac DOWN |
| EXO1 | 0.82 | 0.561 | 0.33 | 0.013 | Mac DOWN |
| EZH2 | 0.96 | 0.818 | 0.47 | 0.035 | Mac DOWN |
| F5 | 0.33 | 0.342 | 0.36 | 0.033 | Mac DOWN |
| FABP5 | 0.80 | 0.150 | 0.45 | 0.031 | Mac DOWN |
| FADS2 | 0.98 | 0.806 | 0.43 | 0.029 | Mac DOWN |
| FAM210B | 0.92 | 0.516 | 2.52 | 0.018 | Mac UP |
| FAM219B | 1.01 | 0.871 | 2.05 | 0.046 | Mac UP |
| FAM66B | 1.79 | 0.629 | 2.04 | 0.046 | Mac UP |
| FAM72B | 1.20 | 0.625 | 0.38 | 0.019 | Mac DOWN |
| FAM72D | 1.57 | 0.369 | 0.36 | 0.041 | Mac DOWN |
| FAM83A | 0.35 | 0.088 | 4.41 | 0.047 | Mac UP |
| FAM86JP | 1.73 | 0.181 | 3.06 | 0.048 | Mac UP |
| FANCI | 0.55 | 0.100 | 0.46 | 0.047 | Mac DOWN |
| FAP | 0.76 | 0.593 | 0.38 | 0.014 | Mac DOWN |
| FASN | 0.46 | 0.049 | 0.50 | 0.075 | LC DOWN |
| FBXO10 | 1.25 | 0.515 | 2.59 | 0.030 | Mac UP |
| FBXO27 | 1.69 | 0.460 | 2.99 | 0.025 | Mac UP |
| FBXO44 | 1.22 | 0.516 | 2.10 | 0.030 | Mac UP |
| FBXO5 | 1.13 | 0.387 | 0.44 | 0.015 | Mac DOWN |
| FBXW4 | 1.21 | 0.225 | 2.10 | 0.013 | Mac UP |
| FCGR3A | 0.32 | 0.438 | 0.20 | 0.046 | Mac DOWN |
| FCRLB | 1.53 | 0.510 | 0.15 | 0.014 | Mac DOWN |
| FECH | 1.30 | 0.139 | 2.03 | 0.015 | Mac UP |
| FEN1 | 0.77 | 0.231 | 0.47 | 0.014 | Mac DOWN |
| FGR | 1.66 | 0.143 | 2.58 | 0.011 | Mac UP |
| FKBP14 | 2.09 | 0.009 | 0.99 | 0.897 | LC UP |
| FLNB-AS1 | 4.46 | 0.603 | 0.00 | 0.049 | Mac DOWN |
| FLYWCH1 | 1.32 | 0.105 | 2.03 | 0.018 | Mac UP |
| FOXD3 | | | 2.96 | 0.024 | Mac UP |
| FOXM1 | 0.59 | 0.066 | 0.42 | 0.011 | Mac DOWN |
| FPR3 | 0.00 | 0.629 | 0.13 | 0.024 | Mac DOWN |

| | | | | | |
|---|---|---|---|---|---|
| FTCDNL1 | 2.71 | 0.108 | **2.65** | **0.027** | Mac UP |
| FTH1 | 1.52 | 0.050 | **2.17** | **0.014** | Mac UP |
| FXYD2 | 0.50 | 0.138 | **0.11** | **0.016** | Mac DOWN |
| G0S2 | 1.02 | 0.682 | **0.47** | **0.043** | Mac DOWN |
| GAB2 | 1.58 | 0.096 | **2.34** | **0.020** | Mac UP |
| GABRG1 | **5.80** | **0.050** | 1.82 | 0.707 | LC UP |
| GDF1 | 1.12 | 0.868 | **2.14** | **0.044** | Mac UP |
| GGH | 0.77 | 0.111 | **0.45** | **0.037** | Mac DOWN |
| GINS1 | 0.58 | 0.109 | **0.36** | **0.027** | Mac DOWN |
| GINS2 | 0.37 | 0.205 | **0.38** | **0.035** | Mac DOWN |
| GJD3-AS1 | 0.96 | 0.686 | **2.67** | **0.025** | Mac UP |
| GMNN | 0.90 | 0.373 | **0.40** | **0.018** | Mac DOWN |
| GPCPD1 | 1.68 | 0.038 | **2.43** | **0.014** | Mac UP |
| GPR15 | **0.46** | **0.048** | 2.82 | 0.699 | LC DOWN |
| GPR156 | **2.79** | **0.021** | 0.66 | 0.308 | LC UP |
| GPSM2 | 1.27 | 0.167 | **0.48** | **0.032** | Mac DOWN |
| GPT2 | 1.08 | 0.516 | **2.24** | **0.034** | Mac UP |
| GRB10 | 2.37 | 0.401 | **3.62** | **0.012** | Mac UP |
| GRIA2 | 1.73 | 0.662 | **11.90** | **0.038** | Mac UP |
| GRIP2 | 0.37 | 0.263 | **0.38** | **0.015** | Mac DOWN |
| GSDMB | 0.97 | 0.891 | **2.07** | **0.032** | Mac UP |
| GTF2IRD2B | 1.56 | 0.236 | **2.30** | **0.025** | Mac UP |
| GTPBP2 | 1.30 | 0.120 | **2.07** | **0.013** | Mac UP |
| GTSE1 | 0.57 | 0.109 | **0.37** | **0.014** | Mac DOWN |
| H2AC18 | 0.36 | 0.249 | **0.50** | **0.018** | Mac DOWN |
| H2AC19 | 0.67 | 0.428 | **0.49** | **0.018** | Mac DOWN |
| H2BC21 | 0.66 | 0.307 | **0.40** | **0.024** | Mac DOWN |
| HAVCR2 | 1.71 | 0.338 | **0.49** | **0.041** | Mac DOWN |
| HCG20 | 1.89 | 0.636 | **3.54** | **0.038** | Mac UP |
| HDAC4-AS1 | 0.88 | 0.726 | **0.47** | **0.033** | Mac DOWN |
| HELLS | 0.68 | 0.106 | **0.32** | **0.034** | Mac DOWN |
| HEMK1 | 1.50 | 0.088 | **2.29** | **0.044** | Mac UP |
| HERC2P3 | 1.10 | 0.394 | **2.01** | **0.033** | Mac UP |
| HERC5 | **0.36** | **0.047** | 0.32 | 0.117 | LC DOWN |
| HERC6 | 0.38 | 0.059 | **0.32** | **0.049** | Mac DOWN |
| HILPDA | 0.57 | 0.057 | **0.35** | **0.039** | Mac DOWN |
| HIP1 | 1.06 | 0.420 | **3.63** | **0.022** | Mac UP |
| HJURP | 0.82 | 0.160 | **0.32** | **0.025** | Mac DOWN |
| HK3 | 0.35 | 0.432 | **0.41** | **0.049** | Mac DOWN |
| HMGB3 | 1.03 | 0.661 | **0.49** | **0.018** | Mac DOWN |
| HMMR | 0.62 | 0.258 | **0.40** | **0.014** | Mac DOWN |
| HPSE | 1.24 | 0.249 | **0.23** | **0.017** | Mac DOWN |
| HSF4 | 1.05 | 0.840 | **2.21** | **0.041** | Mac UP |
| HSH2D | **0.12** | **0.035** | 0.12 | 0.122 | LC DOWN |
| HSPA1A | 0.52 | 0.112 | **0.42** | **0.042** | Mac DOWN |
| HSPA1B | 0.50 | 0.127 | **0.47** | **0.049** | Mac DOWN |
| HSPA6 | **0.10** | **0.038** | **0.16** | **0.014** | Common DOWN |
| HTD2 | 0.98 | 0.980 | **2.88** | **0.048** | Mac UP |
| HTR2C | 0.00 | 0.769 | **319.89**[3)] | **0.033** | Mac UP |
| IARS1 | 1.78 | 0.018 | **2.22** | **0.012** | Mac UP |
| ICA1 | 1.50 | 0.102 | **2.25** | **0.017** | Mac UP |
| ID1 | 0.76 | 0.384 | **0.47** | **0.033** | Mac DOWN |
| IDNK | 1.34 | 0.300 | **2.02** | **0.036** | Mac UP |
| IER5L | **0.42** | **0.023** | 0.85 | 0.222 | LC DOWN |
| IFI35 | **0.39** | **0.015** | **0.42** | **0.033** | Common DOWN |
| IFI44 | **0.40** | **0.039** | 0.07 | 0.053 | LC DOWN |
| IFI44L | **0.30** | **0.024** | 0.06 | 0.120 | LC DOWN |
| IFI6 | 0.13 | 0.100 | **0.16** | **0.024** | Mac DOWN |
| IFIH1 | **0.40** | **0.044** | 0.38 | 0.066 | LC DOWN |
| IFIT5 | **0.38** | **0.037** | **0.32** | **0.032** | Common DOWN |

| | | | | | |
|---|---|---|---|---|---|
| IFITM1 | 0.05 | 0.127 | **0.07** | **0.042** | Mac DOWN |
| IFITM3 | **0.36** | **0.048** | 0.33 | 0.083 | LC DOWN |
| IFRD1 | 1.48 | 0.067 | **2.09** | **0.012** | Mac UP |
| IGSF6 | 0.22 | 0.603 | **0.25** | **0.021** | Mac DOWN |
| IL1RN | 0.48 | 0.147 | **0.41** | **0.012** | Mac DOWN |
| IL7R | 0.89 | 0.601 | **0.43** | **0.025** | Mac DOWN |
| INAVA | 0.66 | 0.198 | **0.18** | **0.033** | Mac DOWN |
| INCENP | 1.07 | 0.820 | **0.49** | **0.014** | Mac DOWN |
| INHBE | 1.94 | 0.395 | **3.12** | **0.014** | Mac UP |
| IPCEF1 | 0.94 | 0.956 | **0.37** | **0.045** | Mac DOWN |
| IQGAP3 | 0.60 | 0.260 | **0.42** | **0.016** | Mac DOWN |
| IRF7 | 0.93 | 0.863 | **0.40** | **0.023** | Mac DOWN |
| IRF8 | 0.54 | 0.261 | **0.37** | **0.021** | Mac DOWN |
| IRF9 | 0.60 | 0.058 | **0.48** | **0.021** | Mac DOWN |
| ISG15 | **0.16** | **0.044** | **0.16** | **0.042** | Common DOWN |
| ISG20 | **0.27** | **0.016** | 0.25 | 0.139 | LC DOWN |
| ITGA1 | 0.74 | 0.363 | **0.39** | **0.047** | Mac DOWN |
| ITGAM | 0.63 | 0.166 | **0.50** | **0.018** | Mac DOWN |
| JDP2 | 1.68 | 0.100 | **2.09** | **0.014** | Mac UP |
| KCNA2 | 2.88 | 0.217 | **2.22** | **0.036** | Mac UP |
| KCNMB3 | 1.24 | 0.707 | **2.72** | **0.033** | Mac UP |
| KCTD15 | 1.75 | 0.069 | **2.60** | **0.036** | Mac UP |
| KDM5D | 0.79 | 0.630 | **2.06** | **0.029** | Mac UP |
| KIF15 | 0.58 | 0.332 | **0.45** | **0.022** | Mac DOWN |
| KIF18A | 0.99 | 0.954 | **0.37** | **0.014** | Mac DOWN |
| KIF18B | 0.67 | 0.285 | **0.32** | **0.015** | Mac DOWN |
| KIF20A | 0.77 | 0.520 | **0.25** | **0.022** | Mac DOWN |
| KIF23 | 0.83 | 0.458 | **0.39** | **0.048** | Mac DOWN |
| KIF27 | 1.74 | 0.123 | **2.31** | **0.016** | Mac UP |
| KIF2C | 0.75 | 0.106 | **0.30** | **0.022** | Mac DOWN |
| KIF3C | 0.94 | 0.418 | **0.49** | **0.037** | Mac DOWN |
| KIF4A | 0.91 | 0.672 | **0.36** | **0.015** | Mac DOWN |
| KIFC1 | 0.63 | 0.110 | **0.34** | **0.011** | Mac DOWN |
| KLF8 | 1.39 | 0.745 | **2.12** | **0.040** | Mac UP |
| KNL1 | 0.57 | 0.337 | **0.42** | **0.040** | Mac DOWN |
| LAMB3 | 0.62 | 0.189 | **0.39** | **0.014** | Mac DOWN |
| LAP3 | **0.50** | **0.016** | 0.48 | 0.051 | LC DOWN |
| LGALS3 | 1.04 | 0.778 | **2.61** | **0.015** | Mac UP |
| LGALSL | 1.14 | 0.611 | **2.25** | **0.047** | Mac UP |
| LILRA1 | 2.46 | 0.778 | **0.26** | **0.050** | Mac DOWN |
| LILRB2 | | | **0.39** | **0.014** | Mac DOWN |
| LIMA1 | 1.36 | 0.147 | **0.49** | **0.032** | Mac DOWN |
| LINC00174 | 1.73 | 0.178 | **2.34** | **0.047** | Mac UP |
| LINC00638 | 0.80 | 0.696 | **3.05** | **0.027** | Mac UP |
| LINC00662 | 1.45 | 0.303 | **2.36** | **0.043** | Mac UP |
| LINC01127 | 0.63 | 0.739 | **0.29** | **0.015** | Mac DOWN |
| LINC01186 | **0.30** | **0.044** | 0.00 | 0.730 | LC DOWN |
| LINC01235 | 4.26 | 0.455 | **0.24** | **0.024** | Mac DOWN |
| LINC01670 | 0.64 | 0.643 | **0.18** | **0.038** | Mac DOWN |
| LINC01943 | 1.04 | 0.926 | **0.38** | **0.045** | Mac DOWN |
| LINC02065 | | | **798.00**[3)] | **0.033** | Mac UP |
| LINC02193 | 2.50 | 0.249 | **3.52** | **0.048** | Mac UP |
| LINC02574 | 0.23 | 0.219 | **0.06** | **0.036** | Mac DOWN |
| LINC02611 | 0.45 | 0.622 | **0.19** | **0.013** | Mac DOWN |
| LINC02801 | 0.56 | 0.642 | **0.40** | **0.031** | Mac DOWN |
| LINC02925 | 0.97 | 0.952 | **2.14** | **0.036** | Mac UP |
| LINC03025_2 | **0.05** | **0.039** | 0.36 | 0.592 | LC DOWN |
| LIPT2-AS1 | 1.45 | 0.332 | **2.04** | **0.043** | Mac UP |
| LMNB1 | 0.75 | 0.160 | **0.27** | **0.031** | Mac DOWN |
| LNCAROD | 0.00 | 0.629 | **0.21** | **0.020** | Mac DOWN |

| | | | | | |
|---|---|---|---|---|---|
| LPCAT3 | 0.26 | 0.011 | 0.60 | 0.016 | LC DOWN |
| LPL | 0.39 | 0.520 | 0.39 | 0.036 | Mac DOWN |
| LRRC24 | 1.74 | 0.126 | 2.74 | 0.042 | Mac UP |
| LY6E | 0.47 | 0.189 | 0.37 | 0.014 | Mac DOWN |
| LYZ | 0.39 | 0.089 | 0.35 | 0.013 | Mac DOWN |
| MAB21L2 | 0.42 | 0.035 | | | LC DOWN |
| MAD2L1 | 0.80 | 0.145 | 0.36 | 0.049 | Mac DOWN |
| MAN2A2 | 1.15 | 0.279 | 2.04 | 0.015 | Mac UP |
| MAP1B | 1.40 | 0.142 | 2.62 | 0.013 | Mac UP |
| MAP7 | 1.00 | 0.993 | 2.47 | 0.032 | Mac UP |
| MARCO | 0.53 | 0.203 | 0.42 | 0.031 | Mac DOWN |
| MARS1 | 1.42 | 0.022 | 2.24 | 0.016 | Mac UP |
| MATN1-AS1 | 1.61 | 0.147 | 2.32 | 0.040 | Mac UP |
| MCM2 | 0.64 | 0.080 | 0.40 | 0.012 | Mac DOWN |
| MCM3 | 0.55 | 0.057 | 0.37 | 0.012 | Mac DOWN |
| MCM4 | 0.61 | 0.043 | 0.40 | 0.012 | Mac DOWN |
| MCM5 | 0.56 | 0.047 | 0.50 | 0.022 | Mac DOWN |
| MCOLN3 | 1.10 | 0.754 | 2.81 | 0.026 | Mac UP |
| MELK | 0.77 | 0.320 | 0.44 | 0.044 | Mac DOWN |
| Metazoa_SRP_56 | 1266.06[3)] | 0.769 | 2879.04[3)] | 0.033 | Mac UP |
| MFSD4B-DT | 0.00 | 0.644 | 0.00 | 0.049 | Mac DOWN |
| MIA2 | 1.54 | 0.190 | 2.39 | 0.022 | Mac UP |
| MID1IP1 | 0.55 | 0.043 | 0.45 | 0.012 | Mac DOWN |
| MIR3142HG | 2.14 | 0.041 | 1.61 | 0.313 | LC UP |
| MIR4435-2HG | 0.77 | 0.130 | 0.49 | 0.048 | Mac DOWN |
| MKNK2 | 1.34 | 0.066 | 2.69 | 0.037 | Mac UP |
| MMP2 | 1.05 | 0.804 | 0.30 | 0.033 | Mac DOWN |
| MMP20 | 1.26 | 0.892 | 0.00 | 0.049 | Mac DOWN |
| MND1 | 1.27 | 0.219 | 0.45 | 0.031 | Mac DOWN |
| MT-ATP6 | 1.41 | 0.089 | 0.47 | 0.024 | Mac DOWN |
| MT-ATP8 | 1.27 | 0.190 | 0.45 | 0.028 | Mac DOWN |
| MT-CO1 | 0.86 | 0.495 | 0.34 | 0.013 | Mac DOWN |
| MT-CO2 | 0.81 | 0.259 | 0.29 | 0.013 | Mac DOWN |
| MT-CO3 | 1.03 | 0.772 | 0.40 | 0.014 | Mac DOWN |
| MT-CYB | 1.16 | 0.348 | 0.45 | 0.018 | Mac DOWN |
| MT-ND4 | 1.06 | 0.677 | 0.42 | 0.024 | Mac DOWN |
| MT-ND4L | 1.09 | 0.748 | 0.44 | 0.024 | Mac DOWN |
| MT-ND5 | 0.92 | 0.653 | 0.35 | 0.018 | Mac DOWN |
| MT1E | 1.61 | 0.067 | 5.85 | 0.045 | Mac UP |
| MT1F | 13.08 | 0.251 | 7.68 | 0.035 | Mac UP |
| MT1G | 1.54 | 0.395 | 8.54 | 0.045 | Mac UP |
| MTHFD2 | 1.38 | 0.064 | 2.09 | 0.019 | Mac UP |
| MTSS2 | 2.65 | 0.136 | 3.13 | 0.015 | Mac UP |
| MUC12 | 1.30 | 0.825 | 5.62 | 0.030 | Mac UP |
| MX1 | 0.26 | 0.015 | 0.09 | 0.053 | LC DOWN |
| MXD3 | 1.11 | 0.736 | 0.48 | 0.038 | Mac DOWN |
| MYB | 0.41 | 0.523 | 0.31 | 0.029 | Mac DOWN |
| MYBL2 | 0.41 | 0.063 | 0.33 | 0.013 | Mac DOWN |
| MYBPH | 0.49 | 0.144 | 0.28 | 0.012 | Mac DOWN |
| MYLIP | 0.20 | 0.101 | 0.14 | 0.031 | Mac DOWN |
| MYO15B | 1.65 | 0.110 | 2.61 | 0.013 | Mac UP |
| NAMPT | 1.63 | 0.259 | 2.61 | 0.040 | Mac UP |
| NBL1 | 0.79 | 0.232 | 0.39 | 0.013 | Mac DOWN |
| NBPF1 | 1.39 | 0.162 | 2.46 | 0.013 | Mac UP |
| NBPF11 | 1.42 | 0.222 | 2.28 | 0.018 | Mac UP |
| NBPF15 | 1.36 | 0.146 | 2.04 | 0.020 | Mac UP |
| NCAPG | 0.73 | 0.224 | 0.39 | 0.019 | Mac DOWN |
| NCAPG2 | 0.74 | 0.101 | 0.49 | 0.034 | Mac DOWN |
| NCAPH | 0.62 | 0.203 | 0.44 | 0.014 | Mac DOWN |
| NCKAP1 | 1.30 | 0.129 | 2.11 | 0.032 | Mac UP |

| | | | | | |
|---|---|---|---|---|---|
| NDC80 | 0.73 | 0.187 | 0.34 | 0.024 | Mac DOWN |
| NEIL3 | 0.70 | 0.506 | 0.45 | 0.022 | Mac DOWN |
| NEK2 | 1.34 | 0.199 | 0.47 | 0.015 | Mac DOWN |
| NET1 | 0.95 | 0.778 | 0.39 | 0.027 | Mac DOWN |
| NEXN | 0.94 | 0.547 | 0.32 | 0.046 | Mac DOWN |
| NFE2L1 | 1.34 | 0.105 | 2.03 | 0.033 | Mac UP |
| NLRC4 | 3.45 | 0.564 | 2.26 | 0.033 | Mac UP |
| NLRP12 | 12.46 | 0.298 | 5.73 | 0.045 | Mac UP |
| NLRP3 | 0.93 | 0.526 | 0.33 | 0.037 | Mac DOWN |
| NMRK1 | 1.36 | 0.138 | 2.03 | 0.030 | Mac UP |
| NOXA1 | 1.71 | 0.114 | 2.92 | 0.018 | Mac UP |
| NPAS2 | 1.90 | 0.072 | 3.17 | 0.035 | Mac UP |
| NPFFR2 | 1.27 | 0.085 | 5.75 | 0.042 | Mac UP |
| NPIPB2 | 1.39 | 0.470 | 2.99 | 0.036 | Mac UP |
| NPL | 2.73 | 0.479 | 2.52 | 0.020 | Mac UP |
| NR1D2 | 1.35 | 0.197 | 2.03 | 0.032 | Mac UP |
| NR4A3 | 1.84 | 0.095 | 2.23 | 0.013 | Mac UP |
| NSMAF | 1.48 | 0.050 | 2.65 | 0.018 | Mac UP |
| NTF4 | 0.40 | 0.047 | 0.91 | 0.976 | LC DOWN |
| NUF2 | 0.79 | 0.464 | 0.34 | 0.020 | Mac DOWN |
| NUPR1 | 1.67 | 0.543 | 3.04 | 0.018 | Mac UP |
| NUSAP1 | 0.69 | 0.154 | 0.30 | 0.020 | Mac DOWN |
| OAS1 | 0.17 | 0.018 | 0.10 | 0.050 | LC DOWN |
| OAS2 | 0.08 | 0.032 | 0.03 | 0.096 | LC DOWN |
| OASL | 0.17 | 0.074 | 0.11 | 0.016 | Mac DOWN |
| OLFML3 | 0.29 | 0.117 | 0.24 | 0.039 | Mac DOWN |
| OR2A7 | 1.22 | 0.519 | 3.56 | 0.020 | Mac UP |
| ORC1 | 0.47 | 0.084 | 0.32 | 0.048 | Mac DOWN |
| P2RX4 | 1.62 | 0.086 | 2.27 | 0.012 | Mac UP |
| P2RY8_2 | 2.23 | 0.043 | 0.64 | 0.629 | LC UP |
| PACERR | 3.27 | 0.472 | 0.32 | 0.013 | Mac DOWN |
| PAG1 | 1.02 | 0.953 | 2.03 | 0.033 | Mac UP |
| PAPPA2 | 3.78 | 0.037 | 12.31 | 0.011 | Common UP |
| PAQR6 | 1.36 | 0.432 | 2.55 | 0.034 | Mac UP |
| PARD6G-AS1 | 0.43 | 0.042 | 1.81 | 0.516 | LC DOWN |
| PARP6 | 1.21 | 0.122 | 2.33 | 0.014 | Mac UP |
| PARP9 | 0.33 | 0.046 | 0.36 | 0.041 | Common DOWN |
| PBK | 0.66 | 0.151 | 0.32 | 0.048 | Mac DOWN |
| PCDH1 | 2.63 | 0.412 | 2.18 | 0.033 | Mac UP |
| PCDHAC2 | 1.52 | 0.836 | 3.42 | 0.040 | Mac UP |
| PCK2 | 1.41 | 0.085 | 2.76 | 0.018 | Mac UP |
| PCLAF | 0.67 | 0.179 | 0.31 | 0.045 | Mac DOWN |
| PCNA | 0.74 | 0.110 | 0.41 | 0.018 | Mac DOWN |
| PDCD1LG2 | 0.70 | 0.108 | 0.45 | 0.027 | Mac DOWN |
| PDE4B | 0.93 | 0.723 | 0.41 | 0.018 | Mac DOWN |
| PDE8A | 1.15 | 0.329 | 2.13 | 0.029 | Mac UP |
| PDK4 | 5.25 | 0.322 | 3.05 | 0.031 | Mac UP |
| PER3 | 1.56 | 0.136 | 2.76 | 0.045 | Mac UP |
| PGGHG | 0.97 | 0.808 | 2.07 | 0.027 | Mac UP |
| PHF1 | 1.21 | 0.236 | 2.16 | 0.025 | Mac UP |
| PHGDH | 1.95 | 0.036 | 5.96 | 0.012 | Mac UP |
| PIK3CD | 1.23 | 0.342 | 0.45 | 0.037 | Mac DOWN |
| PIK3R6 | 0.00 | 0.615 | 0.47 | 0.044 | Mac DOWN |
| PKD1P6-NPIPP1 | 1.07 | 0.875 | 2.14 | 0.039 | Mac UP |
| PKMYT1 | 0.53 | 0.208 | 0.45 | 0.013 | Mac DOWN |
| PLCB4 | 0.70 | 0.279 | 2.08 | 0.015 | Mac UP |
| PLIN2 | 1.46 | 0.194 | 2.13 | 0.020 | Mac UP |
| PLK1 | 0.99 | 0.947 | 0.46 | 0.018 | Mac DOWN |
| PLK4 | 0.94 | 0.714 | 0.40 | 0.039 | Mac DOWN |
| PLSCR1 | 0.42 | 0.033 | 0.45 | 0.087 | LC DOWN |

| | | | | | |
|---|---|---|---|---|---|
| PNPLA1 | 3.73 | 0.567 | **2.52** | **0.035** | Mac UP |
| PPIL6 | 3.22 | 0.154 | **3.08** | **0.011** | Mac UP |
| PRC1 | 0.88 | 0.440 | **0.34** | **0.014** | Mac DOWN |
| PRKACB | 1.40 | 0.151 | **2.08** | **0.018** | Mac UP |
| PROCR | 0.71 | 0.057 | **0.35** | **0.045** | Mac DOWN |
| PRSS23 | **0.39** | **0.029** | 0.26 | 0.078 | LC DOWN |
| PRSS27 | 1.21 | 0.668 | **2.35** | **0.023** | Mac UP |
| PRSS36 | **0.39** | **0.023** | 0.62 | 0.216 | LC DOWN |
| PRSS8 | **0.30** | **0.023** | 0.62 | 0.590 | LC DOWN |
| PSAT1 | **2.07** | **0.044** | **9.33** | **0.020** | Common UP |
| PSRC1 | 0.88 | 0.692 | **0.48** | **0.043** | Mac DOWN |
| PTER | 1.17 | 0.671 | **2.99** | **0.012** | Mac UP |
| PTTG1 | 0.90 | 0.278 | **0.34** | **0.024** | Mac DOWN |
| PYCR1 | 2.60 | 0.283 | **2.23** | **0.049** | Mac UP |
| PYGO1 | 1.46 | 0.197 | **2.04** | **0.015** | Mac UP |
| RAB3IP | 1.28 | 0.079 | **2.39** | **0.032** | Mac UP |
| RACGAP1 | 1.02 | 0.791 | **0.48** | **0.016** | Mac DOWN |
| RAD51 | 0.75 | 0.539 | **0.46** | **0.032** | Mac DOWN |
| RAD54B | 0.91 | 0.696 | **0.42** | **0.026** | Mac DOWN |
| RAD54L | 0.47 | 0.136 | **0.44** | **0.022** | Mac DOWN |
| RASGRP3 | 0.85 | 0.818 | **0.36** | **0.017** | Mac DOWN |
| RCC1 | 0.89 | 0.313 | **0.45** | **0.018** | Mac DOWN |
| RELN | **2.09** | **0.044** | 0.87 | 0.917 | LC UP |
| RFFL | 1.06 | 0.299 | **2.02** | **0.022** | Mac UP |
| RFNG | 1.21 | 0.203 | **2.18** | **0.030** | Mac UP |
| RGS1 | 0.38 | 0.469 | **0.33** | **0.033** | Mac DOWN |
| RGS3 | 0.66 | 0.094 | **0.48** | **0.018** | Mac DOWN |
| RILPL1 | 1.08 | 0.451 | **2.00** | **0.027** | Mac UP |
| RMI2 | 0.47 | 0.119 | **0.42** | **0.019** | Mac DOWN |
| RNASE2 | 1358.45[3)] | 0.575 | **0.16** | **0.019** | Mac DOWN |
| RNF170 | 1.26 | 0.518 | **2.03** | **0.034** | Mac UP |
| RNF187 | 1.04 | 0.377 | **2.39** | **0.020** | Mac UP |
| ROPN1 | 0.00 | 0.769 | **685.67[3)]** | **0.033** | Mac UP |
| RPS6KA2 | 3.10 | 0.134 | **4.30** | **0.020** | Mac UP |
| RRM2 | **0.43** | **0.020** | 0.24 | 0.053 | LC DOWN |
| RTL3 | 0.00 | 0.630 | **0.26** | **0.034** | Mac DOWN |
| RUFY3 | 1.38 | 0.266 | **2.14** | **0.027** | Mac UP |
| S100A10 | 0.60 | 0.045 | **0.45** | **0.018** | Mac DOWN |
| S100A12 | 0.00 | 0.629 | **0.44** | **0.025** | Mac DOWN |
| S100A4 | 0.39 | 0.219 | **0.28** | **0.030** | Mac DOWN |
| SAMHD1 | **0.44** | **0.022** | 0.51 | 0.046 | LC DOWN |
| SARS1 | 1.65 | 0.021 | **2.12** | **0.040** | Mac UP |
| SBSN | 1.74 | 0.047 | **0.28** | **0.016** | Mac DOWN |
| SCD | 0.25 | 0.051 | **0.38** | **0.016** | Mac DOWN |
| SDSL | 1.61 | 0.212 | **2.38** | **0.021** | Mac UP |
| SEL1L3 | 1.17 | 0.186 | **2.05** | **0.018** | Mac UP |
| SENP6 | 1.33 | 0.118 | **2.10** | **0.021** | Mac UP |
| SERPINA1 | 1.25 | 0.102 | **0.38** | **0.031** | Mac DOWN |
| SERPINB2 | **3.52** | **0.039** | 1.59 | 0.434 | LC UP |
| SERPINB7 | 1.76 | 0.104 | **3.25** | **0.046** | Mac UP |
| SESN2 | 1.90 | 0.070 | **3.25** | **0.026** | Mac UP |
| SETD9 | 1.33 | 0.561 | **2.11** | **0.042** | Mac UP |
| SGIP1 | 0.84 | 0.347 | **0.42** | **0.050** | Mac DOWN |
| SH2D2A | 0.80 | 0.334 | **0.42** | **0.042** | Mac DOWN |
| SH3GLB1 | 1.08 | 0.359 | **2.28** | **0.015** | Mac UP |
| SHCBP1 | 0.64 | 0.256 | **0.33** | **0.023** | Mac DOWN |
| SHROOM2 | 0.51 | 0.769 | **0.11** | **0.014** | Mac DOWN |
| SIGLEC1 | 0.10 | 0.312 | **0.03** | **0.014** | Mac DOWN |
| SIGLEC15 | 0.95 | 0.229 | **0.32** | **0.044** | Mac DOWN |
| SIPA1L2 | 1.60 | 0.128 | **2.30** | **0.012** | Mac UP |

| | | | | | |
|---|---|---|---|---|---|
| SIRPB2 | **0.10** | **0.016** | 0.37 | 0.112 | LC DOWN |
| SKA2 | 0.88 | 0.122 | **0.42** | **0.016** | Mac DOWN |
| SLAMF9 | 0.65 | 0.553 | **0.33** | **0.013** | Mac DOWN |
| SLC1A4 | 1.59 | 0.102 | **2.06** | **0.015** | Mac UP |
| SLC1A5 | 1.39 | 0.062 | **2.54** | **0.012** | Mac UP |
| SLC22A15 | 2.47 | 0.209 | **2.69** | **0.020** | Mac UP |
| SLC25A19 | 0.92 | 0.551 | **0.44** | **0.022** | Mac DOWN |
| SLC25A34 | 0.90 | 0.755 | **2.19** | **0.028** | Mac UP |
| SLC2A10 | 0.59 | 0.768 | **2.77** | **0.032** | Mac UP |
| SLC30A1 | 2.15 | 0.087 | **2.81** | **0.024** | Mac UP |
| SLC38A1 | 1.34 | 0.125 | **2.74** | **0.016** | Mac UP |
| SLC38A6 | 1.28 | 0.516 | **2.64** | **0.029** | Mac UP |
| SLC39A10 | 0.42 | 0.071 | **0.27** | **0.026** | Mac DOWN |
| SLC41A2 | 1.23 | 0.233 | **2.45** | **0.036** | Mac UP |
| SLC48A1 | 1.19 | 0.133 | **2.16** | **0.041** | Mac UP |
| SLC7A10 | | | **2.32** | **0.034** | Mac UP |
| SLC7A11 | 3.00 | 0.132 | **4.64** | **0.029** | Mac UP |
| SLC7A2 | **3.52** | **0.044** | 1.89 | 0.096 | LC UP |
| SLFN5 | 0.98 | 0.884 | **2.49** | **0.014** | Mac UP |
| SLIT2 | **2.14** | **0.044** | 0.46 | 0.551 | LC UP |
| SLITRK6 | | | **6.36** | **0.040** | Mac UP |
| SMARCD3 | 1.18 | 0.630 | **2.23** | **0.038** | Mac UP |
| SMPDL3A | 1.01 | 0.971 | **0.33** | **0.015** | Mac DOWN |
| SNHG12 | **0.50** | **0.038** | 1.44 | 0.376 | LC DOWN |
| SNHG7 | 1.23 | 0.197 | **2.36** | **0.022** | Mac UP |
| SNN | 1.56 | 0.136 | **2.53** | **0.046** | Mac UP |
| SNRNP25 | 0.70 | 0.147 | **0.45** | **0.033** | Mac DOWN |
| SNX20 | 1.04 | 0.962 | **0.48** | **0.028** | Mac DOWN |
| SOBP | 1.38 | 0.195 | **0.39** | **0.040** | Mac DOWN |
| SP110 | **0.49** | **0.035** | **0.42** | **0.025** | Common DOWN |
| SPAG5 | 0.89 | 0.485 | **0.38** | **0.027** | Mac DOWN |
| SPARC | 0.89 | 0.140 | **0.48** | **0.018** | Mac DOWN |
| SPATA18 | 0.80 | 0.890 | **3.51** | **0.048** | Mac UP |
| SPC24 | 0.33 | 0.116 | **0.33** | **0.021** | Mac DOWN |
| SPC25 | 0.28 | 0.302 | **0.26** | **0.015** | Mac DOWN |
| SPEG | 1.06 | 0.743 | **2.36** | **0.029** | Mac UP |
| SPIRE1 | 1.37 | 0.059 | **2.17** | **0.015** | Mac UP |
| SPOPL-DT | 1.11 | 0.752 | **6.40** | **0.039** | Mac UP |
| SPP1 | 1.02 | 0.938 | **0.42** | **0.024** | Mac DOWN |
| SQSTM1 | 1.60 | 0.045 | **3.30** | **0.013** | Mac UP |
| SREBF1 | **0.33** | **0.021** | **0.34** | **0.048** | Common DOWN |
| SRGN | 0.85 | 0.490 | **0.48** | **0.021** | Mac DOWN |
| STAR | 0.56 | 0.734 | **0.21** | **0.049** | Mac DOWN |
| STARD7-AS1 | 1.08 | 0.760 | **2.38** | **0.024** | Mac UP |
| STAT1 | **0.44** | **0.047** | 0.90 | 0.664 | LC DOWN |
| STC2 | 0.77 | 0.138 | **2.11** | **0.040** | Mac UP |
| STK38L | 0.78 | 0.197 | **0.35** | **0.040** | Mac DOWN |
| STMN1 | 0.75 | 0.090 | **0.34** | **0.016** | Mac DOWN |
| STRADB | 1.22 | 0.403 | **2.05** | **0.029** | Mac UP |
| STRBP | 1.26 | 0.172 | **2.04** | **0.044** | Mac UP |
| STRC | 2.43 | 0.352 | **2.68** | **0.038** | Mac UP |
| SUSD5 | 0.96 | 0.946 | **2.23** | **0.012** | Mac UP |
| SYBU | 1.96 | 0.256 | **12.59** | **0.033** | Mac UP |
| SYCE2 | **2.24** | **0.048** | 0.40 | 0.215 | LC UP |
| SYNE1 | 1.30 | 0.184 | **0.46** | **0.041** | Mac DOWN |
| SYTL1 | 1.78 | 0.667 | **6.71** | **0.020** | Mac UP |
| TACC3 | 0.65 | 0.171 | **0.47** | **0.023** | Mac DOWN |
| TAGAP | 1.20 | 0.872 | **0.45** | **0.027** | Mac DOWN |
| TAL1 | 0.13 | 0.697 | **0.27** | **0.048** | Mac DOWN |
| TAPT1-AS1 | 1.95 | 0.397 | **3.81** | **0.027** | Mac UP |

| | | | | | |
|---|---|---|---|---|---|
| TAS2R4 | 1.71 | 0.651 | 5.94 | 0.049 | Mac UP |
| TBL1X | 1.42 | 0.086 | 3.80 | 0.022 | Mac UP |
| TBX15 | 6.79 | 0.606 | 2.46 | 0.030 | Mac UP |
| TCEA1 | 1.83 | 0.174 | 2.85 | 0.031 | Mac UP |
| TCF19 | 0.50 | 0.073 | 0.37 | 0.012 | Mac DOWN |
| TDRD3 | 1.36 | 0.248 | 2.10 | 0.018 | Mac UP |
| TDRD9 | 2.17 | 0.711 | 0.45 | 0.016 | Mac DOWN |
| TESK1 | 1.26 | 0.108 | 2.10 | 0.033 | Mac UP |
| TFPI | 0.77 | 0.118 | 0.42 | 0.022 | Mac DOWN |
| TGM3 | 0.12 | 0.528 | 0.35 | 0.038 | Mac DOWN |
| THBD | 0.67 | 0.461 | 0.49 | 0.043 | Mac DOWN |
| THNSL2 | 1.17 | 0.921 | 2.38 | 0.015 | Mac UP |
| TIFAB | 0.62 | 0.731 | 0.42 | 0.048 | Mac DOWN |
| TK1 | 0.43 | 0.151 | 0.37 | 0.014 | Mac DOWN |
| TLCD2 | 1.39 | 0.149 | 3.29 | 0.018 | Mac UP |
| TLR8 | 2.33 | 0.570 | 2.18 | 0.046 | Mac UP |
| TLX2 | 501.53[3)] | 0.517 | 1102.44[3)] | 0.033 | Mac UP |
| TM4SF19-DYNLT2B | 2.03 | 0.545 | 7.14 | 0.037 | Mac UP |
| TMEM119 | 0.50 | 0.699 | 0.42 | 0.045 | Mac DOWN |
| TMEM140 | 0.42 | 0.039 | 0.70 | 0.122 | LC DOWN |
| TMEM150A | 1.22 | 0.197 | 2.63 | 0.024 | Mac UP |
| TMEM182 | 1.61 | 0.160 | 2.08 | 0.043 | Mac UP |
| TMEM232 | 1.67 | 0.618 | 3.25 | 0.035 | Mac UP |
| TMEM254-AS1 | 1.63 | 0.333 | 2.66 | 0.033 | Mac UP |
| TMEM42 | 1.29 | 0.176 | 2.12 | 0.031 | Mac UP |
| TMIE | | | 2.88 | 0.050 | Mac UP |
| TNIP3 | 2.14 | 0.375 | 2.83 | 0.039 | Mac UP |
| TNNC1 | 0.61 | 0.753 | 4.89 | 0.028 | Mac UP |
| TP73 | 0.52 | 0.089 | 0.35 | 0.018 | Mac DOWN |
| TPX2 | 0.78 | 0.201 | 0.33 | 0.013 | Mac DOWN |
| TRAF3IP3 | 1.50 | 0.727 | 0.33 | 0.014 | Mac DOWN |
| TRAM2-AS1 | 1.18 | 0.570 | 2.50 | 0.025 | Mac UP |
| TREM1 | 0.17 | 0.291 | 0.18 | 0.013 | Mac DOWN |
| TRIB2 | 1.28 | 0.338 | 0.48 | 0.014 | Mac DOWN |
| TRIM14 | 0.38 | 0.035 | 0.58 | 0.017 | LC DOWN |
| TRIM16L | 1.09 | 0.684 | 3.14 | 0.014 | Mac UP |
| TRIM34 | 0.50 | 0.156 | 0.41 | 0.036 | Mac DOWN |
| TRIP13 | 0.94 | 0.514 | 0.45 | 0.030 | Mac DOWN |
| TROAP | 0.79 | 0.330 | 0.44 | 0.022 | Mac DOWN |
| TSEN15 | 1.31 | 0.120 | 2.09 | 0.020 | Mac UP |
| TSTD3 | 1.01 | 0.949 | 2.04 | 0.016 | Mac UP |
| TTK | 0.94 | 0.721 | 0.38 | 0.032 | Mac DOWN |
| TTLL4 | 1.29 | 0.089 | 2.18 | 0.018 | Mac UP |
| TUBA1A | 0.70 | 0.094 | 0.39 | 0.018 | Mac DOWN |
| TUBA1B | 0.58 | 0.100 | 0.47 | 0.015 | Mac DOWN |
| TUBB3 | 0.60 | 0.048 | 0.41 | 0.018 | Mac DOWN |
| TUBE1 | 1.52 | 0.180 | 2.51 | 0.018 | Mac UP |
| TYMS | 0.53 | 0.023 | 0.28 | 0.014 | Mac DOWN |
| UBE2C | 1.01 | 0.977 | 0.40 | 0.017 | Mac DOWN |
| UBE2T | 0.80 | 0.247 | 0.41 | 0.032 | Mac DOWN |
| UCN2 | 0.72 | 0.158 | 0.31 | 0.037 | Mac DOWN |
| UHRF1 | 0.52 | 0.101 | 0.25 | 0.015 | Mac DOWN |
| ULBP1 | 2.68 | 0.147 | 3.36 | 0.015 | Mac UP |
| UNC13A | 0.89 | 0.601 | 2.26 | 0.024 | Mac UP |
| UNC5B | 6.43 | 0.129 | 5.24 | 0.025 | Mac UP |
| UST-AS2 | 1.04 | 0.959 | 0.31 | 0.038 | Mac DOWN |
| UVSSA | 1.53 | 0.113 | 2.37 | 0.030 | Mac UP |
| VCAM1 | 0.33 | 0.040 | 0.35 | 0.160 | LC DOWN |
| VPS13C | 1.35 | 0.230 | 2.24 | 0.030 | Mac UP |
| VTN | 1.06 | 0.973 | 0.32 | 0.021 | Mac DOWN |

| | | | | | |
|---|---|---|---|---|---|
| VWA7 | 0.69 | 0.498 | **4.24** | **0.043** | Mac UP |
| WARS1 | 1.53 | 0.175 | **2.27** | **0.044** | Mac UP |
| WARS2-AS1 | 1.24 | 0.044 | **2.31** | **0.011** | Mac UP |
| WDR31 | 1.44 | 0.042 | **2.08** | **0.049** | Mac UP |
| WDR76 | 0.84 | 0.148 | **0.36** | **0.024** | Mac DOWN |
| WIPI1 | 1.72 | 0.071 | **2.02** | **0.031** | Mac UP |
| XAF1 | **0.21** | **0.034** | **0.41** | **0.034** | Common DOWN |
| XPOT | 1.70 | 0.054 | **3.05** | **0.024** | Mac UP |
| ZBED3 | 1.31 | 0.591 | **3.18** | **0.024** | Mac UP |
| ZDHHC1 | 1.45 | 0.737 | **2.28** | **0.024** | Mac UP |
| ZFAND3 | 1.22 | 0.134 | **2.16** | **0.037** | Mac UP |
| ZFP41 | 1.34 | 0.253 | **2.22** | **0.046** | Mac UP |
| ZNF181 | 1.82 | 0.181 | **2.35** | **0.020** | Mac UP |
| ZNF22-AS1 | 1.15 | 0.962 | **2.49** | **0.033** | Mac UP |
| ZNF252P | 1.34 | 0.147 | **2.17** | **0.015** | Mac UP |
| ZNF367 | 0.61 | 0.322 | **0.38** | **0.019** | Mac DOWN |
| ZNF385C | 2.90 | 0.087 | **5.93** | **0.020** | Mac UP |
| ZNF425 | 1.66 | 0.445 | **2.34** | **0.028** | Mac UP |
| ZNF853 | 1.82 | 0.485 | **2.23** | **0.014** | Mac UP |
| ZWINT | 0.41 | 0.151 | **0.40** | **0.012** | Mac DOWN |

The numbers in blue, red and purple indicate |log2FC| > 1 and Padj < 0.05 in HLF cells, macrophages and common, respectively, where FC and Padj mean fold change and adjusted P value, respectively.

1) Fold changes were calculated as the ratio of mean expression levels (TPM) in HX531 treatment group to those in control group.

2) Blank indicates no expression.

3) Zero TPM in control group was replaced with 0.0001, and then fold changes were calculated.

**Supplementary Table 5. Pathways predicted by GoTerm Biological Process**

**Pathways enriched from DEGs in HLF cells**

| Term | Count | Fold Enric | FDR | Genes |
|---|---|---|---|---|
| GO:0051607~defense response to virus | 15 | 22.9 | 0.000 | IFITM3, STAT1, MX1, IFIT5, ISG15, PARP9, SAMHD1, IFI44L, IFIH1, ISG20, HERC5, PLSCR1, OAS1, OAS2, BCL2L1 |
| GO:0045071~negative regulation of viral genome replication | 9 | 72.0 | 0.000 | ISG20, IFIH1, IFITM3, PLSCR1, OAS1, OAS2, MX1, IFIT5, ISG15 |
| GO:0009615~response to virus | 9 | 29.5 | 0.000 | ISG20, IFIH1, IFITM3, OAS1, OAS2, MX1, IFI44, ISG15, BCL2L1 |
| GO:0070106~interleukin-27-mediated signaling pathway | 4 | 205.9 | 0.000 | OAS1, STAT1, OAS2, MX1 |
| GO:0045087~innate immune response | 12 | 6.8 | 0.000 | ISG20, IFIH1, HERC5, OAS1, OAS2, MX1, IFIT5, TRIM14, ISG15, IFI35, PARP9, SAMHD1 |
| GO:0035456~response to interferon-beta | 4 | 131.0 | 0.000 | IFITM3, PLSCR1, STAT1, XAF1 |
| GO:0060337~type I interferon signaling pathway | 5 | 32.2 | 0.001 | IFIH1, IFITM3, OAS1, STAT1, OAS2 |
| GO:0032728~positive regulation of interferon-beta production | 4 | 33.5 | 0.015 | IFIH1, OAS1, OAS2, ISG15 |
| GO:0060339~negative regulation of type I interferon-mediated signaling pathway | 3 | 49.1 | 0.099 | OAS1, ISG15, SAMHD1 |
| GO:0034341~response to interferon-gamma | 3 | 40.0 | 0.135 | IFITM3, CD74, STAT1 |
| GO:0006508~proteolysis | 6 | 4.7 | 0.402 | RELN, LAP3, PRSS36, PRSS8, PRSS23, PAPPA2 |
| GO:0060700~regulation of ribonuclease activity | 2 | 144.1 | 0.622 | OAS1, OAS2 |
| GO:0032020~ISG15-protein conjugation | 2 | 120.1 | 0.688 | HERC5, ISG15 |
| GO:0034340~response to type I interferon | 2 | 72.0 | 0.993 | MX1, ISG15 |
| GO:0098586~cellular response to virus | 3 | 11.4 | 0.993 | IFIH1, CXCL10, OAS1 |
| GO:0032760~positive regulation of tumor necrosis factor production | 3 | 10.0 | 0.993 | IFIH1, OAS1, OAS2 |
| GO:0010818~T cell chemotaxis | 2 | 48.0 | 0.993 | CXCL10, CXCL11 |
| GO:0032897~negative regulation of viral transcription | 2 | 45.0 | 0.993 | IFITM3, TRIM14 |
| GO:0060384~innervation | 2 | 42.4 | 0.993 | NTF4, VCAM1 |
| GO:0006955~immune response | 5 | 3.5 | 0.993 | IFITM3, CD74, IFI44, SAMHD1, IFI44L |
| GO:0051259~protein oligomerization | 2 | 34.3 | 0.993 | IFIH1, OAS1 |
| GO:0071398~cellular response to fatty acid | 2 | 31.3 | 0.993 | SREBF1, E2F1 |
| GO:0046597~negative regulation of viral entry into host cell | 2 | 30.0 | 0.993 | IFITM3, CD74 |
| GO:0032727~positive regulation of interferon-alpha production | 2 | 27.7 | 0.993 | IFIH1, STAT1 |
| GO:0035458~cellular response to interferon-beta | 2 | 25.7 | 0.993 | OAS1, STAT1 |
| GO:0007214~gamma-aminobutyric acid signaling pathway | 2 | 24.8 | 0.993 | GPR156, GABRG1 |
| GO:0051281~positive regulation of release of sequestered calcium ion into cytosol | 2 | 23.2 | 0.993 | CXCL10, CXCL11 |
| GO:0042127~regulation of cell proliferation | 3 | 6.2 | 0.993 | CXCL10, CXCL11, STAT1 |
| GO:0045664~regulation of neuron differentiation | 2 | 22.5 | 0.993 | NTF4, RELN |
| GO:0045089~positive regulation of innate immune response | 2 | 22.5 | 0.993 | PLSCR1, IFI35 |
| GO:0008283~cell proliferation | 3 | 6.1 | 0.993 | CD74, MAB21L2, BCL2L1 |
| GO:0008015~blood circulation | 2 | 21.2 | 0.993 | CXCL10, STAT1 |
| GO:0006401~RNA catabolic process | 2 | 21.2 | 0.993 | ISG20, OAS2 |
| GO:0051085~chaperone mediated protein folding requiring cofactor | 2 | 21.2 | 0.993 | CD74, HSPA6 |
| GO:0002230~positive regulation of defense response to virus by host | 2 | 20.6 | 0.993 | STAT1, PARP9 |
| GO:0045648~positive regulation of erythrocyte differentiation | 2 | 20.6 | 0.993 | STAT1, ISG15 |
| GO:0000077~DNA damage checkpoint | 2 | 19.5 | 0.993 | E2F1, PARP9 |
| GO:0007616~long-term memory | 2 | 19.0 | 0.993 | NTF4, RELN |

**Pathways enriched from DEGs in macrophages**

| Term | Count | Fold Enric | FDR | Genes |
|---|---|---|---|---|

| | | | | |
|---|---|---|---|---|
| GO:0051301~cell division | 51 | 5.3 | 0.000 | GPSM2, ERCC6L, CCNF, NCAPG2, BUB1B, CDC20, TUBA1B, TUBA1A, PTTG1, NUF2, RCC1, NEK2, FBXO5, HELLS, KNL1, CDC25C, CDC25A, CCNA2, PSRC1, CCNE2, KIFC1, CKS2, BIRC5, MCM5, KIF2C, CDCA3, CDCA5, NCAPG, CDCA8, NCAPH, AURKB, AURKA, SKA2, CCNB2, CCNB1, CDT1, CENPW, SPAG5, UBE2C, CDC7, NDC80, ZWINT, CENPE, TPX2, KIF18B, PRC1, CDK1, TACC3, SPC24, SPC25, MAD2L1 |
| GO:0007052~mitotic spindle organization | 18 | 12.3 | 0.000 | GPSM2, PLK1, CDCA8, TTK, NDC80, AURKB, AURKA, CENPE, CCNB1, INCENP, KIF4A, NUF2, STMN1, TACC3, BIRC5, RCC1, DLGAP5, SPC25 |
| GO:0000070~mitotic sister chromatid segregation | 15 | 15.7 | 0.000 | SPAG5, PLK1, NCAPG2, CDCA8, KNL1, NDC80, ZWINT, KIF18A, KIF18B, ESPL1, INCENP, KIFC1, NUSAP1, NEK2, MAD2L1 |
| GO:0000278~mitotic cell cycle | 25 | 6.5 | 0.000 | CDCA5, CDCA8, PKMYT1, AURKB, AURKA, SKA2, KIF15, TUBA1B, TUBA1A, TUBB3, MYB, PBK, NEK2, MYBL2, PLK4, CDT1, CENPW, PLK1, TUBE1, NDC80, CENPE, TPX2, KIF18B, INCENP, BIRC5 |
| GO:0007094~mitotic spindle assembly checkpoint | 13 | 16.2 | 0.000 | PLK1, BUB1B, TTK, NDC80, ZWINT, AURKB, CDC20, NUF2, BIRC5, TRIP13, SPC24, MAD2L1, SPC25 |
| GO:0007059~chromosome segregation | 18 | 7.4 | 0.000 | CDT1, CENPW, SPAG5, HJURP, TTK, ESCO2, NDC80, SKA2, CENPE, INCENP, NUF2, BIRC5, RCC1, NEK2, CENPQ, SPC24, DLGAP5, SPC25 |
| GO:0051256~mitotic spindle midzone assembly | 8 | 28.9 | 0.000 | RACGAP1, INCENP, PRC1, KIF4A, CDCA8, BIRC5, KIF23, AURKB |
| GO:0000281~mitotic cytokinesis | 14 | 8.2 | 0.000 | PLK1, CDCA8, KIF23, CENPA, AURKB, ANLN, ESPL1, RACGAP1, INCENP, KIF4A, STMN1, NUSAP1, BIRC5, KIF20A |
| GO:0006268~DNA unwinding involved in DNA replication | 9 | 16.3 | 0.000 | GINS1, GINS2, BLM, RAD51, CDC45, MCM3, MCM4, MCM5, MCM2 |
| GO:0000727~double-strand break repair via break-induced replication | 7 | 23.2 | 0.000 | GINS2, CDC45, MCM3, MCM4, CDC7, MCM5, MCM2 |
| GO:0000079~regulation of cyclin-dependent protein serine/threonine kinase activity | 11 | 8.1 | 0.000 | CDKN2D, CCNA2, CCNB2, BLM, CCNB1, CCNE2, CCNF, PKMYT1, CDC25C, CDC25A, CDKN3 |
| GO:0000086~G2/M transition of mitotic cell cycle | 11 | 7.5 | 0.000 | CCNA2, CCNB1, MELK, PLK1, CDK1, BIRC5, PKMYT1, FOXM1, CDC25C, CDC25A, AURKA |
| GO:0007049~cell cycle | 27 | 2.9 | 0.000 | ERCC6L, CDCA3, UHRF1, CDCA5, HJURP, CDCA8, AURKA, CDC45, NUF2, E2F2, CDKN2D, FANCI, CDT1, HELLS, STRADB, PLK1, CDC25C, CDC25A, CCPG1, MCM3, BIRC5, MCM4, MCM5, SPC24, CDKN3, MCM2, TP73 |
| GO:0010971~positive regulation of G2/M transition of mitotic cell cycle | 8 | 11.0 | 0.001 | CCNB1, SMARCD3, CDK1, CDC7, FBXO5, CDC25C, DTL, CDC25A |
| GO:0090267~positive regulation of mitotic cell cycle spindle assembly checkpoint | 6 | 19.9 | 0.001 | INCENP, CDCA8, BIRC5, NDC80, AURKB, MAD2L1 |
| GO:0007018~microtubule-based movement | 12 | 5.4 | 0.002 | DNAI4, CENPE, KIF18A, KIF18B, KIFC1, KIF4A, KIF23, KIF2C, KIF20A, KIF3C, KIF27, KIF15 |
| GO:0030174~regulation of DNA-dependent DNA replication initiation | 6 | 17.0 | 0.002 | CDT1, GMNN, MCM3, MCM4, MCM5, MCM2 |
| GO:0006270~DNA replication initiation | 7 | 9.9 | 0.007 | CDC45, ORC1, CCNE2, MCM3, MCM4, MCM5, MCM2 |
| GO:0006260~DNA replication | 13 | 4.2 | 0.007 | GINS2, BLM, FEN1, RMI2, PCNA, DSCC1, ORC1, PCLAF, CDK1, MCM4, MCM5, DTL, MCM2 |
| GO:0007057~spindle assembly involved in female meiosis I | 4 | 39.8 | 0.007 | CCNB2, FBXO5, NDC80, AURKA |
| GO:1901970~positive regulation of mitotic sister chromatid separation | 4 | 39.8 | 0.007 | INCENP, CDCA8, BIRC5, AURKB |
| GO:0008608~attachment of spindle microtubules to kinetochore | 6 | 13.3 | 0.007 | NUF2, KNL1, NDC80, SPC24, AURKB, SPC25 |
| GO:0007080~mitotic metaphase plate congression | 8 | 7.6 | 0.008 | CENPE, KIF18A, CCNB1, PSRC1, KIFC1, CDCA5, CDCA8, KIF2C |
| GO:0051315~attachment of mitotic spindle microtubules to kinetochore | 5 | 16.6 | 0.016 | CDT1, CENPE, NUF2, KIF2C, NDC80 |
| GO:1903490~positive regulation of mitotic cytokinesis | 4 | 26.5 | 0.028 | INCENP, CDCA8, BIRC5, AURKB |
| GO:0051310~metaphase plate congression | 5 | 14.2 | 0.028 | CENPE, INCENP, KIF2C, CENPQ, NDC80 |
| GO:0000082~G1/S transition of mitotic cell cycle | 9 | 5.2 | 0.028 | CCNA2, CCNE2, CDK1, RCC1, CDC7, IQGAP3, CDC25A, EZH2, CDKN3 |

| | | | | |
|---|---|---|---|---|
| GO:0044772~mitotic cell cycle phase transition | 6 | 9.5 | 0.029 | CCNA2, CCNB2, CCNB1, CCNE2, CCNF, CKS2 |
| GO:0042773~ATP synthesis coupled electron transport | 4 | 22.7 | 0.042 | MT-ND4L, MT-ND4, MT-ND5, MT-CO2 |
| GO:0032465~regulation of cytokinesis | 7 | 6.6 | 0.046 | SH3GLB1, PLK4, PRC1, PLK1, KIF20A, AURKB, AURKA |
| GO:1902425~positive regulation of attachment of mitotic spindle microtubules to kinetochore | 4 | 19.9 | 0.061 | INCENP, CDCA8, BIRC5, AURKB |
| GO:0007076~mitotic chromosome condensation | 5 | 10.5 | 0.081 | CDCA5, PLK1, NUSAP1, NCAPG, NCAPH |
| GO:0030593~neutrophil chemotaxis | 9 | 4.3 | 0.081 | LGALS3, CCL7, CCL20, ITGA1, CCL4, PDE4B, S100A12, PIK3CD, TREM1 |
| GO:0006868~glutamine transport | 4 | 17.7 | 0.082 | SLC38A1, SLC38A6, SLC1A4, SLC1A5 |
| GO:0001578~microtubule bundle formation | 6 | 7.2 | 0.087 | PSRC1, PRC1, MAP1B, PLK1, CCSER2, KIF20A |
| GO:0007346~regulation of mitotic cell cycle | 8 | 4.7 | 0.093 | CDC20, PLK1, GMNN, CKS2, BIRC5, FBXO5, DLGAP5, TP73 |
| GO:0044267~cellular protein metabolic process | 5 | 9.5 | 0.107 | FBXO27, MMP2, D2HGDH, FBXO44, PAPPA2 |
| GO:0007088~regulation of mitotic nuclear division | 5 | 8.6 | 0.148 | RCC1, NEK2, FBXO5, PKMYT1, CDC25C |
| GO:0030218~erythrocyte differentiation | 7 | 5.1 | 0.150 | RACGAP1, FECH, EPAS1, TAL1, MYB, NCAPG2, SLC1A5 |
| GO:0006418~tRNA aminoacylation for protein translation | 5 | 8.3 | 0.165 | AARS1, EPRS1, MARS1, IARS1, WARS1 |
| GO:0034501~protein localization to kinetochore | 4 | 13.3 | 0.168 | CDK1, TTK, KNL1, AURKB |
| GO:1905342~positive regulation of protein localization to kinetochore | 3 | 29.8 | 0.191 | CDT1, CENPQ, NDC80 |
| GO:0015990~electron transport coupled proton transport | 3 | 29.8 | 0.191 | MT-ND4, MT-ND5, MT-CO1 |
| GO:0000226~microtubule cytoskeleton organization | 11 | 3.0 | 0.191 | TUBA1B, TTLL4, TUBA1A, TUBB3, PRC1, MAP1B, CDK1, TACC3, MAP7, BIRC5, TUBE1 |
| GO:0051444~negative regulation of ubiquitin-protein transferase activity | 4 | 12.2 | 0.191 | CDC20, BUB1B, FBXO5, MAD2L1 |
| GO:1900264~positive regulation of DNA-directed DNA polymerase activity | 4 | 12.2 | 0.191 | GINS1, GINS2, PCNA, DSCC1 |
| GO:0009636~response to toxic substance | 8 | 3.9 | 0.212 | SNN, RAD51, CCL4, CDK1, NUPR1, SLC7A11, TYMS, ATF4 |
| GO:0045842~positive regulation of mitotic metaphase/anaphase transition | 4 | 11.4 | 0.229 | CDC20, ESPL1, UBE2C, DLGAP5 |
| GO:0007165~signal transduction | 49 | 1.5 | 0.250 | ARHGAP11A, IL1RN, ECM1, PIK3CD, RGS3, RGS1, ADORA3, RPS6KA2, NAMPT, TAGAP, STMN1, CAPN5, PDE4B, GRB10, PDE8A, PRKACB, ARAP2, GAB2, THNSL2, NBL1, NSMAF, GRIA2, CHRNA5, EPAS1, SH2D2A, ARHGAP18, FPR3, AKAP5, IQGAP3, RACGAP1, CCL7, OLFML3, CCL4, SPP1, NLRP3, APBB1, PAG1, FAM83A, CCL20, GDF1, CDC7, LILRB2, INHBE, NET1, NLRP12, P2RX4, NR4A3, IL7R, HSPA1A |
| GO:1902975~mitotic DNA replication initiation | 3 | 23.9 | 0.283 | MCM3, MCM4, MCM2 |
| GO:0006275~regulation of DNA replication | 6 | 5.1 | 0.291 | CCNA2, PCNA, DSCC1, GMNN, ESCO2, FBXO5 |
| GO:0009615~response to virus | 9 | 3.3 | 0.296 | FGR, BST2, IFITM1, STMN1, CCL4, IRF7, TLR8, ISG15, OASL |
| GO:0032731~positive regulation of interleukin-1 beta production | 7 | 4.2 | 0.296 | NLRP12, NLRP3, LPL, TLR8, NLRC4, CD36, INAVA |
| GO:0007019~microtubule depolymerization | 4 | 9.9 | 0.296 | KIF18A, KIF18B, STMN1, KIF2C |
| GO:0051382~kinetochore assembly | 4 | 9.9 | 0.296 | CENPE, CENPW, CENPA, DLGAP5 |
| GO:0090102~cochlea development | 5 | 6.4 | 0.304 | CCNA2, TIFAB, SLITRK6, SOBP, MCM2 |
| GO:0031536~positive regulation of exit from mitosis | 3 | 19.9 | 0.366 | UBE2C, CDCA5, BIRC5 |
| GO:0007131~reciprocal meiotic recombination | 5 | 6.0 | 0.367 | RAD51, RAD54L, MND1, TRIP13, RAD54B |
| GO:0019538~protein metabolic process | 4 | 8.8 | 0.388 | HEMK1, MMP2, D2HGDH, PAPPA2 |
| GO:0045931~positive regulation of mitotic cell cycle | 5 | 5.9 | 0.394 | CCNB1, TAL1, ASNS, BIRC5, AURKA |
| GO:0045648~positive regulation of erythrocyte differentiation | 5 | 5.7 | 0.423 | FAM210B, TAL1, ISG15, HSPA1B, HSPA1A |
| GO:0007051~spindle organization | 4 | 8.4 | 0.423 | SPAG5, TTK, AURKB, AURKA |
| GO:0042692~muscle cell differentiation | 4 | 8.4 | 0.423 | SMARCD3, IFRD1, SPEG, SYNE1 |
| GO:0051383~kinetochore organization | 3 | 17.0 | 0.431 | CDT1, NUF2, NDC80 |
| GO:0000132~establishment of mitotic spindle orientation | 5 | 5.5 | 0.431 | GPSM2, PLK1, CENPA, NDC80, MAD2L1 |
| GO:0006882~cellular zinc ion homeostasis | 5 | 5.5 | 0.431 | MT1F, SLC39A10, MT1G, SLC30A1, MT1E |
| GO:0097421~liver regeneration | 5 | 5.5 | 0.431 | VTN, PCNA, TYMS, EZH2, AURKA |

| | | | | |
|---|---|---|---|---|
| GO:2001240~negative regulation of extrinsic apoptotic signaling pathway in absence of ligand | 5 | 5.4 | 0.467 | UNC5B, STRADB, IFI6, HSPA1B, HSPA1A |
| GO:0006915~apoptotic process | 26 | 1.7 | 0.471 | NCKAP1, HIP1, IFI6, BUB1B, NLRC4, AURKA, NLRP3, APBB1, CD14, RFFL, SH3GLB1, SRGN, HELLS, UNC5B, FBXO10, TPX2, NLRP12, MELK, ESPL1, CDK1, BIRC5, BIRC7, XAF1, SQSTM1, MCM2, TP73 |
| GO:0032729~positive regulation of interferon-gamma production | 7 | 3.5 | 0.483 | CEBPG, PDE4B, IRF8, TLR8, ISG15, CD14, HAVCR2 |
| GO:0007017~microtubule-based process | 5 | 5.2 | 0.483 | TUBA1B, TUBA1A, TUBB3, TUBE1, GTSE1 |
| GO:0006954~inflammatory response | 20 | 1.8 | 0.483 | IL1RN, ECM1, CCL20, PIK3CD, FPR3, NLRC4, LYZ, CCL7, ADORA3, TNIP3, CCL4, SPP1, S100A12, TLR8, NLRP3, CD14, HPSE, SIGLEC1, HAVCR2, CMKLR1 |
| GO:0060236~regulation of mitotic spindle organization | 4 | 7.6 | 0.483 | GPSM2, TPX2, PSRC1, TACC3 |
| GO:0002244~hematopoietic progenitor cell differentiation | 7 | 3.5 | 0.487 | DNAI4, PYGO1, ANLN, TIFAB, ESCO2, BVES, HERC6 |
| GO:0015825~L-serine transport | 3 | 14.9 | 0.494 | SLC1A4, SLC1A5, SLC7A10 |
| GO:0045143~homologous chromosome segregation | 3 | 14.9 | 0.494 | PTTG1, ESPL1, PLK1 |
| GO:0007095~mitotic G2 DNA damage checkpoint | 5 | 5.1 | 0.494 | BLM, PLK1, CDK1, CLSPN, DTL |
| GO:0051607~defense response to virus | 13 | 2.2 | 0.506 | IFITM1, IFI6, IFIT5, ISG15, PARP9, RNASE2, OASL, BST2, IRF7, TLR8, NLRP3, IRF9, TRIM34 |
| GO:1900182~positive regulation of protein localization to nucleus | 5 | 5.0 | 0.525 | SESN2, PLK1, CDK1, TESK1, PARP9 |
| GO:0018105~peptidyl-serine phosphorylation | 11 | 2.4 | 0.532 | RPS6KA2, PLK1, MKNK2, PBK, CDK1, STK38L, TTK, CDC7, CLSPN, PRKACB, AURKA |
| GO:0034644~cellular response to UV | 6 | 3.9 | 0.535 | PCNA, PBK, CDC25A, AURKB, ATF4, AQP1 |
| GO:0090307~mitotic spindle assembly | 5 | 4.9 | 0.545 | CDC20, TPX2, KIFC1, MYBL2, NEK2 |
| GO:0006974~cellular response to DNA damage stimulus | 15 | 2.0 | 0.545 | BLM, UHRF1, PHF1, WDR76, NPAS2, RAD51, UBE2T, PCLAF, IRF7, FLYWCH1, APBB1, FBXO5, SPATA18, DTL, TP73 |
| GO:0007166~cell surface receptor signaling pathway | 16 | 1.9 | 0.552 | CD86, IFITM1, IGSF6, LILRB2, LILRA1, PDCD1LG2, FCRLB, MARCO, FCGR3A, MKNK2, CD14, CD36, IL7R, IRF9, PAPPA2, LY6E |
| GO:0007091~metaphase/anaphase transition of mitotic cell cycle | 3 | 13.3 | 0.552 | PLK1, TACC3, BUB1B |
| GO:1905821~positive regulation of chromosome condensation | 3 | 13.3 | 0.552 | NCAPG2, NCAPG, NCAPH |
| GO:0006865~amino acid transport | 5 | 4.7 | 0.552 | SLC38A1, SLC1A4, SLC7A11, SLC1A5, SLC7A10 |
| GO:0045333~cellular respiration | 5 | 4.7 | 0.552 | NR4A3, MT-CO1, MT-CO2, MT-CO3, MT-CYB |
| GO:0008203~cholesterol metabolic process | 7 | 3.2 | 0.552 | CYP27A1, SREBF1, LIMA1, STAR, FECH, APOC1, NFE2L1 |
| GO:0071294~cellular response to zinc ion | 4 | 6.6 | 0.552 | P2RX4, MT1F, MT1G, MT1E |
| GO:1901673~regulation of mitotic spindle assembly | 4 | 6.6 | 0.552 | PLK1, HSPA1B, KIF15, HSPA1A |
| GO:0048260~positive regulation of receptor-mediated endocytosis | 4 | 6.6 | 0.552 | VTN, HIP1, SGIP1, SYNE1 |
| GO:0034080~CENP-A containing nucleosome assembly | 3 | 11.9 | 0.629 | CENPW, HJURP, CENPA |
| GO:0045071~negative regulation of viral genome replication | 5 | 4.4 | 0.646 | BST2, IFITM1, IFIT5, ISG15, OASL |
| GO:0002548~monocyte chemotaxis | 5 | 4.4 | 0.646 | LGALS3, CCL7, CCL20, CCL4, S100A12 |
| GO:0031214~biomineral tissue development | 4 | 5.9 | 0.683 | SRGN, ECM1, TMEM119, SPP1 |
| GO:0071280~cellular response to copper ion | 4 | 5.9 | 0.683 | MT1F, MT1G, AQP1, MT1E |
| GO:0070301~cellular response to hydrogen peroxide | 6 | 3.5 | 0.683 | NET1, PCNA, MYB, CDK1, EZH2, AQP1 |
| GO:1905820~positive regulation of chromosome separation | 3 | 10.8 | 0.683 | NCAPG2, NCAPG, NCAPH |
| GO:0035456~response to interferon-beta | 3 | 10.8 | 0.683 | BST2, IFITM1, XAF1 |
| GO:1904668~positive regulation of ubiquitin protein ligase activity | 3 | 10.8 | 0.683 | CDC20, UBE2C, PLK1 |
| GO:0051988~regulation of attachment of spindle microtubules to kinetochore | 3 | 10.8 | 0.683 | SPAG5, RACGAP1, NEK2 |
| GO:0000076~DNA replication checkpoint | 3 | 10.8 | 0.683 | CDT1, CDC45, CLSPN |

| | | | | |
|---|---|---|---|---|
| GO:0045087~innate immune response | 25 | 1.6 | 0.699 | ITGAM, IFIT5, IFI6, HMGB3, PIK3CD, IFI35, NLRC4, TREM1, INAVA, OASL, LGALS3, S100A12, NLRP3, CD14, HAVCR2, ISG15, PARP9, FGR, BST2, MARCO, IRF7, SPIRE1, TLR8, TRIM34, RNF187 |
| GO:0007098~centrosome cycle | 5 | 4.1 | 0.713 | TUBA1A, PLK1, CDK1, PCLAF, TUBE1 |
| GO:0097009~energy homeostasis | 5 | 4.1 | 0.713 | NR4A3, SGIP1, NR1D2, CD36, SQSTM1 |
| GO:0045740~positive regulation of DNA replication | 4 | 5.7 | 0.714 | CDT1, PCNA, BRPF3, CDK1 |
| GO:0016310~phosphorylation | 14 | 1.9 | 0.731 | TESK1, PIK3CD, BUB1B, TTK, CDC7, PKMYT1, NMRK1, RPS6KA2, PBK, PDK4, SPEG, IDNK, TK1, ALPK2 |
| GO:0006955~immune response | 21 | 1.6 | 0.731 | CD86, IL1RN, IGSF6, CCL20, CEBPG, IFI6, PIK3CD, LILRB2, PDCD1LG2, PIK3R6, VTN, FCGR3A, RGS1, FTH1, CCL4, IRF8, ACKR3, CD36, IL7R, ULBP1, CMKLR1 |
| GO:0051056~regulation of small GTPase mediated signal transduction | 8 | 2.6 | 0.735 | ARHGAP11A, NET1, RACGAP1, TAGAP, ARAP2, ARHGAP18, SIPA1L2, ARHGEF5 |
| GO:1901224~positive regulation of NIK/NF-kappaB signaling | 6 | 3.3 | 0.736 | CD86, NLRP12, NLRP3, CD14, IFI35, HAVCR2 |
| GO:0034599~cellular response to oxidative stress | 7 | 2.9 | 0.745 | SESN2, PYCR1, SLC7A11, HSPA1B, NFE2L1, ATF4, HSPA1A |
| GO:0051726~regulation of cell cycle | 13 | 1.9 | 0.827 | PLK1, CCNF, FOXM1, CCNE2, FAP, PCLAF, BIRC5, E2F2, BIRC7, DTL, PRKACB, CDKN3, TP73 |
| GO:0071248~cellular response to metal ion | 3 | 9.2 | 0.846 | MT1F, MT1G, MT1E |
| GO:0003333~amino acid transmembrane transport | 4 | 5.0 | 0.915 | SLC38A1, SLC38A6, SLC7A11, SLC7A10 |
| GO:0051276~chromosome organization | 4 | 5.0 | 0.915 | CENPW, PTTG1, RAD54L, CDCA8 |
| GO:0051984~positive regulation of chromosome segregation | 3 | 8.5 | 0.915 | NCAPG2, NCAPG, NCAPH |
| GO:0019985~translesion synthesis | 3 | 8.5 | 0.915 | PCNA, PCLAF, DTL |
| GO:0030195~negative regulation of blood coagulation | 3 | 8.5 | 0.915 | THBD, VTN, TFPI |
| GO:0016567~protein ubiquitination | 20 | 1.6 | 0.915 | MYLIP, DET1, CDCA3, UHRF1, UBE2C, FBXO27, HERC2P3, PLK1, CCNF, FBXO10, FBXO44, CDC20, UVSSA, RNF170, DCAF10, BIRC7, FBXO5, HERC6, RNF187, TRIM34 |
| GO:0009410~response to xenobiotic stimulus | 13 | 1.9 | 0.915 | SREBF1, FECH, MMP2, LPL, TYMS, NPAS2, RAD54B, RAD51, MAP1B, CDK1, RAD54L, PLIN2, TP73 |
| GO:0031334~positive regulation of protein complex assembly | 5 | 3.6 | 0.915 | SH3GLB1, CDT1, LGALS3, TAL1, WARS1 |
| GO:2000104~negative regulation of DNA-dependent DNA replication | 2 | 39.8 | 0.915 | CDT1, GMNN |
| GO:2000521~negative regulation of immunological synapse formation | 2 | 39.8 | 0.915 | LGALS3, HAVCR2 |
| GO:0000022~mitotic spindle elongation | 2 | 39.8 | 0.915 | PRC1, KIF23 |
| GO:2000866~positive regulation of estradiol secretion | 2 | 39.8 | 0.915 | SPP1, CYP19A1 |
| GO:0071163~DNA replication preinitiation complex assembly | 2 | 39.8 | 0.915 | CDT1, GMNN |
| GO:0008315~meiotic G2/MI transition | 2 | 39.8 | 0.915 | CCNB2, NDC80 |
| GO:0043547~positive regulation of GTPase activity | 10 | 2.1 | 0.941 | ARHGAP11A, NET1, CCL7, CCL20, RGS1, CCL4, ARAP2, EZH2, RASGRP3, S100A10 |
| GO:0001932~regulation of protein phosphorylation | 5 | 3.6 | 0.941 | RAD51, SESN2, TLR8, ROPN1, EZH2 |
| GO:0002221~pattern recognition receptor signaling pathway | 3 | 8.0 | 0.952 | NLRP3, NLRC4, INAVA |
| GO:0006636~unsaturated fatty acid biosynthetic process | 3 | 8.0 | 0.952 | FADS2, SCD, ELOVL6 |
| GO:0032436~positive regulation of proteasomal ubiquitin-dependent protein catabolic process | 6 | 2.9 | 0.987 | DET1, PLK1, TRIB2, HSPA1B, AURKA, HSPA1A |
| GO:0001525~angiogenesis | 12 | 1.9 | 0.987 | ECM1, FAP, UNC5B, EPAS1, TAL1, ADM2, SH2D2A, MMP2, ID1, ACKR3, PIK3R6, WARS1 |
| GO:0015909~long-chain fatty acid transport | 3 | 7.5 | 0.987 | FABP5, PLIN2, CD36 |
| GO:0010273~detoxification of copper ion | 3 | 7.5 | 0.987 | MT1F, MT1G, MT1E |
| GO:0035563~positive regulation of chromatin binding | 3 | 7.5 | 0.987 | CDT1, GMNN, PARP9 |
| GO:0030071~regulation of mitotic metaphase/anaphase transition | 4 | 4.4 | 0.987 | CENPE, SMARCD3, UBE2C, PLK1 |
| GO:0071222~cellular response to lipopolysaccharide | 10 | 2.0 | 0.987 | CD86, TNIP3, PDE4B, IRF8, NLRP3, LILRB2, CD14, CD36, PDCD1LG2, HAVCR2 |

| | | | | |
|---|---|---|---|---|
| GO:0071383~cellular response to steroid hormone stimulus | 3 | 7.0 | 0.987 | TFPI, HSPA1B, HSPA1A |
| GO:0043525~positive regulation of neuron apoptotic process | 5 | 3.2 | 0.987 | ITGA1, MYB, MYBL2, NUPR1, ATF4 |
| GO:0042149~cellular response to glucose starvation | 5 | 3.2 | 0.987 | SH3GLB1, SESN2, ASNS, PLIN2, ATF4 |
| GO:2001189~negative regulation of T cell activation via T cell receptor contact with antigen bound to MHC | 2 | 26.5 | 0.987 | LGALS3, HAVCR2 |
| GO:0034421~post-translational protein acetylation | 2 | 26.5 | 0.987 | DSCC1, ESCO2 |
| GO:1902850~microtubule cytoskeleton organization involved in mitosis | 2 | 26.5 | 0.987 | CDK1, TACC3 |
| GO:0009056~catabolic process | 2 | 26.5 | 0.987 | TMEM150A, PTER |
| GO:0032461~positive regulation of protein oligomerization | 2 | 26.5 | 0.987 | ISG15, ZDHHC1 |
| GO:0034371~chylomicron remodeling | 2 | 26.5 | 0.987 | APOC2, LPL |
| GO:0055096~low-density lipoprotein particle mediated signaling | 2 | 26.5 | 0.987 | LPL, CD36 |
| GO:0045926~negative regulation of growth | 3 | 6.6 | 0.987 | MT1F, MT1G, MT1E |
| GO:0071392~cellular response to estradiol stimulus | 4 | 4.0 | 0.987 | FAM210B, CCNA2, KIF18A, MMP2 |
| GO:0006281~DNA repair | 13 | 1.7 | 0.987 | FANCI, BLM, FEN1, RMI2, UHRF1, FOXM1, RAD51, PTTG1, EXO1, UBE2T, CDK1, RAD54L, CLSPN |
| GO:0008283~cell proliferation | 9 | 2.0 | 0.987 | MELK, SH2D2A, CDK1, CKS2, TACC3, CDC25C, CDC25A, AURKB, FAM83A |
| GO:0042776~mitochondrial ATP synthesis coupled proton transport | 5 | 3.1 | 0.987 | MT-ATP6, MT-ND4L, MT-ND4, MT-ND5, MT-ATP8 |
| GO:0042326~negative regulation of phosphorylation | 3 | 6.3 | 0.987 | CDKN2D, GRB10, TESK1 |
| GO:0033280~response to vitamin D | 3 | 6.3 | 0.987 | CDKN2D, STC2, SPP1 |
| GO:0007064~mitotic sister chromatid cohesion | 3 | 6.3 | 0.987 | CDC20, CDCA5, ESCO2 |
| GO:0035094~response to nicotine | 4 | 3.9 | 0.987 | MT-ND4, CHRNA5, MMP2, SLC7A11 |
| GO:0007596~blood coagulation | 6 | 2.6 | 0.987 | PROCR, THBD, SERPINA1, CD36, TFPI, F5 |
| GO:1901796~regulation of signal transduction by p53 class mediator | 4 | 3.8 | 0.987 | SETD9, RFFL, AURKB, AURKA |
| GO:0071168~protein localization to chromatin | 3 | 6.0 | 0.987 | PLK1, ESCO2, EZH2 |
| GO:0015813~L-glutamate transport | 3 | 6.0 | 0.987 | SLC38A6, SLC1A4, SLC7A11 |
| GO:0010165~response to X-ray | 3 | 6.0 | 0.987 | THBD, BLM, RAD51 |
| GO:0009060~aerobic respiration | 5 | 2.9 | 0.987 | MT-ND4L, MT-ND4, MT-ND5, MT-CO1, MT-CO3 |
| GO:0016477~cell migration | 12 | 1.7 | 0.987 | LIMA1, NCKAP1, VTN, DEPDC1B, LAMB3, MMP2, SHROOM2, CDK1, PIK3CD, CTNNA3, CORO7, PARP9 |
| GO:0015804~neutral amino acid transport | 3 | 5.7 | 0.987 | SLC38A1, SLC1A5, SLC7A10 |
| GO:0002526~acute inflammatory response | 3 | 5.7 | 0.987 | NLRP3, NUPR1, TREM1 |
| GO:1905322~positive regulation of mesenchymal stem cell migration | 2 | 19.9 | 0.987 | ACKR3, FBXO5 |
| GO:0000915~actomyosin contractile ring assembly | 2 | 19.9 | 0.987 | ANLN, RACGAP1 |
| GO:0017085~response to insecticide | 2 | 19.9 | 0.987 | FECH, MAP1B |
| GO:0072757~cellular response to camptothecin | 2 | 19.9 | 0.987 | BLM, RAD51 |
| GO:0032375~negative regulation of cholesterol transport | 2 | 19.9 | 0.987 | APOC2, APOC1 |
| GO:1905448~positive regulation of mitochondrial ATP synthesis coupled electron transport | 2 | 19.9 | 0.987 | CCNB1, CDK1 |
| GO:0007079~mitotic chromosome movement towards spindle pole | 2 | 19.9 | 0.987 | CENPE, DLGAP5 |
| GO:0010916~negative regulation of very-low-density lipoprotein particle clearance | 2 | 19.9 | 0.987 | APOC2, APOC1 |
| GO:2000479~regulation of cAMP-dependent protein kinase activity | 2 | 19.9 | 0.987 | NPFFR2, SESN2 |
| GO:0016321~female meiosis chromosome segregation | 2 | 19.9 | 0.987 | PLK1, TTK |
| GO:0070098~chemokine-mediated signaling pathway | 5 | 2.8 | 0.987 | CCL7, CCL20, CCL4, ACKR3, CMKLR1 |

Red indicates FDR < 0.05.

**Supplementary Table 6. Pathways predicted by Reactome**

**Pathways enriched from DEGs in HLF cells**

| Term | Count | Fold Enric | FDR | Genes |
|---|---|---|---|---|
| R-HSA-909733~Interferon alpha/beta signaling | 11 | 36.7 | 0.000 | ISG20, IFITM3, OAS1, STAT1, OAS2, MX1, IFIT5, ISG15, IFI35, XAF1, SAMHD1 |
| R-HSA-913531~Interferon Signaling | 14 | 13.4 | 0.000 | IFITM3, VCAM1, STAT1, MX1, IFIT5, ISG15, IFI35, SAMHD1, ISG20, HERC5, OAS1, OAS2, TRIM14, XAF1 |
| R-HSA-1280215~Cytokine Signaling in Immune system | 17 | 5.5 | 0.000 | IFITM3, VCAM1, SERPINB2, STAT1, MX1, IFIT5, ISG15, IFI35, SAMHD1, ISG20, HERC5, CXCL10, OAS1, OAS2, TRIM14, XAF1, BCL2L1 |
| R-HSA-168256~Immune System | 20 | 2.5 | 0.004 | IFITM3, CD74, VCAM1, SERPINB2, STAT1, HSPA6, MX1, IFIT5, ISG15, IFI35, SAMHD1, IFIH1, ISG20, HERC5, CXCL10, OAS1, OAS2, TRIM14, XAF1, BCL2L1 |
| R-HSA-1169410~Antiviral mechanism by IFN-stimulated genes | 6 | 10.3 | 0.012 | HERC5, OAS1, STAT1, OAS2, MX1, ISG15 |
| R-HSA-877300~Interferon gamma signaling | 5 | 13.4 | 0.020 | VCAM1, OAS1, STAT1, OAS2, TRIM14 |
| R-HSA-1169408~ISG15 antiviral mechanism | 4 | 14.1 | 0.097 | HERC5, STAT1, MX1, ISG15 |
| R-HSA-9833110~RSV-host interactions | 4 | 10.5 | 0.194 | IFIH1, HERC5, OAS2, ISG15 |
| R-HSA-936440~Negative regulators of DDX58/IFIH1 signaling | 3 | 22.0 | 0.221 | IFIH1, HERC5, ISG15 |
| R-HSA-9820952~Respiratory Syncytial Virus Infection Pathway | 4 | 8.6 | 0.270 | IFIH1, HERC5, OAS2, ISG15 |
| R-HSA-9833482~PKR-mediated signaling | 3 | 10.3 | 0.664 | HERC5, STAT1, ISG15 |
| R-HSA-8983711~OAS antiviral response | 2 | 57.1 | 0.664 | OAS1, OAS2 |
| R-HSA-9029558~NR1H2 & NR1H3 regulate gene expression linked to lipc | 2 | 57.1 | 0.664 | SREBF1, FASN |
| R-HSA-168928~DDX58/IFIH1-mediated induction of interferon-alpha/beta | 3 | 9.4 | 0.708 | IFIH1, HERC5, ISG15 |
| R-HSA-6785807~Interleukin-4 and Interleukin-13 signaling | 3 | 7.1 | 1.000 | VCAM1, STAT1, BCL2L1 |
| R-HSA-9679506~SARS-CoV Infections | 5 | 3.1 | 1.000 | IFIH1, STAT1, ISG15, PARP9, BCL2L1 |
| R-HSA-9705671~SARS-CoV-2 activates/modulates innate and adaptive imr | 3 | 6.1 | 1.000 | IFIH1, STAT1, ISG15 |
| R-HSA-9824446~Viral Infection Pathways | 7 | 2.2 | 1.000 | IFIH1, HERC5, STAT1, OAS2, ISG15, PARP9, BCL2L1 |

**Pathways enriched from DEGs in macrophages**

| Term | Count | Fold Enric | FDR | Genes |
|---|---|---|---|---|
| R-HSA-1640170~Cell Cycle | 83 | 3.7 | 0.000 | ERCC6L, FEN1, DSCC1, NCAPG2, GMNN, HJURP, H2AC19, BUB1B, FOXM1, H2AC18, LMNB1, SYNE1, CDC20, TUBA1B, TUBA1A, PTTG1, TUBB3, EXO1, H2BC21, NUF2, RCC1, NEK2, MYBL2, FBXO5, TK1, GTSE1, RMI2, KIF23, ESCO2, KNL1, CDC25C, CDC25A, CCNA2, CCNE2, ESPL1, INCENP, MCM3, BIRC5, MCM4, MCM5, KIF2C, KIF20A, MCM2, BLM, PCNA, CDCA5, NCAPG, CDCA8, HMMR, PKMYT1, TYMS, CENPA, NCAPH, AURKB, AURKA, SKA2, CCNB2, CCNB1, CDC45, ORC1, E2F2, CLSPN, CDKN2D, PLK4, GINS1, GINS2, CDT1, CENPW, UBE2C, PLK1, MND1, CDC7, NDC80, ZWINT, CENPE, TPX2, KIF18A, RAD51, CDK1, CENPQ, SPC24, SPC25, MAD2L1 |

| | | | |
|---|---|---|---|
| R-HSA-69278~Cell Cycle, Mitotic | 73 | 4.0 | 0.000 ERCC6L, FEN1, NCAPG2, GMNN, H2AC19, BUB1B, FOXM1, H2AC18, LMNB1, CDC20, TUBA1B, TUBA1A, PTTG1, TUBB3, H2BC21, NUF2, RCC1, NEK2, MYBL2, FBXO5, TK1, GTSE1, KIF23, ESCO2, KNL1, CDC25C, CDC25A, CCNA2, CCNE2, ESPL1, INCENP, MCM3, BIRC5, MCM4, MCM5, KIF2C, KIF20A, MCM2, PCNA, CDCA5, NCAPG, CDCA8, HMMR, PKMYT1, TYMS, CENPA, NCAPH, AURKB, AURKA, SKA2, CCNB2, CCNB1, CDC45, ORC1, E2F2, CDKN2D, PLK4, GINS1, GINS2, CDT1, UBE2C, PLK1, CDC7, NDC80, ZWINT, CENPE, TPX2, KIF18A, CDK1, CENPQ, SPC24, SPC25, MAD2L1 |
| R-HSA-69620~Cell Cycle Checkpoints | 43 | 4.5 | 0.000 BLM, ERCC6L, CDCA8, BUB1B, PKMYT1, CENPA, AURKB, SKA2, CDC20, CCNB2, CCNB1, CDC45, ORC1, EXO1, H2BC21, NUF2, CLSPN, GTSE1, RMI2, UBE2C, PLK1, CDC7, KNL1, CDC25C, CDC25A, NDC80, ZWINT, CCNA2, CENPE, KIF18A, CCNE2, INCENP, MCM3, CDK1, BIRC5, MCM4, MCM5, KIF2C, CENPQ, SPC24, MCM2, MAD2L1, SPC25 |
| R-HSA-2500257~Resolution of Sister Chromatid Cohesion | 28 | 6.8 | 0.000 ERCC6L, CDCA5, CDCA8, BUB1B, CENPA, AURKB, SKA2, CDC20, CCNB2, TUBA1B, CCNB1, TUBA1A, TUBB3, NUF2, PLK1, KNL1, NDC80, ZWINT, CENPE, KIF18A, INCENP, CDK1, BIRC5, KIF2C, CENPQ, SPC24, MAD2L1, SPC25 |
| R-HSA-68877~Mitotic Prometaphase | 32 | 4.8 | 0.000 ERCC6L, CDCA5, NCAPG, CDCA8, BUB1B, CENPA, NCAPH, AURKB, SKA2, CDC20, CCNB2, TUBA1B, CCNB1, TUBA1A, TUBB3, NUF2, NEK2, PLK4, PLK1, KNL1, NDC80, ZWINT, CENPE, KIF18A, INCENP, CDK1, BIRC5, KIF2C, CENPQ, SPC24, MAD2L1, SPC25 |
| R-HSA-2555396~Mitotic Metaphase and Anaphase | 34 | 4.4 | 0.000 ERCC6L, CDCA5, CDCA8, BUB1B, CENPA, AURKB, SKA2, LMNB1, CDC20, CCNB2, TUBA1B, CCNB1, TUBA1A, PTTG1, TUBB3, NUF2, RCC1, FBXO5, UBE2C, PLK1, KNL1, NDC80, ZWINT, CENPE, KIF18A, ESPL1, INCENP, CDK1, BIRC5, KIF2C, CENPQ, SPC24, MAD2L1, SPC25 |
| R-HSA-9648025~EML4 and NUDC in mitotic spindle formation | 24 | 6.3 | 0.000 ERCC6L, PLK1, CDCA8, BUB1B, KNL1, CENPA, NDC80, ZWINT, AURKB, SKA2, CDC20, CENPE, TUBA1B, KIF18A, TUBA1A, INCENP, TUBB3, NUF2, BIRC5, KIF2C, CENPQ, SPC24, MAD2L1, SPC25 |
| R-HSA-68882~Mitotic Anaphase | 33 | 4.3 | 0.000 ERCC6L, CDCA5, CDCA8, BUB1B, CENPA, AURKB, SKA2, LMNB1, CDC20, CCNB2, TUBA1B, CCNB1, TUBA1A, PTTG1, TUBB3, NUF2, RCC1, UBE2C, PLK1, KNL1, NDC80, ZWINT, CENPE, KIF18A, ESPL1, INCENP, CDK1, BIRC5, KIF2C, CENPQ, SPC24, MAD2L1, SPC25 |
| R-HSA-68886~M Phase | 44 | 3.3 | 0.000 ERCC6L, CDCA5, NCAPG2, H2AC19, NCAPG, CDCA8, BUB1B, CENPA, NCAPH, AURKB, SKA2, H2AC18, LMNB1, CDC20, CCNB2, TUBA1B, CCNB1, TUBA1A, PTTG1, TUBB3, H2BC21, NUF2, RCC1, NEK2, FBXO5, PLK4, UBE2C, PLK1, KIF23, KNL1, NDC80, ZWINT, CENPE, KIF18A, ESPL1, INCENP, CDK1, BIRC5, KIF2C, KIF20A, CENPQ, SPC24, MAD2L1, SPC25 |
| R-HSA-5663220~RHO GTPases Activate Formins | 25 | 5.5 | 0.000 ERCC6L, CDCA8, BUB1B, CENPA, AURKB, SKA2, CDC20, TUBA1B, TUBA1A, TUBB3, NUF2, PLK1, KNL1, NDC80, ZWINT, CENPE, KIF18A, DIAPH3, INCENP, BIRC5, KIF2C, CENPQ, SPC24, MAD2L1, SPC25 |
| R-HSA-141424~Amplification of signal from the kinetochores | 21 | 6.7 | 0.000 ERCC6L, PLK1, CDCA8, BUB1B, KNL1, CENPA, NDC80, ZWINT, AURKB, SKA2, CDC20, CENPE, KIF18A, INCENP, NUF2, BIRC5, KIF2C, CENPQ, SPC24, MAD2L1, SPC25 |

| | | | | |
|---|---|---|---|---|
| R-HSA-141444~Amplification of signal from unattached kinetochores via a | 21 | 6.7 | 0.000 | ERCC6L, PLK1, CDCA8, BUB1B, KNL1, CENPA, NDC80, ZWINT, AURKB, SKA2, CDC20, CENPE, KIF18A, INCENP, NUF2, BIRC5, KIF2C, CENPQ, SPC24, MAD2L1, SPC25 |
| R-HSA-69618~Mitotic Spindle Checkpoint | 22 | 6.0 | 0.000 | ERCC6L, UBE2C, PLK1, CDCA8, BUB1B, KNL1, CENPA, NDC80, ZWINT, AURKB, SKA2, CDC20, CENPE, KIF18A, INCENP, NUF2, BIRC5, KIF2C, CENPQ, SPC24, MAD2L1, SPC25 |
| R-HSA-2467813~Separation of Sister Chromatids | 28 | 4.5 | 0.000 | ERCC6L, CDCA5, CDCA8, BUB1B, CENPA, AURKB, SKA2, CDC20, TUBA1B, TUBA1A, PTTG1, TUBB3, NUF2, UBE2C, PLK1, KNL1, NDC80, ZWINT, CENPE, KIF18A, ESPL1, INCENP, BIRC5, KIF2C, CENPQ, SPC24, MAD2L1, SPC25 |
| R-HSA-983189~Kinesins | 15 | 7.8 | 0.000 | KIF23, KIF27, KIF15, CENPE, TUBA1B, KIF18A, TUBA1A, KIF18B, RACGAP1, TUBB3, KIFC1, KIF4A, KIF2C, KIF20A, KIF3C |
| R-HSA-195258~RHO GTPase Effectors | 34 | 3.2 | 0.000 | NCKAP1, ERCC6L, H2AC19, CDCA8, BUB1B, CENPA, IQGAP3, AURKB, SKA2, H2AC18, CDC20, TUBA1B, TUBA1A, TUBB3, H2BC21, NUF2, PLK1, KNL1, CDC25C, NDC80, ZWINT, CENPE, KIF18A, DIAPH3, PRC1, INCENP, NOXA1, BIRC5, KIF2C, CENPQ, SPC24, ROPN1, MAD2L1, SPC25 |
| R-HSA-453279~Mitotic G1 phase and G1/S transition | 21 | 4.3 | 0.000 | CDKN2D, CDT1, PCNA, GMNN, CDC7, TYMS, CDC25A, CCNA2, CCNB1, CDC45, ORC1, CCNE2, MCM3, CDK1, MCM4, E2F2, MCM5, MYBL2, TK1, FBXO5, MCM2 |
| R-HSA-156711~Polo-like kinase mediated events | 8 | 15.4 | 0.000 | CCNB2, CCNB1, PLK1, MYBL2, PKMYT1, FOXM1, CDC25C, CDC25A |
| R-HSA-69273~Cyclin A/B1/B2 associated events during G2/M transition | 9 | 11.1 | 0.000 | CCNA2, CCNB2, CCNB1, PLK1, CDK1, PKMYT1, FOXM1, CDC25C, CDC25A |
| R-HSA-176974~Unwinding of DNA | 7 | 17.9 | 0.000 | GINS1, GINS2, CDC45, MCM3, MCM4, MCM5, MCM2 |
| R-HSA-69206~G1/S Transition | 18 | 4.2 | 0.000 | CDT1, PCNA, GMNN, CDC7, TYMS, CDC25A, CCNA2, CCNB1, CDC45, ORC1, CCNE2, MCM3, CDK1, MCM4, MCM5, TK1, FBXO5, MCM2 |
| R-HSA-176187~Activation of ATR in response to replication stress | 10 | 8.3 | 0.000 | CDC45, ORC1, MCM3, MCM4, CDC7, MCM5, CLSPN, CDC25C, CDC25A, MCM2 |
| R-HSA-6811434~COPI-dependent Golgi-to-ER retrograde traffic | 15 | 4.7 | 0.000 | KIF23, KIF27, KIF15, CENPE, TUBA1B, KIF18A, TUBA1A, KIF18B, RACGAP1, TUBB3, KIFC1, KIF4A, KIF2C, KIF20A, KIF3C |
| R-HSA-69190~DNA strand elongation | 9 | 8.7 | 0.000 | GINS1, GINS2, FEN1, PCNA, CDC45, MCM3, MCM4, MCM5, MCM2 |
| R-HSA-69481~G2/M Checkpoints | 19 | 3.5 | 0.000 | BLM, RMI2, CDC7, PKMYT1, CDC25C, CDC25A, CCNB2, CCNB1, CDC45, ORC1, EXO1, H2BC21, MCM3, CDK1, MCM4, MCM5, CLSPN, GTSE1, MCM2 |
| R-HSA-68962~Activation of the pre-replicative complex | 9 | 8.4 | 0.000 | CDT1, CDC45, ORC1, GMNN, MCM3, MCM4, CDC7, MCM5, MCM2 |
| R-HSA-194315~Signaling by Rho GTPases | 46 | 2.0 | 0.000 | ARHGAP11A, NCKAP1, ERCC6L, H2AC19, ARHGAP18, CDCA8, BUB1B, SLC1A5, CENPA, IQGAP3, AURKB, SKA2, H2AC18, LMNB1, CDC20, TUBA1B, TUBA1A, RACGAP1, TUBB3, H2BC21, ARHGDIB, NUF2, TAGAP, PLK1, ARAP2, KNL1, CDC25C, NDC80, ZWINT, NET1, CENPE, ANLN, KIF18A, DEPDC1B, DIAPH3, PRC1, INCENP, NOXA1, BIRC5, KIF2C, CENPQ, SPC24, ROPN1, ARHGEF5, MAD2L1, SPC25 |
| R-HSA-176417~Phosphorylation of Emi1 | 5 | 25.6 | 0.001 | CDC20, CCNB1, PLK1, CDK1, FBXO5 |
| R-HSA-9716542~Signaling by Rho GTPases, Miro GTPases and RHOBTI | 46 | 2.0 | 0.001 | ARHGAP11A, NCKAP1, ERCC6L, H2AC19, ARHGAP18, CDCA8, BUB1B, SLC1A5, CENPA, IQGAP3, AURKB, SKA2, H2AC18, LMNB1, CDC20, TUBA1B, TUBA1A, RACGAP1, TUBB3, H2BC21, ARHGDIB, NUF2, TAGAP, PLK1, ARAP2, KNL1, CDC25C, NDC80, ZWINT, NET1, CENPE, ANLN, KIF18A, DEPDC1B, DIAPH3, PRC1, INCENP, NOXA1, BIRC5, KIF2C, CENPQ, SPC24, ROPN1, ARHGEF5, MAD2L1, SPC25 |

| | | | | |
|---|---|---|---|---|
| R-HSA-69242~S Phase | 18 | 3.4 | 0.001 | GINS1, GINS2, CDT1, FEN1, PCNA, UBE2C, CDCA5, GMNN, ESCO2, CDC25A, CCNA2, CDC45, ORC1, CCNE2, MCM3, MCM4, MCM5, MCM2 |
| R-HSA-174143~APC/C-mediated degradation of cell cycle proteins | 13 | 4.6 | 0.001 | UBE2C, PLK1, BUB1B, AURKB, AURKA, CCNA2, CDC20, CCNB1, PTTG1, CDK1, NEK2, FBXO5, MAD2L1 |
| R-HSA-453276~Regulation of mitotic cell cycle | 13 | 4.6 | 0.001 | UBE2C, PLK1, BUB1B, AURKB, AURKA, CCNA2, CDC20, CCNB1, PTTG1, CDK1, NEK2, FBXO5, MAD2L1 |
| R-HSA-69205~G1/S-Specific Transcription | 8 | 8.8 | 0.001 | CDT1, PCNA, CDC45, ORC1, CDK1, TK1, FBXO5, TYMS |
| R-HSA-69239~Synthesis of DNA | 15 | 3.8 | 0.001 | GINS1, GINS2, CDT1, FEN1, PCNA, UBE2C, GMNN, CCNA2, CDC45, ORC1, CCNE2, MCM3, MCM4, MCM5, MCM2 |
| R-HSA-69306~DNA Replication | 19 | 3.1 | 0.001 | GINS1, GINS2, CDT1, FEN1, PCNA, UBE2C, GMNN, H2AC19, CDC7, H2AC18, CCNA2, CDC45, ORC1, CCNE2, H2BC21, MCM3, MCM4, MCM5, MCM2 |
| R-HSA-69275~G2/M Transition | 19 | 3.0 | 0.002 | PLK4, PLK1, HMMR, PKMYT1, FOXM1, CDC25C, CDC25A, AURKA, CCNA2, CCNB2, TPX2, TUBA1B, CCNB1, TUBA1A, TUBB3, CDK1, NEK2, MYBL2, GTSE1 |
| R-HSA-9648895~Response of EIF2AK1 (HRI) to heme deficiency | 6 | 12.3 | 0.002 | CEBPG, GRB10, ASNS, CHAC1, ATF5, ATF4 |
| R-HSA-453274~Mitotic G2-G2/M phases | 19 | 2.9 | 0.002 | PLK4, PLK1, HMMR, PKMYT1, FOXM1, CDC25C, CDC25A, AURKA, CCNA2, CCNB2, TPX2, TUBA1B, CCNB1, TUBA1A, TUBB3, CDK1, NEK2, MYBL2, GTSE1 |
| R-HSA-379716~Cytosolic tRNA aminoacylation | 7 | 9.0 | 0.002 | AARS1, SARS1, EPRS1, MARS1, IARS1, CARS1, WARS1 |
| R-HSA-8856688~Golgi-to-ER retrograde transport | 15 | 3.5 | 0.002 | KIF23, KIF27, KIF15, CENPE, TUBA1B, KIF18A, TUBA1A, KIF18B, RACGAP1, TUBB3, KIFC1, KIF4A, KIF2C, KIF20A, KIF3C |
| R-HSA-983231~Factors involved in megakaryocyte development and platele | 17 | 3.1 | 0.003 | KIF23, KIF27, KIF15, CENPE, TUBA1B, KIF18A, TUBA1A, KIF18B, RACGAP1, TUBB3, KIFC1, KIF4A, MYB, KIF2C, KIF20A, KIF3C, PRKACB |
| R-HSA-909733~Interferon alpha/beta signaling | 11 | 4.4 | 0.004 | BST2, IFITM1, IRF7, IFI6, IFIT5, IRF8, ISG15, IFI35, XAF1, IRF9, OASL |
| R-HSA-109582~Hemostasis | 38 | 1.9 | 0.006 | ECM1, SERPINA1, SPARC, ITGAM, SLC7A11, SLC7A10, TFPI, TREM1, PIK3R6, KIF15, THBD, TUBA1B, TUBA1A, RACGAP1, TUBB3, MYB, KCNMB3, KIF3C, CD36, PRKACB, S100A10, SRGN, BRPF3, ITGA1, KIF23, GTPBP2, KIF27, F5, FGR, CENPE, PROCR, KIF18A, KIF18B, P2RX4, KIFC1, KIF4A, KIF2C, KIF20A |
| R-HSA-2514853~Condensation of Prometaphase Chromosomes | 5 | 14.0 | 0.007 | CCNB2, CCNB1, CDK1, NCAPG, NCAPH |
| R-HSA-69478~G2/M DNA replication checkpoint | 4 | 24.6 | 0.007 | CCNB2, CCNB1, CDK1, PKMYT1 |
| R-HSA-176814~Activation of APC/C and APC/C:Cdc20 mediated degradat | 10 | 4.0 | 0.016 | CCNA2, CDC20, CCNB1, PTTG1, UBE2C, PLK1, CDK1, BUB1B, NEK2, MAD2L1 |
| R-HSA-2980767~Activation of NIMA Kinases NEK9, NEK6, NEK7 | 4 | 17.6 | 0.022 | CCNB2, CCNB1, PLK1, CDK1 |
| R-HSA-1538133~G0 and Early G1 | 6 | 6.8 | 0.031 | CCNA2, PCNA, CCNE2, CDK1, MYBL2, CDC25A |
| R-HSA-3700989~Transcriptional Regulation by TP53 | 24 | 2.0 | 0.032 | FANCI, BLM, RMI2, PCNA, MT-CO1, SETD9, BRPF3, NLRC4, CDC25C, AURKB, AURKA, CCNA2, TPX2, CCNB1, CCNE2, EXO1, SESN2, CDK1, TCEA1, BIRC5, MT-CO2, MT-CO3, RFFL, TP73 |
| R-HSA-73886~Chromosome Maintenance | 13 | 2.9 | 0.038 | BLM, FEN1, PCNA, CENPW, DSCC1, H2AC19, HJURP, KNL1, CENPA, H2AC18, CCNA2, H2BC21, CENPQ |
| R-HSA-2132295~MHC class II antigen presentation | 12 | 3.0 | 0.040 | CENPE, TUBA1B, KIF18A, TUBA1A, RACGAP1, TUBB3, KIF4A, KIF23, KIF2C, KIF20A, KIF3C, KIF15 |
| R-HSA-379724~tRNA Aminoacylation | 7 | 5.1 | 0.040 | AARS1, SARS1, EPRS1, MARS1, IARS1, CARS1, WARS1 |

| | | | | |
|---|---|---|---|---|
| R-HSA-6804114~TP53 Regulates Transcription of Genes Involved in G2 C | 5 | 8.5 | 0.042 | CCNB1, PCNA, CDK1, CDC25C, AURKA |
| R-HSA-6811442~Intra-Golgi and retrograde Golgi-to-ER traffic | 16 | 2.4 | 0.042 | KIF23, KIF27, KIF15, CENPE, TUBA1B, KIF18A, TUBA1A, KIF18B, RACGAP1, MAN2A2, TUBB3, KIFC1, KIF4A, KIF2C, KIF20A, KIF3C |
| R-HSA-69052~Switching of origins to a post-replicative state | 10 | 3.4 | 0.047 | CCNA2, CDT1, ORC1, CCNE2, UBE2C, GMNN, MCM3, MCM4, MCM5, MCM2 |
| R-HSA-176409~APC/C:Cdc20 mediated degradation of mitotic proteins | 9 | 3.7 | 0.049 | CCNA2, CDC20, CCNB1, PTTG1, UBE2C, CDK1, BUB1B, NEK2, MAD2L1 |
| R-HSA-176408~Regulation of APC/C activators between G1/S and early an | 9 | 3.5 | 0.072 | CCNA2, CDC20, CCNB1, UBE2C, PLK1, CDK1, BUB1B, FBXO5, MAD2L1 |
| R-HSA-6791312~TP53 Regulates Transcription of Cell Cycle Genes | 7 | 4.5 | 0.072 | CCNA2, CCNB1, PCNA, CCNE2, CDK1, CDC25C, AURKA |
| R-HSA-69002~DNA Replication Pre-Initiation | 13 | 2.5 | 0.086 | CDT1, UBE2C, GMNN, H2AC19, CDC7, H2AC18, CDC45, ORC1, H2BC21, MCM3, MCM4, MCM5, MCM2 |
| R-HSA-913531~Interferon Signaling | 18 | 2.1 | 0.106 | IFITM1, IFI6, IFIT5, ISG15, IFI35, OASL, BST2, TUBA1B, TUBA1A, TUBB3, IRF7, CDK1, IRF8, XAF1, HSPA1B, IRF9, HSPA1A, TRIM34 |
| R-HSA-179409~APC-Cdc20 mediated degradation of Nek2A | 5 | 6.2 | 0.126 | CDC20, UBE2C, BUB1B, NEK2, MAD2L1 |
| R-HSA-8980692~RHOA GTPase cycle | 12 | 2.5 | 0.128 | ARHGAP11A, NET1, ANLN, DEPDC1B, DIAPH3, RACGAP1, TAGAP, ARHGDIB, ARAP2, ARHGAP18, IQGAP3, ARHGEF5 |
| R-HSA-774815~Nucleosome assembly | 8 | 3.4 | 0.136 | CENPW, H2BC21, H2AC19, HJURP, KNL1, CENPA, CENPQ, H2AC18 |
| R-HSA-606279~Deposition of new CENPA-containing nucleosomes at the | 8 | 3.4 | 0.136 | CENPW, H2BC21, H2AC19, HJURP, KNL1, CENPA, CENPQ, H2AC18 |
| R-HSA-8854518~AURKA Activation by TPX2 | 8 | 3.4 | 0.136 | PLK4, TPX2, TUBA1A, PLK1, CDK1, NEK2, HMMR, AURKA |
| R-HSA-162658~Golgi Cisternae Pericentriolar Stack Reorganization | 4 | 8.8 | 0.137 | CCNB2, CCNB1, PLK1, CDK1 |
| R-HSA-8953897~Cellular responses to stimuli | 40 | 1.5 | 0.139 | SMARCD3, MT-CO1, EPAS1, CEBPG, H2AC19, SLC7A11, NPAS2, H2AC18, LMNB1, TUBA1B, TUBA1A, TUBB3, H2BC21, RPS6KA2, SESN2, GRB10, E2F2, NLRP3, CHAC1, TBL1X, CDKN2D, ABCC3, UBE2C, HSPA6, ASNS, WIPI1, CCNA2, CCNE2, ID1, MT1F, MT1G, MT-CO2, MT-CO3, ATF5, SQSTM1, HSPA1B, MT1E, ATF4, EZH2, HSPA1A |
| R-HSA-2995410~Nuclear Envelope (NE) Reassembly | 8 | 3.3 | 0.150 | CCNB2, TUBA1B, CCNB1, TUBA1A, TUBB3, CDK1, RCC1, LMNB1 |
| R-HSA-8852276~The role of GTSE1 in G2/M progression after G2 checkp | 8 | 3.2 | 0.170 | CCNB2, TUBA1B, CCNB1, TUBA1A, TUBB3, PLK1, CDK1, GTSE1 |
| R-HSA-73894~DNA Repair | 20 | 1.8 | 0.176 | FANCI, BLM, FEN1, RMI2, PCNA, H2AC19, ISG15, H2AC18, CCNA2, NEIL3, RAD51, EXO1, UBE2T, UVSSA, H2BC21, PCLAF, TCEA1, APBB1, CLSPN, DTL |
| R-HSA-4615885~SUMOylation of DNA replication proteins | 6 | 4.1 | 0.199 | PCNA, INCENP, CDCA8, BIRC5, AURKB, AURKA |
| R-HSA-5633007~Regulation of TP53 Activity | 12 | 2.3 | 0.200 | CCNA2, TPX2, BLM, RMI2, BRPF3, EXO1, SETD9, CDK1, RFFL, AURKB, AURKA, TP73 |
| R-HSA-168256~Immune System | 84 | 1.3 | 0.207 | CD86, DET1, NCKAP1, IFITM1, IL1RN, SERPINA1, ITGAM, FBXO27, CCNF, IFIT5, PIK3CD, IFI35, TREM1, KIF15, LMNB1, OASL, CDC20, HK3, LGALS3, TUBA1B, FCGR3A, TUBA1A, TUBB3, RPS6KA2, FTH1, GRB10, MUC12, CD36, PRKACB, HAVCR2, HERC6, FBXW4, MYLIP, MMP2, KIF23, GAB2, PDCD1LG2, RNASE2, FBXO10, FGR, IRF7, IRF8, BIRC5, TLR8, KIF2C, KIF20A, SQSTM1, IRF9, IFI6, LILRA1, NLRC4, FBXO44, RASGRP3, VTN, RACGAP1, CCL4, S100A12, NLRP3, CD14, KIF3C, PAG1, ATP8B4, SIGLEC15, UBE2C, CCL20, HSPA6, GGH, ISG15, LILRB2, LYZ, BST2, CENPE, KIF18A, FABP5, KIF4A, CDK1, SIGLEC1, HPSE, XAF1, IL7R, ULBP1, HSPA1B, TRIM34, HSPA1A |

| | | | | |
|---|---|---|---|---|
| R-HSA-1280218~Adaptive Immune System | 37 | 1.5 | 0.207 | CD86, DET1, IFITM1, FBXO27, CCNF, PIK3CD, LILRA1, TREM1, FBXO44, KIF15, RASGRP3, CDC20, TUBA1B, FCGR3A, TUBA1A, RACGAP1, TUBB3, KIF3C, CD14, CD36, PRKACB, PAG1, HERC6, FBXW4, MYLIP, UBE2C, KIF23, LILRB2, PDCD1LG2, FBXO10, CENPE, KIF18A, KIF4A, KIF2C, KIF20A, SIGLEC1, ULBP1 |
| R-HSA-68867~Assembly of the pre-replicative complex | 11 | 2.4 | 0.210 | CDT1, ORC1, UBE2C, H2BC21, GMNN, H2AC19, MCM3, MCM4, MCM5, H2AC18, MCM2 |
| R-HSA-9024446~NR1H2 and NR1H3-mediated signaling | 6 | 3.9 | 0.221 | SREBF1, MYLIP, SCD, APOC2, APOC1, TBL1X |
| R-HSA-352230~Amino acid transport across the plasma membrane | 5 | 4.7 | 0.264 | SLC38A1, SLC1A4, SLC7A11, SLC1A5, SLC7A10 |
| R-HSA-2995383~Initiation of Nuclear Envelope (NE) Reformation | 4 | 6.5 | 0.271 | CCNB2, CCNB1, CDK1, LMNB1 |
| R-HSA-176412~Phosphorylation of the APC/C | 4 | 6.5 | 0.271 | CCNB1, UBE2C, PLK1, CDK1 |
| R-HSA-2565942~Regulation of PLK1 Activity at G2/M Transition | 8 | 2.8 | 0.287 | PLK4, CCNB2, CCNB1, TUBA1A, PLK1, CDK1, NEK2, AURKA |
| R-HSA-141430~Inactivation of APC/C via direct inhibition of the APC/C cc | 4 | 6.2 | 0.297 | CDC20, UBE2C, BUB1B, MAD2L1 |
| R-HSA-141405~Inhibition of the proteolytic activity of APC/C required for | 4 | 6.2 | 0.297 | CDC20, UBE2C, BUB1B, MAD2L1 |
| R-HSA-9013026~RHOB GTPase cycle | 7 | 3.1 | 0.297 | NET1, ANLN, DEPDC1B, DIAPH3, RACGAP1, IQGAP3, ARHGEF5 |
| R-HSA-68949~Orc1 removal from chromatin | 7 | 3.0 | 0.312 | CCNA2, CDT1, ORC1, MCM3, MCM4, MCM5, MCM2 |
| R-HSA-162582~Signal Transduction | 101 | 1.2 | 0.314 | NPFFR2, CD86, NCKAP1, ERCC6L, SPARC, H2AC19, BUB1B, H2AC18, CDC20, RGS3, RGS1, TUBB3, RPS6KA2, ADORA3, STMN1, ARHGDIB, MYB, PDK4, GRB10, PDE8A, PRKACB, CMKLR1, MYLIP, KNL1, CDC25C, PYGO1, DEPDC1B, SQSTM1, ROPN1, NSMAF, EPAS1, SH2D2A, CDCA8, ARHGAP18, FPR3, SLC1A5, IQGAP3, PIK3R6, SKA2, RASGRP3, RACGAP1, PAG1, FAM83A, ARHGEF35, PLK1, ZWINT, NDC80, NET1, ANLN, KIF18A, DIAPH3, NOXA1, ID1, CDK1, EZH2, ARHGAP11A, GPSM2, HTR2C, PIK3CD, LMNB1, TUBA1B, TUBA1A, UCN2, H2BC21, TAGAP, NUF2, PDE4B, DLGAP5, TAS2R4, SREBF1, STRADB, MMP2, ARAP2, GAB2, PLCB4, INCENP, BIRC5, KIF2C, ARHGEF5, RFNG, CENPA, AURKB, CCL7, CCL4, SPP1, TBL1X, LAMB3, CCL20, CENPE, FABP5, PRC1, SCD, ADM2, APOC2, APOC1, TACC3, ACKR3, CENPQ, SPC24, SPC25, MAD2L1 |
| R-HSA-5653656~Vesicle-mediated transport | 32 | 1.5 | 0.330 | SERPINA1, HIP1, SPARC, MIA2, KIF15, TUBA1B, ALS2CL, TUBA1A, RACGAP1, MAN2A2, TUBB3, FTH1, KIF3C, CD36, DENND2D, CD163, SGIP1, DENND2B, RAB3IP, SYTL1, KIF23, KIF27, F5, CENPE, MARCO, KIF18A, KIF18B, KIFC1, KIF4A, KIF2C, KIF20A, IL7R |
| R-HSA-2262752~Cellular responses to stress | 37 | 1.4 | 0.330 | SMARCD3, MT-CO1, EPAS1, CEBPG, H2AC19, SLC7A11, NPAS2, H2AC18, LMNB1, TUBA1B, TUBA1A, TUBB3, H2BC21, RPS6KA2, SESN2, GRB10, E2F2, NLRP3, CHAC1, TBL1X, CDKN2D, ABCC3, UBE2C, HSPA6, ASNS, WIPI1, CCNA2, CCNE2, ID1, MT-CO2, MT-CO3, ATF5, SQSTM1, HSPA1B, ATF4, EZH2, HSPA1A |
| R-HSA-179419~APC:Cdc20 mediated degradation of cell cycle proteins pric | 7 | 2.9 | 0.332 | CCNA2, CDC20, UBE2C, CDK1, BUB1B, NEK2, MAD2L1 |
| R-HSA-2299718~Condensation of Prophase Chromosomes | 7 | 2.9 | 0.332 | CCNB1, H2BC21, PLK1, NCAPG2, H2AC19, CDK1, H2AC18 |
| R-HSA-113507~E2F-enabled inhibition of pre-replication complex formatio | 3 | 10.3 | 0.342 | CCNB1, ORC1, CDK1 |
| R-HSA-9013106~RHOC GTPase cycle | 7 | 2.9 | 0.342 | ANLN, DEPDC1B, DIAPH3, RACGAP1, ARHGAP18, IQGAP3, ARHGEF5 |
| R-HSA-9013423~RAC3 GTPase cycle | 8 | 2.6 | 0.342 | NCKAP1, DEPDC1B, DIAPH3, RACGAP1, NOXA1, ARHGDIB, ARAP2, SLC1A5 |
| R-HSA-9833482~PKR-mediated signaling | 7 | 2.9 | 0.356 | TUBA1B, TUBA1A, TUBB3, CDK1, ISG15, HSPA1B, HSPA1A |
| R-HSA-174048~APC/C:Cdc20 mediated degradation of Cyclin B | 4 | 5.4 | 0.378 | CDC20, CCNB1, UBE2C, CDK1 |

| | | | | |
|---|---|---|---|---|
| R-HSA-9709603~Impaired BRCA2 binding to PALB2 | 4 | 5.1 | 0.418 | BLM, RMI2, RAD51, EXO1 |
| R-HSA-9701193~Defective homologous recombination repair (HRR) due tc | 4 | 4.9 | 0.440 | BLM, RMI2, RAD51, EXO1 |
| R-HSA-9704331~Defective HDR through Homologous Recombination Rep | 4 | 4.9 | 0.440 | BLM, RMI2, RAD51, EXO1 |
| R-HSA-9704646~Defective HDR through Homologous Recombination Rep | 4 | 4.9 | 0.440 | BLM, RMI2, RAD51, EXO1 |
| R-HSA-9701192~Defective homologous recombination repair (HRR) due tc | 4 | 4.9 | 0.440 | BLM, RMI2, RAD51, EXO1 |
| R-HSA-2173782~Binding and Uptake of Ligands by Scavenger Receptors | 5 | 3.7 | 0.440 | MARCO, CD163, SPARC, FTH1, CD36 |
| R-HSA-2426168~Activation of gene expression by SREBF (SREBP) | 5 | 3.7 | 0.440 | SREBF1, SMARCD3, SCD, ELOVL6, TBL1X |
| R-HSA-5661231~Metallothioneins bind metals | 3 | 8.4 | 0.448 | MT1F, MT1G, MT1E |
| R-HSA-1280215~Cytokine Signaling in Immune system | 35 | 1.4 | 0.483 | CD86, IFITM1, IL1RN, ITGAM, IFIT5, IFI6, PIK3CD, IFI35, LMNB1, OASL, TUBA1B, TUBA1A, TUBB3, RPS6KA2, CCL4, GRB10, S100A12, CD36, HAVCR2, CCL20, MMP2, ISG15, GAB2, BST2, IRF7, CDK1, IRF8, BIRC5, XAF1, IL7R, SQSTM1, IRF9, HSPA1B, TRIM34, HSPA1A |
| R-HSA-2559583~Cellular Senescence | 12 | 1.9 | 0.487 | CDKN2D, CCNA2, CCNE2, UBE2C, RPS6KA2, H2BC21, ID1, H2AC19, E2F2, H2AC18, LMNB1, EZH2 |
| R-HSA-1368108~BMAL1:CLOCK,NPAS2 activates circadian gene expres | 4 | 4.6 | 0.503 | SMARCD3, NAMPT, TBL1X, NPAS2 |
| R-HSA-5693554~Resolution of D-loop Structures through Synthesis-Deper | 4 | 4.6 | 0.503 | BLM, RMI2, RAD51, EXO1 |
| R-HSA-9707564~Cytoprotection by HMOX1 | 6 | 2.9 | 0.503 | SMARCD3, MT-CO1, NLRP3, MT-CO2, MT-CO3, TBL1X |
| R-HSA-168330~Viral RNP Complexes in the Host Cell Nucleus | 2 | 30.8 | 0.558 | HSPA1B, HSPA1A |
| R-HSA-68881~Mitotic Metaphase/Anaphase Transition | 2 | 30.8 | 0.558 | PLK1, FBXO5 |
| R-HSA-75035~Chk1/Chk2(Cds1) mediated inactivation of Cyclin B:Cdk1 c | 3 | 7.1 | 0.558 | CCNB1, CDK1, CDC25C |
| R-HSA-68884~Mitotic Telophase/Cytokinesis | 3 | 7.1 | 0.558 | PLK1, KIF23, KIF20A |
| R-HSA-437239~Recycling pathway of L1 | 5 | 3.3 | 0.558 | TUBA1B, TUBA1A, TUBB3, RPS6KA2, KIF4A |
| R-HSA-157579~Telomere Maintenance | 8 | 2.2 | 0.608 | CCNA2, BLM, FEN1, PCNA, DSCC1, H2BC21, H2AC19, H2AC18 |
| R-HSA-5660526~Response to metal ions | 3 | 6.6 | 0.622 | MT1F, MT1G, MT1E |
| R-HSA-196807~Nicotinate metabolism | 4 | 4.0 | 0.645 | PARP6, NMRK1, NAMPT, PARP9 |
| R-HSA-6804756~Regulation of TP53 Activity through Phosphorylation | 7 | 2.3 | 0.645 | CCNA2, TPX2, BLM, RMI2, EXO1, AURKB, AURKA |
| R-HSA-9013149~RAC1 GTPase cycle | 11 | 1.8 | 0.645 | NCKAP1, DEPDC1B, DIAPH3, RACGAP1, TAGAP, NOXA1, ARHGDIB, ARAP2, SLC1A5, IQGAP3, ARHGEF5 |
| R-HSA-5693538~Homology Directed Repair | 9 | 2.0 | 0.648 | CCNA2, BLM, FEN1, RMI2, RAD51, PCNA, EXO1, H2BC21, CLSPN |
| R-HSA-9675135~Diseases of DNA repair | 5 | 3.0 | 0.648 | BLM, RMI2, NEIL3, RAD51, EXO1 |
| R-HSA-180786~Extension of Telomeres | 5 | 3.0 | 0.648 | CCNA2, BLM, FEN1, PCNA, DSCC1 |
| R-HSA-4419969~Depolymerization of the Nuclear Lamina | 3 | 6.2 | 0.648 | CCNB1, CDK1, LMNB1 |
| R-HSA-174184~Cdc20:Phospho-APC/C mediated degradation of Cyclin A | 6 | 2.6 | 0.648 | CCNA2, CDC20, UBE2C, CDK1, BUB1B, MAD2L1 |
| R-HSA-5626467~RHO GTPases activate IQGAPs | 4 | 3.8 | 0.648 | TUBA1B, TUBA1A, TUBB3, IQGAP3 |
| R-HSA-69473~G2/M DNA damage checkpoint | 7 | 2.3 | 0.648 | BLM, CCNB1, RMI2, EXO1, H2BC21, CDK1, CDC25C |
| R-HSA-380320~Recruitment of NuMA to mitotic centrosomes | 7 | 2.3 | 0.648 | PLK4, TUBA1B, TUBA1A, TUBB3, PLK1, CDK1, NEK2 |
| R-HSA-1989781~PPARA activates gene expression | 8 | 2.1 | 0.648 | SREBF1, SMARCD3, ACSL1, G0S2, PLIN2, CD36, TBL1X, NPAS2 |
| R-HSA-1500620~Meiosis | 8 | 2.1 | 0.648 | BLM, RAD51, H2BC21, H2AC19, MND1, H2AC18, SYNE1, LMNB1 |
| R-HSA-2980766~Nuclear Envelope Breakdown | 5 | 3.0 | 0.648 | CCNB2, CCNB1, PLK1, CDK1, LMNB1 |

| | | | | |
|---|---|---|---|---|
| R-HSA-174178~APC/C:Cdh1 mediated degradation of Cdc20 and other AP | 6 | 2.5 | 0.648 | CDC20, PTTG1, UBE2C, PLK1, AURKB, AURKA |
| R-HSA-68875~Mitotic Prophase | 9 | 2.0 | 0.648 | CCNB2, CCNB1, H2BC21, PLK1, NCAPG2, H2AC19, CDK1, H2AC18, LMNB1 |
| R-HSA-199991~Membrane Trafficking | 28 | 1.4 | 0.648 | SERPINA1, HIP1, MIA2, KIF15, TUBA1B, ALS2CL, TUBA1A, RACGAP1, MAN2A2, TUBB3, FTH1, KIF3C, DENND2D, SGIP1, DENND2B, RAB3IP, SYTL1, KIF23, KIF27, F5, CENPE, KIF18A, KIF18B, KIFC1, KIF4A, KIF2C, KIF20A, IL7R |
| R-HSA-400206~Regulation of lipid metabolism by PPARalpha | 8 | 2.1 | 0.668 | SREBF1, SMARCD3, ACSL1, G0S2, PLIN2, CD36, TBL1X, NPAS2 |
| R-HSA-1362300~Transcription of E2F targets under negative control by p1( | 3 | 5.8 | 0.674 | CCNA2, CDK1, MYBL2 |
| R-HSA-170145~Phosphorylation of proteins involved in the G2/M transitio | 2 | 20.5 | 0.675 | CCNA2, CDK1 |
| R-HSA-5693532~DNA Double-Strand Break Repair | 10 | 1.8 | 0.678 | CCNA2, BLM, FEN1, RMI2, RAD51, PCNA, EXO1, H2BC21, APBB1, CLSPN |
| R-HSA-174417~Telomere C-strand (Lagging Strand) Synthesis | 4 | 3.6 | 0.678 | BLM, FEN1, PCNA, DSCC1 |
| R-HSA-5693568~Resolution of D-loop Structures through Holliday Junctio | 4 | 3.6 | 0.678 | BLM, RMI2, RAD51, EXO1 |

Red indicates FDR < 0.05.

**Supplementary Table 7. Transcription factors predicted by iRegulon**

**Transcription factors predicted by DEGs upregulated in HLF cells**

| Rank | Track id | AUC | NES | ClusterCode | Transcription factor | Target genes |
|---|---|---|---|---|---|---|
| 1 | GSM1208780_batch2_chrom1_LoVo_NKX2.2_PassedQC_peaks_hg19 | 0.161 | 5.054 | T1 | NKX2.2 | PSAT1,ASNS,FKBP14 |
| 2 | wgEncodeHaibTfbsA549MaxV0422111PkRep2.broadPeak.gz | 0.150 | 4.651 | T2 | MAX | PSAT1,SYCE2 |
| 3 | wgEncodeSydhTfbsHepg2CebpbForsklnStdPk.narrowPeak.gz | 0.146 | 4.532 | T3 | CEBPB | PSAT1,SLC7A2 |
| 4 | wgEncodeHaibTfbsK562Stat5asc74442V0422111PkRep1.broadPeak.gz | 0.138 | 4.249 | T4 | STAT5A | RELN,SLC7A2 |
| 5 | wgEncodeHaibTfbsK562Trim28sc81411V0422111PkRep2.broadPeak.gz | 0.133 | 4.074 | T5 | TRIM28 | RELN,SLC7A2 |
| 6 | wgEncodeSydhTfbsK562Gata1bIggmusPk.narrowPeak.gz | 0.132 | 4.050 | T6 | GATA1 | SLC7A2,ABLIM3,RELN,PAPPA2,GABRG1,ABCA10,SYCE2 |
| 7 | wgEncodeSydhTfbsH1hescUsf2IggrabPk.narrowPeak.gz | 0.132 | 4.038 | T7 | USF2 | PAPPA2,RELN |
| 8 | wgEncodeSydhTfbsHelas3Prdm19115IggrabPk.narrowPeak.gz | 0.132 | 4.034 | T8 | PRDM1 | ABLIM3,SLC7A2,SLIT2,ABCA10,PSAT1 |
| 9 | GSM1208731_batch2_chrom1_LoVo_E2F8_PassedQC_peaks_hg19 | 0.129 | 3.926 | T9 | E2F8 | PSAT1,ASNS |
| 10 | wgEncodeHaibTfbsK562Stat5asc74442V0422111PkRep2.broadPeak.gz | 0.129 | 3.914 | T4 | STAT5A | RELN,SLC7A2 |
| 11 | wgEncodeHaibTfbsK562Gata2sc267Pcr1xPkRep2.broadPeak.gz | 0.128 | 3.902 | T10 | GATA2 | RELN,SLC7A2 |
| 12 | wgEncodeSydhTfbsPbdefetalGata1UcdPk.narrowPeak.gz | 0.128 | 3.898 | T6 | GATA1 | GPR156,SLC7A2,RELN,ASNS,SLIT2,ABLIM3,GABRG1 |
| 13 | wgEncodeUchicagoTfbsK562Egata2ControlPk.narrowPeak.gz | 0.121 | 3.639 | T10 | GATA2 | RELN,SLC7A2,SYCE2 |
| 14 | wgEncodeHaibTfbsK562Tead4sc101184V0422111PkRep2.broadPeak.gz | 0.117 | 3.528 | T11 | TEAD4 | SLC7A2,RELN |
| 15 | wgEncodeHaibTfbsHct116Rad21V0422111PkRep1.broadPeak.gz | 0.113 | 3.392 | T12 | RAD21 | ABCA10,GPR156 |
| 16 | wgEncodeSydhTfbsHelas3CfosStdPk.narrowPeak.gz | 0.113 | 3.376 | T13 | FOS | SERPINB2,ABLIM3,FKBP14,PSAT1 |
| 17 | wgEncodeSydhTfbsK562Irf1Ifna6hStdPk.narrowPeak.gz | 0.111 | 3.317 | T14 | IRF1 | RELN,ASNS |
| 18 | wgEncodeHaibTfbsGm12878Rad21V0416101PkRep1.broadPeak.gz | 0.106 | 3.125 | T12 | RAD21 | ABCA10,GPR156 |
| 19 | wgEncodeSydhTfbsHuvecCfosUcdPk.narrowPeak.gz | 0.105 | 3.106 | T13 | FOS | ABLIM3,SERPINB2,SLC7A2 |
| 20 | wgEncodeSydhTfbsHelas3CmycStdPk.narrowPeak.gz | 0.105 | 3.086 | T15 | MYC | SERPINB2,ABLIM3 |

**Transcription factors predicted by DEGs downregulated in HLF cells**

| Rank | Track id | AUC | NES | ClusterCode | Transcription factor | Target genes |
|---|---|---|---|---|---|---|
| 1 | wgEncodeSydhTfbsK562Stat2Ifna6hStdPk.narrowPeak.gz | 0.464 | 14.397 | T1 | STAT2 | PARP9,IFIT5,MX1,OAS2,IFITM3,EPSTI1,PLSCR1,HERC5,XAF1,OAS1,IFI44L,ISG15,IFIH1,HSH2D,STAT1,CMPK2,SP110,IFI35,TMEM140,LAP3,SAMHD1,ISG20,TRIM14 |
| 2 | wgEncodeSydhTfbsK562Stat1Ifna6hStdPk.narrowPeak.gz | 0.448 | 13.877 | T2 | STAT1 | PARP9,OAS2,MX1,IFIT5,EPSTI1,ISG15,PLSCR1,OAS1,LAP3,TRIM14,IFITM3,STAT1,XAF1,IFI44L,IFIH1,TMEM140,SP110,HSH2D,CMPK2,IFI35,SAMHD1,ISG20 |
| 3 | wgEncodeSydhTfbsK562Stat2Ifna30StdPk.narrowPeak.gz | 0.391 | 12.013 | T1 | STAT2 | MX1,PARP9,EPSTI1,SP110,STAT1,IFIT5,IFIH1,PLSCR1,LAP3,ISG15,CMPK2,OAS1,IFITM3,TMEM140,XAF1,IFI35,OAS2,TRIM14,SAMHD1 |
| 4 | wgEncodeSydhTfbsK562Stat1Ifna30StdPk.narrowPeak.gz | 0.350 | 10.697 | T2 | STAT1 | PARP9,MX1,STAT1,PLSCR1,CMPK2,ISG15,SP110,LAP3,IFIT5,EPSTI1,IFIH1,TMEM140,OAS1,IFITM3,IFI35,TRIM14,SAMHD1 |
| 5 | wgEncodeSydhTfbsK562Irf1Ifna30StdPk.narrowPeak.gz | 0.198 | 5.755 | T3 | IRF1 | MX1,PARP9,IFIH1,IFI35,TMEM140,STAT1,ISG15,SP110,SAMHD1,IFIT5,XAF1,IFITM3,PLSCR1,LPCAT3,CMPK2,ANKRD37 |
| 6 | wgEncodeSydhTfbsK562Stat1Ifng6hStdPk.narrowPeak.gz | 0.144 | 4.011 | T2 | STAT1 | PARP9,STAT1,LAP3,EPSTI1,CXCL10,TMEM140,PLSCR1,E2F1,IFI44L,IFI35 |

**Transcription factors predicted by DEGs upregulated in macrophages**

| Rank | Track id | AUC | NES | ClusterCode | Transcription factor | Target genes |
|---|---|---|---|---|---|---|

| | | | | | | |
|---|---|---|---|---|---|---|
| 1 | GSM1208731_batch2_chrom1_LoVo_E2F8_PassedQC_peaks_hg19 | 0.064 | 5.122 | T1 | E2F8 | NFE2L1,RNF187,PSAT1,ASNS,SESN2,SLC7A11,CCPG1,TCEA1,SLC1A5,BEST1,CHAC1,GPT2,ULBP1,ZNF181,IFRD1,NCKAP1,NUPR1,PCK2,GTPBP2,SLC38A1,MTHFD2,WIPI1,XPOT,MKNK2,PHGDH,TSEN15,GRB10,INHBE |
| 2 | wgEncodeHaibTfbsK562Cebpbsc150V0422111PkRep2.broadPeak.gz | 0.058 | 4.519 | T2 | CEBPB | CLGN,RFFL,SLC48A1,GRB10,SMARCD3,BEST1,FAM83A,EPAS1,TBL1X,ULBP1,INHBE,PYCR1,TNNC1,ATF5,NCKAP1,SLC38A1,MKNK2,XPOT,MTHFD2,PHGDH,RNF187,SLC7A11,NFE2L1,DHRS11,GTPBP2,RAB3IP,SESN2,CEBPG,ASNS,FTH1,CHAC1,HIP1,ZFAND3,TSEN15,GTF2IRD2B,GAB2,CCPG1,SLC1A4,SLC22A15,CBS,SQSTM1,ZNF181,SLC1A5,RPS6KA2,PNPLA1,MAP1B,TCEA1,NPFFR2,RUFY3,P2RX4,PAG1 |
| 3 | wgEncodeSydhTfbsH1hescCebpbIggrabPk.narrowPeak.gz | 0.054 | 4.080 | T2 | CEBPB | TMEM182,TNIP3,TBX15,SYBU,CBS,CHAC1,ULBP1,TNNC1,PYCR1,SESN2,MKNK2,TESK1,GPT2,FAM83A,PCDH1,ASNS,CACNA1F,PLCB4,MAP1B,SLC38A1,SLC1A5,INHBE,CLGN,ABCC3,NFE2L1,ZBED3,SQSTM1,MTHFD2,EPAS1,ELOVL6,TTLL4,ZFAND3,SLC7A11,PHGDH,GRB10,SNN,CYP4V2,PCDHAC2,BEST1,SLC48A1,PSAT1,GAB2,UNC13A,PYGO1,NUPR1,MAP7,XPOT,RNF187,ZFP41,NAMPT,MT1F,PCK2,ATF5,TBL1X,PPIL6,ADM2,SPIRE1,SMARCD3,ZDHHC1,GPCPD1,SNHG7,HIP1,ZNF853,GDF1,TSEN15,SLC1A4,P2RX4,SPATA18,UNC5B,NCKAP1,STC2,PRSS27,NPAS2,SLC7A10,SEL1L3,SLC2A10,RPS6KA2,MCOLN3,DET1,CEBPG,TMEM150A,DCAF10,RUFY3,SERPINB7,ALDH1L2,ERMAP,SPEG |
| 4 | wgEncodeHaibTfbsEcc1Cebpbsc150V0422111PkRep1.broadPeak.gz | 0.054 | 4.007 | T2 | CEBPB | CHAC1,SLC48A1,CLGN,TNNC1,FAM83A,SLC38A1,EPAS1,MKNK2,TBL1X,PYCR1,RFFL,SLC7A11,CEBPG,RAB3IP,ALDH1L2,ZBED3,JDP2,SQSTM1,ABCC3,ZFAND3,PLCB4,PSAT1,ASNS,GRB10,SESN2,SPATA18,XPOT,INHBE,ULBP1,RPS6KA2,STRBP,CLIP4,BEST1,SLC1A4,FTH1,SLC1A5,NR1D2,MTHFD2,CBS,TSEN15,NCKAP1,SYBU,CYP4V2,DHRS12,SEL1L3,PHGDH,GPT2,IFRD1,PER3,MT1F,PNPLA1,RNF187,STC2,SMARCD3,AKAP5,DCAF10,FBXO27,P2RX4,NPL,UNC5B,ADM2 |
| 5 | GSM1208598_batch1_chrom1_LoVo_CEBPB_PassedQC_peaks_hg19 | 0.053 | 3.977 | T2 | CEBPB | SLC38A1,MKNK2,XPOT,ZBED3,TNNC1,LGALS3,RNF187,ASNS,SESN2,CCPG1,EPAS1,CHAC1,RAB3IP,GRB10,SLC7A11,PSAT1,RFFL,FAM83A,ABCC3,IFRD1,SQSTM1,BEST1,GPT2,GTPBP2 |
| 6 | wgEncodeHaibTfbsK562Cebpbsc150V0422111PkRep1.broadPeak.gz | 0.053 | 3.964 | T2 | CEBPB | INHBE,CLGN,FAM83A,SLC7A11,BEST1,RFFL,TBL1X,GRB10,SLC38A1,TNNC1,SLC48A1,MKNK2,ULBP1,PYCR1,XPOT,MAN2A2,SESN2,NCKAP1,ASNS,CHAC1,DHRS11,EPAS1,SMARCD3,FTH1,GTPBP2,RNF187,NFE2L1,SEL1L3,ZNF181,MTHFD2,RAB3IP,SQSTM1,SLC1A5,CCPG1,TSEN15,ZFAND3,ATF5,PHGDH,CEBPG,ERMAP,NR1D2,SLC38A6,TLX2,CBS,TCEA1,P2RX4,HIP1,PCK2,PNPLA1,ELOVL6,KCNMB3,SLC1A4,FECH,TESK1,NR4A3 |

| | | | | | | |
|---|---|---|---|---|---|---|
| 7 | wgEncodeHaibTfbsHct116Cebpbsc150V0422111PkRep1.broadPeak.gz | 0.050982 | 3.71627 | T2 | CEBPB | RFFL,FAM83A,CLGN,CLIP4,SLC7A11,TNNC1,CHAC1,RNF187,TBL1X,FLYWCH1,SLC22A15,NCKAP1,BEST1,SESN2,SLFN5,GTPBP2,MKNK2,XPOT,SLC38A1,NFE2L1,SLC48A1,TCEA1,MT1F,PYCR1,ULBP1,RAB3IP,SIPA1L2,CCPG1,EPAS1,INHBE,RPS6KA2,ZFAND3,ABCC3,FTH1 |
| 8 | wgEncodeSydhTfbsH1hescBach1sc14700IggrabPk.narrowPeak.gz | 0.0503215 | 3.64311 | T3 | BACH1 | EPAS1,SLC30A1,GRB10,TESK1,CACNA1F,SQSTM1,SPIRE1,ULBP1,D2HGDH,FECH,SLC48A1,GTF2IRD2B,SNHG7,NAMPT,MKNK2,LRRC24,SLC7A11,FTH1,NPAS2,IFRD1,TMEM150A,PHF1,ERMAP,RNF187,SEL1L3,ACSL1,RFFL |
| 9 | wgEncodeSydhTfbsImr90CebpbIggrabPk.narrowPeak.gz | 0.0497167 | 3.57612 | T2 | CEBPB | GPT2,SLC1A5,PRSS27,NUPR1,PNPLA1,BEST1,SLC7A10,CLGN,TNNC1,MTHFD2,CHAC1,PYCR1,SLC1A4,ELOVL6,PHGDH,TBL1X,SESN2,MKNK2,GAB2,NCKAP1,DENND2D,INHBE,ABCC3,ASNS,NFE2L1,PSAT1,EPAS1,PLCB4,RFFL,SLC38A1,CLIP4,GRB10,TNIP3,TTLL4,SLC48A1,CEBPG,SLC7A11,FLYWCH1,RNF187,PCDHAC2 |
| 10 | GSM1208709_batch2_chrom1_LoVo_AEBP2_PassedQC_peaks_hg19 | 0.0493274 | 3.533 | T4 | AEBP2 | MKNK2,PSAT1,FTH1,PCK2,CHAC1,ZFAND3,CCPG1,ARVCF,NUPR1,XPOT,GPT2,TCEA1,SESN2,BEST1,NFE2L1,ADM2,CEBPG,PHGDH,RNF187,C1QTNF6,MT1E,GTPBP2,MUC12,UNC5B,SMARCD3,SEL1L3,DHRS12,FBXW4 |
| 11 | wgEncodeHaibTfbsA549Cebpbsc150V0422111PkRep1.broadPeak.gz | 0.0492787 | 3.52761 | T2 | CEBPB | RFFL,ABCC3,EPAS1,TBL1X,CLGN,SLC7A11,PYCR1,CHAC1,SLC48A1,CLIP4,SLC38A1,ZBED3,TNNC1,FTH1,SLC1A5,GRB10,NCKAP1,SQSTM1,MKNK2,FAM83A,PLCB4,CEBPG,CAPN5,PSAT1,NUPR1,XPOT,ULBP1,BEST1,MTHFD2,RAB3IP,SDSL,FBXO27,DHRS12,NR1D2,ASNS,INHBE,ZFAND3,CBS,MUC12,ZNF181,TCEA1,GAB2,C1QTNF6,TESK1,IFRD1,KCTD15,AKAP5,KIF27,PCDHAC2,JDP2,TSEN15,TNIP3,SLC1A4,SMARCD3,ELOVL6,PYGO1,SESN2,NFE2L1,HIP1,PNPLA1,PCK2,PDE8A,BIRC7,NR4A3,MAP7,SIPA1L2,MT1F,STC2,ERMAP,TUBE1,TLX2,PDK4,CTNNA3,MAP1B,P2RX4,FLYWCH1,ZNF425,HEMK1,PER3,CCPG1,SLFN5,DTNB,BEND6,ALDH1L2 |
| 12 | wgEncodeHaibTfbsA549Cebpbsc150V0422111PkRep2.broadPeak.gz | 0.0484374 | 3.43443 | T2 | CEBPB | RFFL,SLC7A11,CLGN,CHAC1,SLC48A1,EPAS1,CLIP4,FAM83A,SLC38A1,GRB10,TBL1X,ABCC3,SQSTM1,FTH1,TNNC1,SLC1A5,INHBE,ZBED3,PLCB4,NCKAP1,MKNK2,PYCR1,NUPR1,RAB3IP,XPOT,TSEN15,KIF27,CBS,PNPLA1,MT1F,BEST1,TNIP3,PSAT1,ZFAND3,CEBPG,DHRS12,ULBP1,ELOVL6,SESN2,TESK1,AKAP5,MTHFD2,PDE8A,SMARCD3,PYGO1,SDSL,MUC12,SLC1A4,PCK2,BIRC7,HIP1,FBXO27,NR4A3,ASNS,PCDHAC2,SLC2A10,ZNF181 |

| | | | | | | |
|---|---|---|---|---|---|---|
| 13 | wgEncodeSydhTfbsHelas3CebpbIggrabPk.narrowPeak.gz | 0.0476379 | 3.34588 | T2 | CEBPB | STRC,TRIM16L,TNNC1,SMARCD3,SQSTM1,RILPL1,ADM2,PER3,PYCR1,CLGN,ASNS,ZNF425,SLC38A1,SLC1A4,HIP1,MKNK2,NFE2L1,BEST1,DENND2D,ULBP1,INHBE,MT1F,JDP2,CEBPG,EPAS1,GAB2,P2RX4,PCDHAC2,GRB10,CHAC1,FLYWCH1,KCTD15,SLC7A10,CCDC112,SIPA1L2,PCK2,ELOVL6,SLC1A5,ZDHHC1,PSAT1,NUPR1,RNF187,GPT2,UNC13A,TBL1X,SLC22A15,MTHFD2,RAB3IP,NPAS2,PNPLA1,SEL1L3,SLC48A1,LGALS3,ZBED3,FGR,STC2,FTH1,NBPF1,FAM83A,RFFL,ABCC3,XPOT,DCAF10,MYO15B,NR1D2,GTPBP2 |
| 14 | wgEncodeSydhTfbsK562Nfe2StdPk.narrowPeak.gz | 0.0469844 | 3.27349 | T5 | NFE2 | ERMAP,IFRD1,SLC48A1,FTH1,DET1,BEND6,CCPG1,SEL1L3,NR1D2,GAB2,SLC1A5,SMARCD3,SQSTM1,RFFL,SLC7A11 |
| 15 | wgEncodeHaibTfbsEcc1Cebpbsc150V0422111PkRep2.broadPeak.gz | 0.0447527 | 3.02631 | T2 | CEBPB | SLC48A1,CLGN,SLC7A11,FAM83A,RFFL,CHAC1,SLC38A1,SESN2,TNNC1,GRB10,RAB3IP,EPAS1,ALDH1L2,MKNK2,TBL1X,INHBE,ZBED3,NCKAP1,BEST1,SMARCD3,PLCB4,ZFAND3,FBXO27,CBS,ABCC3,JDP2,FTH1,SEL1L3,MT1F,ULBP1,XPOT,PNPLA1,PYCR1,SQSTM1,PSAT1,AKAP5,RPS6KA2,SPATA18,PHGDH,NUPR1,DHRS12,CLIP4,STRBP,NFE2L1,GPT2,MTHFD2,KIF27,SLC1A5,ASNS,ERMAP,RNF187,TSEN15,NPL,RUFY3,STC2,TNIP3,NLRC4 |

**Transcription factors predicted by DEGs downregulated in macrophages**

| Rank | Track id | AUC | NES | ClusterCode | Transcription factor | Target genes |
|---|---|---|---|---|---|---|
| 1 | wgEncodeSydhTfbsHelas3E2f4StdPk.narrowPeak.gz | 0.244247 | 11.5994 | T1 | E2F4 | SPC25,DIAPH3,KIF20A,ESCO2,CENPW,KIF2C,NUF2,CDC25C,AURKB,ZWINT,MAD2L1,MCM3,PBK,CDC7,NDC80,UBE2T,ARHGAP11A,NEIL3,KIF18B,CDCA5,NCAPH,ATAD2,KIF15,PLK1,CDKN2D,TK1,CDC45,NEK2,SPC24,FOXM1,CDK1,KIF4A,PCNA,RAD54B,MELK,GTSE1,IQGAP3,SHCBP1,RAD54L,BUB1B,DTL,ANLN,MYBL2,BLM,SPAG5,CDCA3,CDT1,KIF18A,DLGAP5,HJURP,DEPDC1B,BIRC5,NCAPG2,TCF19,PRC1,HELLS,EXO1,TTK,NCAPG,INCENP,CDCA8,ESPL1,TPX2,ASF1B,TROAP,GGH,UBE2C,FEN1,FANCI,MCM4,WDR76,UHRF1,SKA2,CENPQ,GINS1,NET1,CENPA,KIF23,MND1,CLSPN,GINS2,NUSAP1,HMGB3,DSCC1,EZH2,ORC1,RACGAP1,TUBA1B,FBXO5,CCNA2,RCC1,TRIP13,RAD51,GMNN,CDC25A,KIFC1,TYMS,PLK4,LMNB1,ZNF367,MCM2,RGS3,AURKA,CDC20,CCNE2,MXD3,PKMYT1,E2F2,CORO7,ERCC6L,CDKN3 |

| | | | | | | |
|---|---|---|---|---|---|---|
| 2 | GSM1208691_batch1_chrom1_LoVo_TFDP1_PassedQC_peaks_hg19 | 0.202441 | 9.43286 | T2 | TFDP1 | SKA2,CDK1,MCM5,NDC80,KIF23,PRC1,EXO1,NEK2,CDC25A,KIF18B,CDKN2D,FBXO5,PLK4,CDC7,FEN1,FANCI,INCENP,CDCA8,AURKB,ZWINT,NUF2,CDCA5,IQGAP3,DIAPH3,ESCO2,CLSPN,SPC24,ASF1B,CDC25C,TUBA1B,MCM3,BLM,PCNA,KIF2C,PBK,TK1,CDCA3,ATAD2,SPC25,BUB1B,HJURP,DTL,RAD51,GMNN,NCAPG,NUSAP1,KIF18A,SPAG5,TROAP,DLGAP5,UBE2C,UBE2T,CCNA2,NEIL3,CENPW,PLK1,MAD2L1,RAD54B,RACGAP1,CCNB1,CENPA,CDKN3,RAD54L,MXD3,E2F2,TYMS,KIF15,WDR76,ESPL1,SNRNP25,CCNB2,GINS1,CENPQ,BIRC5,GTSE1,KIF20A,CCNE2,TTK,SLC25A19,NET1,CDC20,ORC1,MELK,LMNB1,NCAPG2,KIF4A,SHCBP1,TPX2,ARHGAP11A,ZNF367,HMGB3,ERCC6L |
| 3 | wgEncodeSydhTfbsGm12878E2f4IggmusPk.narrowPeak.gz | 0.191442 | 8.86286 | T1 | E2F4 | ARHGAP11A,NCAPG2,IQGAP3,BUB1B,CDC20,ANLN,MYBL2,SPC24,TRIP13,DIAPH3,DEPDC1B,CDC25C,TROAP,RAD54L,AURKB,TACC3,NEK2,GGH,CDCA8,KIF4A,SPAG5,KIF20A,PRC1,HMGB3,FOXM1,KIF18B,GINS2,DLGAP5,BIRC5,ASF1B,KIF23,ESCO2,DSCC1,KIF2C,TTK,INCENP,CDCA5,PLK1,TK1,KIF15,HERC6,CDKN3,NCAPG,UBE2C,GTSE1,SPC25,CCNB2,CDKN2D,UHRF1,UBE2T,MND1,BLM,CDK1,CDC25A,CD86,PLK4,FEN1,ATAD2,NEIL3,FBXO5,CLSPN,NCAPH,SHCBP1,EXO1,PBK,ZWINT,MAD2L1,NDC80,OASL,ERCC6L,ZNF367,RAD51,CCNB1,MCM5,CENPW,CENPA,PKMYT1,CDCA3,CCNA2,MXD3,KIF18A,NUF2,GPSM2,TCF19,MELK,CKS2,SNRNP25,HJURP,TYMS,ESPL1,NUSAP1,RACGAP1,NET1,RCC1,MCM3,PCNA,MCM4,CDC7,FANCI,WDR76,CCNE2,E2F2,KIFC1,RAD54B,CENPQ,ID1,TPX2,HMMR,CDT1,TAL1,MCM2,SKA2,DTL,LMNB1,CCNF,RGS3,EZH2,TUBA1A,PTTG1 |
| 4 | wgEncodeSydhTfbsK562E2f4UcdPk.narrowPeak.gz | 0.190607 | 8.81955 | T1 | E2F4 | DIAPH3,FOXM1,NCAPG2,BIRC5,TK1,NEIL3,NCAPG,IQGAP3,TRIP13,GINS2,TACC3,EZH2,CDT1,MND1,KIF15,MCM2,AURKB,BUB1B,ANLN,SHCBP1,WDR76,ZWINT,CDCA5,UBE2T,CDK1,NUF2,ESCO2,ATAD2,ZNF367,MCM4,CDC25A,HMGB3,GTSE1,PBK,MYBL2,CENPA,UHRF1,INCENP,MID1IP1,ASF1B,MELK,CDC25C,HJURP,SPC25,TCF19,CENPW,KIF20A,NEK2,PKMYT1,ARHGAP11A,PRC1,CDC7,NCAPH,DEPDC1B,RAD54L,CLSPN,KIF4A,CCNA2,KIF23,EXO1,PLK4,CORO7,MAD2L1,DSCC1,NUSAP1,MCM5,CDKN2D,SNRNP25,BLM,PCNA,KIF2C,KIF18B,KIFC1,FBXO5,MXD3,KIF18A,SCD,CDKN3,NET1,DTL,CDCA3,RAD51,TYMS,STMN1,LMNB1,GINS1,GGH,PLK1,MCM3,AQP1,RACGAP1,RCC1,CCNF,FANCI,DLGAP5,TUBA1B,GMNN,ESPL1,ERCC6L |

| | | | | | | |
|---|---|---|---|---|---|---|
| 5 | wgEncodeHaibTfbsEcc1Foxm1sc502V0422111PkRep2.broadPeak.gz | 0.18165 | 8.35538 | T3 | FOXM1 | CCNB1,CDK1,GPSM2,CDKN3,UBE2C,CENPA,TROAP,GTSE1,CCNA2,TPX2,CCNF,LMNB1,MXD3,CCNB2,PRC1,PLK1,SKA2,KIF2C,NCAPH,RACGAP1,NUF2,KIF18B,PTTG1,KIF23,KIF20A,HJURP,KIF18A,HMMR,INCENP,RGS3,NUSAP1,CD36,UBE2T,CDKN2D,ESPL1,KIF4A,AURKA,UHRF1,NEK2,KIFC1,TTK,CDC20,AURKB,NEIL3,ANLN,FBXO5,TK1,MAD2L1,S100A10,ATAD2,DLGAP5,BIRC5,CDCA8,E2F2,ASF1B,NDC80,KIF15,EZH2,ID1,NCAPG,PBK,DIAPH3,CDC25C,LAMB3,TACC3,CENPE,G0S2,SPAG5,ESCO2,PCNA,HMGB3,TYMS |
| 6 | wgEncodeHaibTfbsEcc1Foxm1sc502V0422111PkRep1.broadPeak.gz | 0.16689 | 7.5905 | T3 | FOXM1 | GPSM2,CDK1,CCNB1,CDKN3,CCNA2,TROAP,UBE2C,CENPA,TPX2,SKA2,RACGAP1,LMNB1,PTTG1,HJURP,CCNB2,NUSAP1,NUF2,CCNF,GTSE1,PLK1,PRC1,KIF23,INCENP,NEK2,KIF2C,KIF20A,UBE2T,KIF18A,MXD3,AURKA,NCAPH,KIF18B,CDKN2D,CD36,KIF4A,HMMR,RGS3,CDC20,AURKB,NDC80,TTK,ESPL1,ANLN,E2F2,DIAPH3,KIFC1,MAD2L1,KIF15,SPAG5,NEIL3,S100A10,ID1,ATAD2,FBXO5,EZH2,UHRF1,DEPDC1B,LAMB3,SPC25,ASF1B,BUB1B,DLGAP5,BIRC5,CENPE,G0S2,PBK,CDCA8,PCNA,CDC25C,TACC3,CKS2,PLK4,TK1 |
| 7 | wgEncodeHaibTfbsSknshFoxm1sc502V0422111PkRep2.broadPeak.gz | 0.165501 | 7.51852 | T3 | FOXM1 | GPSM2,CCNA2,CDKN3,CENPA,CCNB1,CDK1,GTSE1,CCNF,RASGRP3,LMNB1,HJURP,CDKN2D,PTTG1,UBE2C,NUF2,KIF20A,TPX2,PLK1,KIF23,TROAP,PRC1,NEK2,CCNB2,KIF18B,RACGAP1,INCENP,EZH2,SKA2,KIF2C,MXD3,UBE2T,KIF18A,NCAPH,PDCD1LG2,ESPL1,KIF15,AURKA,HMMR,PBK,KIF4A,BUB1B,TTK,KIFC1,NUSAP1,BIRC5,CDC20,ARHGAP11A,NEIL3,DLGAP5,MMP2,SIGLEC1,ZWINT,IFI35 |
| 8 | wgEncodeHaibTfbsMcf7Foxm1sc502V0422111PkRep1.broadPeak.gz | 0.157616 | 7.10985 | T3 | FOXM1 | CDKN3,CDK1,GPSM2,CCNB1,UBE2C,CENPA,LMNB1,SKA2,KIF23,CCNF,TROAP,PTTG1,RACGAP1,NUF2,CCNA2,CDKN2D,PRC1,HJURP,KIF20A,NCAPH,ADORA3,BIRC5,CCNB2,GTSE1,TPX2,KIF18B,MXD3,RGS3,PLK1,KIF2C,KIF18A,UBE2T,FBXO5,INCENP,ATAD2,KIFC1,ANLN,NEK2,ESPL1,HMMR,EZH2,TACC3,AURKA,AURKB,TTK,PBK,SPAG5,NUSAP1,ID1,DEPDC1B,HMGB3,UHRF1,MYB,NEIL3,SMPDL3A,KIF4A,DLGAP5,CDC25C,NCAPG2,CD36,CDCA8,FANCI |
| 9 | wgEncodeHaibTfbsPanc1Sin3ak20V0416101PkRep2.broadPeak.gz | 0.144428 | 6.4264 | T4 | SIN3A | ANLN,ID1,KIF2C,FANCI,DTL,NEK2,NUF2,BLM,FBXO5,TPX2,IQGAP3,RAD51,TTK,CCNB2,CDC20,CDCA5,CDK1,EXO1,AURKA,HELLS,SHCBP1,ORC1,MCM4,RAD54L,BUB1B,CCNB1,CENPQ,KIF23,HMMR,CCNA2,CCNE2,BIRC5,PRC1,UBE2C,CLSPN,ATAD2,KIF18B,KIF20A,PLK4,AURKB,KIF18A,CENPW,NCAPH,NUSAP1,KIF4A,DSCC1,TK1,KIFC1,GGH,NET1,PCNA,TROAP,MAD2L1,ZWINT,CDCA8,PLK1,HJURP,CDC25C,DLGAP5,NCAPG,TCF19,CDC25A,CDC45,TRIP13,HSPA1B,RCC1,RAD54B,SPAG5,MELK,CDCA3,ERCC6L,LMNB1,GINS1,CDKN2D,ASF1B,GTSE1,MYBL2,CENPA,NEIL3,SLC25A19 |

| | | | | | | |
|---|---|---|---|---|---|---|
| 10 | wgEncodeHaibTfbsSknshSin3ak20V0416101PkRep1.broadPeak.gz | 0.141161 | 6.25714 | T4 | SIN3A | FAM72B,KIF2C,FAM72D,SKA2,NCAPG,CDC20,NUF2,KIF20A,SHCBP1,IQGAP3,MAD2L1,GGH,TK1,KIF18B,NEK2,NCAPG2,ESCO2,BIRC5,DLGAP5,BUB1B,CCNB1,HJURP,GTSE1,TROAP,CCNA2,NET1,CLSPN,MND1,CDCA8,RCC1,KIF15,FBXO5,CENPA,TACC3,CCNE2,RAD51,KIF4A,NEIL3,CDCA5,AURKB,EXO1,PLK1,NUSAP1,FANCI,MYBL2,PBK,ARHGAP11A,RAD54L,PCNA,BLM,CENPW,ANLN,UBE2C,CDCA3,KIF18A,TUBA1B,RAD54B,MCM5,DTL,CDC25A,CDK1,ESPL1,KIF23,LMNB1,RACGAP1,TCF19,GINS1,CKS2,MXD3,TTK,RASGRP3,NDC80,CENPQ,CDC45,SLC25A19,UHRF1,KIFC1,TYMS,ZNF367,TUBA1A,CDKN2D,SPC24,ZWINT,UBE2T,DIAPH3,MCM2,AURKA,NCAPH,CDC25C,PARP9,PLK4,ORC1,SPARC,HMMR,CENPE,ERCC6L,TUBB3,LIMA1,PRC1,DSCC1,STMN1,CCNB2,HELLS |
| 11 | wgEncodeSydhTfbsMcf10aesE2f4TamHvdPk.narrowPeak.gz | 0.118446 | 5.07993 | T1 | E2F4 | ZNF367,GINS2,TCF19,NCAPG2,CDT1,TACC3,ASF1B,MYBL2,TYMS,TROAP,HMGB3,FAM72D,BIRC5,ARHGAP11A,PLK1,FOXM1,TK1,CDCA5,CDCA8,CDKN3,CLSPN,CENPA,NUSAP1,CORO7,PRC1,UBE2C,MCM2,NCAPH,FADS2,IRF7,CDKN2D,ERCC6L,ANLN,CDC25A,MCM5,GTSE1,RAD51,E2F2,ESPL1,CDCA3,FAM72B,EZH2,SKA2,MCM4,FANCI,UHRF1,AURKB,NET1,KIF4A,PKMYT1,CCNF,RAD54L,CKS2,RCC1,CDC25C,GMNN,CDC20,AURKA,BLM,MELK,CDC7,DLGAP5,RACGAP1,CCNB1,SHCBP1,ATAD2,INCENP,NCAPG,KIF15,UBE2T,LMNB1,PCNA,MXD3,TUBA1B,SPC25,NEIL3,TRIP13,MND1,MCM3,PBK,FEN1,ZWINT,KIF2C,CENPW,KIFC1,DIAPH3,KIF23,KIF18B,WDR76,GPSM2,SPAG5,TPX2,PLK4,NEK2,FBXO5,DSCC1,MAD2L1,SREBF1,KIF20A,GINS1,IQGAP3,ID1,CCNB2,CDC45,DEPDC1B,ORC1,MID1IP1,CENPQ,GGH,NDC80 |
| 12 | wgEncodeHaibTfbsHepg2Mybl2sc81192V0422111PkRep1.broadPeak.gz | 0.109618 | 4.62246 | T5 | MYBL2 | ID1,SKA2,AURKA,EZH2,CKS2,PLK1,HMMR,SERPINA1,TUBA1B,TPX2,CENPA,CDKN2D,PRC1,CCNB1,LMNB1,NUSAP1,FBXO5,KIFC1,HJURP,TYMS,RACGAP1,BIRC5,UBE2C,PCNA,NCAPG,BUB1B,ANLN,KIF20A,INCENP,KIF15,NDC80,TUBB3,FAM72D,KIF18B,CDC20,CCNA2,CDK1,GTSE1,TK1,PBK,CENPE,AURKB,CCL20,ASF1B,ARHGAP11A,UBE2T,NCAPH,NEK2,CDCA8,ATAD2,PSRC1,FAM72B,NUF2,DEPDC1B,PLK4,DLGAP5,MCM4,MXD3,KIF2C,IL1RN,GMNN,CDCA3,UHRF1,MAD2L1,KIF23,HSPA1B,SPC25,RGS3,DTL,CDKN3,ARHGAP18,CDC25C,CCNB2,LYZ,HSPA6,TTK |
| 13 | GSM1208730_batch2_chrom1_LoVo_E2F7_PassedQC_peaks_hg19 | 0.109471 | 4.61484 | T6 | E2F7 | TYMS,MCM5,E2F2,WDR76,MCM3,ZNF367,RAD51,MCM4,PCNA,ATAD2,TK1,DTL,CDC25A,CDC45,MCM2,LMNB1,HELLS,RAD54L,GMNN,ASF1B,GINS2,MXD3,ORC1,EXO1,SKA2,FANCI,NUSAP1,CDC7,CCNE2,STMN1,BLM,PRC1,CDC25C,FBXO5,DSCC1 |

| | | | | | |
|---|---|---|---|---|---|
| 14 | wgEncodeHaibTfbsMcf7Foxm1sc502V0422111PkRep2.broadPeak.gz | 0.105781 | 4.42362 | T3 | FOXM1 | CDKN3,CDK1,GPSM2,UBE2C,SKA2,CCNB1,CENPA,KIF23,CCNA2,ADORA3,TPX2,LMNB1,CD36,TROAP,CCNF,PRC1,S100A10,NUF2,KIF20A,NEK2,RACGAP1,PTTG1,HJURP,MXD3,NCAPH,KIF18B,FBXO5,CDKN2D,ID1,PLK1,SMPDL3A,IRF8,UBE2T,INCENP,CDC20,GTSE1 |
| 15 | wgEncodeHaibTfbsSknshFoxm1sc502V0422111PkRep1.broadPeak.gz | 0.0855078 | 3.37297 | T3 | FOXM1 | RASGRP3,CDK1,LMNB1,GPSM2,CCNB1,CCNF,UBE2C,NEK2,GTSE1,TROAP,CDKN3,CENPA,CCNA2,PTTG1,CDKN2D,NCAPH,HJURP,CCNB2,MYLIP,TPX2,PDCD1LG2,SKA2 |
| 16 | wgEncodeSydhTfbsMcf7Hae2f1UcdPk.narrowPeak.gz | 0.0854902 | 3.37206 | T7 | E2F1 | UHRF1,CDK1,DTL,RAD54L,MCM3,PBK,PLK4,WDR76,ZWINT,UBE2T,NCAPG,CDC7,PCNA,MCM4,PRC1,TP73,ASF1B,CLSPN,TCF19,CDC25A,CDC45,KIF18B,MCM2,KIF15,CCNE2,NEK2,CDKN2D,NUSAP1,GMNN,NDC80,FEN1,FANCI,GINS1,FBXO5,ESCO2,ORC1,TYMS,EZH2,SKA2,SREBF1,ATAD2,SPAG5,PARP9,EXO1,KIFC1,BLM,SCD,GGH,CDC25C,LMNB1,INCENP,KIF2C,CDT1,E2F2,RCC1,NET1,ADORA3,TUBA1B,MCM5,SPC25,CENPQ,KIF4A,GTSE1,CORO7,GPSM2,KIF18A,RAD54B,SPC24,SLC25A19,DSCC1,RACGAP1,DIAPH3,NCAPH,HJURP,FPR3,ZNF367,MND1,HELLS,NCAPG2,BUB1B,MYB,PKMYT1,AURKB,NUF2,RGS3,HSPA6,ANLN,IQGAP3,MYBL2,NEIL3,SHROOM2,CDCA8,CHRNA5,AURKA |
| 17 | wgEncodeHaibTfbsMcf7Sin3ak20V0422111PkRep2.broadPeak.gz | 0.0814472 | 3.16254 | T4 | SIN3A | SKA2,ATAD2,CCNA2,CDC25C,AURKB,NEIL3,CCNE2,TUBA1B,TTK,ZNF367,FANCI,UBE2C,PLK4,FBXO5,PLK1,BLM,ESCO2,CDKN2D,RAD51,DEPDC1B,CDC20,NET1,MXD3,MCM4,MYB,TUBB3,KIF20A,CCNB1,CDK1,DIAPH3,KIF23,ADORA3,DSCC1,TYMS,NUF2,AURKA,PCNA,HSPA1B,BIRC5,ID1,CCNB2,TRIP13,CD36,ANLN,KIF4A,CDCA5,MYBL2,NEK2,RAD54B,KIF2C,BUB1B,EXO1,NCAPG2,NUSAP1,GINS1,ASF1B,TROAP,MAD2L1,LMNB1,RAD54L,SPARC,CKS2,KIF18A,KIF18B,PRC1,TPX2,IQGAP3,CDCA8,CENPW,GGH,PBK,PKMYT1,S100A10,CLSPN,HJURP,HSPA1A,DTL,RGS3,INCENP,STMN1,NCAPH,WDR76,UBE2T,CDC45,CDC25A,CENPA,MCM3,NCAPG,TDRD9,TUBA1A,GMNN,TACC3,TK1,DLGAP5,FEN1,SLC25A19,E2F2,CENPQ,SPC24,HMMR,SHCBP1,FXYD2,CENPE,NDC80,MYLIP,PTTG1,SREBF1,ZWINT,TCF19,PSRC1,IRF8,MCM2,RACGAP1,KIFC1,SCD,MND1,RCC1,MID1IP1,ORC1,ESPL1,CDC7,CDKN3 |
| 18 | wgEncodeHaibTfbsHepg2Sin3ak20Pcr1xPkRep2.broadPeak.gz | 0.0802232 | 3.0991 | T4 | SIN3A | HSPA1B,HSPA1A,FAM72D,ARHGAP11A,FAM72B,FBXO5,SHCBP1,CDC25A,NEK2,CCNA2,MAD2L1,CCNE2,NCAPG,PCNA,TRIP13,CCNB1,ANLN,NUF2,TACC3,CLSPN,RAD51,PLK1,KIF20A,SKA2,FANCI,CDC20,MYBL2,AURKA,HJURP,FOXM1,KIF18A,RAD54L,CCNB2,KIF15,CKS2,CDKN2D,ATAD2,NCAPG2,KIFC1,NEIL3,PLK4,TUBB3,BLM,PBK,BUB1B,NCAPH,ESCO2,KIF4A,SREBF1,MXD3,CDCA5,ASF1B,KIF18B,NUSAP1,MND1,ID1,IQGAP3,DLGAP5,GINS1,GMNN,CENPQ,AURKB,TROAP,BIRC5,TTK,CDCA8,PRC1,TCF19,DTL,NET1,DIAPH3,KIF2C,FEN1,CDK1,CDC25C,KIF23,TPX2 |

Supplementary Table 8. Transcription factor predicted by Enrichr

**Transcription factors predicted by DEGs upregulated in HLF cells**

| Term | Overlap | P-value | Adjusted P-value | Old P-value | Old Adjusted P-value | Odds Ratio | Combined Score | Genes |
|---|---|---|---|---|---|---|---|---|
| AR CHEA | 5/1095 | 0.0006 | 0.0206 | 0 | 0 | 9.63 | 70.77 | ABCA10;RELN;ABLIM3;PSAT1;PAPPA2 |
| SUZ12 CHEA | 4/1684 | 0.0253 | 0.4042 | 0 | 0 | 4.36 | 16.03 | RELN;SLIT2;GPR156;SLC7A2 |
| FOS ENCODE | 2/637 | 0.0716 | 0.5545 | 0 | 0 | 5.08 | 13.39 | FKBP14;PSAT1 |
| SUZ12 ENCODE | 1/105 | 0.0711 | 0.5545 | 0 | 0 | 14.71 | 38.88 | SLIT2 |
| ZC3H11A ENCODE | 1/129 | 0.0866 | 0.5545 | 0 | 0 | 11.93 | 29.19 | ABLIM3 |
| BCL3 ENCODE | 1/273 | 0.1751 | 0.5881 | 0 | 0 | 5.58 | 9.71 | SYCE2 |
| PBX3 ENCODE | 2/1269 | 0.2217 | 0.5881 | 0 | 0 | 2.46 | 3.71 | SYCE2;FKBP14 |
| SALL4 CHEA | 1/355 | 0.2218 | 0.5881 | 0 | 0 | 4.27 | 6.42 | ASNS |
| SMC3 ENCODE | 2/1181 | 0.1987 | 0.5881 | 0 | 0 | 2.66 | 4.30 | ABLIM3;GPR156 |
| STAT3 CHEA | 1/177 | 0.1171 | 0.5881 | 0 | 0 | 8.66 | 18.57 | ASNS |

**Transcription factors predicted by DEGs downregulated in HLF cells**

| Term | Overlap | P-value | Adjusted P-value | Old P-value | Old Adjusted P-value | Odds Ratio | Combined Score | Genes |
|---|---|---|---|---|---|---|---|---|
| IRF1 ENCODE | 8/263 | 0.0000 | 0.0000 | 0 | 0 | 15.45 | 238.89 | ANKRD37;PLSCR1;STAT1;MX1;CMPK2;ISG15;IFI35;PARP9 |
| IRF8 CHEA | 5/121 | 0.0000 | 0.0004 | 0 | 0 | 19.88 | 228.19 | IFIH1;ISG20;CD74;EPSTI1;IFI35 |
| STAT3 ENCODE | 7/725 | 0.0016 | 0.0430 | 0 | 0 | 4.57 | 29.41 | HERC5;ISG20;PLSCR1;RRM2;SP110;STAT1;TMEM140 |
| VDR CHEA | 2/76 | 0.0144 | 0.2872 | 0 | 0 | 11.68 | 49.56 | SP110;EPSTI1 |
| NELFE ENCODE | 3/234 | 0.0186 | 0.2973 | 0 | 0 | 5.69 | 22.68 | RRM2;SNHG12;FASN |
| SALL4 CHEA | 3/355 | 0.0534 | 0.7117 | 0 | 0 | 3.71 | 10.88 | FASN;E2F1;PRSS23 |
| E2F6 ENCODE | 12/3245 | 0.0781 | 0.8140 | 0 | 0 | 1.72 | 4.40 | SREBF1;ANKRD37;PLSCR1;RRM2;SNHG12;STAT1;FASN;E2F1;TRIM14;CMPK2;LAP3;PARP9 |
| POU5F1 CHEA | 2/261 | 0.1297 | 0.8140 | 0 | 0 | 3.31 | 6.75 | PLSCR1;FASN |
| PPARG CHEA | 3/535 | 0.1368 | 0.8140 | 0 | 0 | 2.43 | 4.84 | LPCAT3;TMEM140;BCL2L1 |
| RELA ENCODE | 3/484 | 0.1099 | 0.8140 | 0 | 0 | 2.70 | 5.96 | IFIH1;CD74;STAT1 |

**Transcription factors predicted by DEGs upregulated in macrophages**

| Term | Overlap | P-value | Adjusted P-value | Old P-value | Old Adjusted P-value | Odds Ratio | Combined Score | Genes |
|---|---|---|---|---|---|---|---|---|

| | | | | | | | | |
|---|---|---|---|---|---|---|---|---|
| CEBPB ENCODE | 7/144 | 0.0033 | 0.1882 | 0 | 0 | 3.84 | 21.89 | XPOT;TNNC1;PYCR1;CHAC1;SLC7A11;TBL1X;GTPBP2 |
| ZBTB7A ENCODE | 44/2184 | 0.0038 | 0.1882 | 0 | 0 | 1.61 | 8.98 | NCKAP1;ZFAND3;PHF1;TESK1;PRSS27;SLC1A4;AKAP5;RFNG;SENP6;CBS;FTH1;CYP4V2;MKNK2;ZNF425;D2HGDH;TBL1X;DCAF10;PRKACB;PDE8A;SH3GLB1;HEMK1;UNC13A;BRPF3;PYCR1;ELOVL6;SLC30A1;GTPBP2;NBPF1;KIF27;P2RX4;XPOT;MYO15B;RILPL1;CCPG1;MAP1B;UVSSA;NOXA1;CCSER2;TCEA1;FLYWCH1;SQSTM1;KCTD15;ATF4;PPIL6 |
| E2F6 ENCODE | 50/3245 | 0.1577 | 1.0000 | 0 | 0 | 1.19 | 2.19 | DHRS12;SETD9;PHF1;FBXO27;TESK1;ALDH1L2;CLGN;PTER;NAMPT;PDE8A;CCDC112;FBXW4;PARP6;SLC38A1;GPT2;VPS13C;WDR31;GAB2;GTPBP2;GPCPD1;NBPF1;KIF27;MTHFD2;RILPL1;ATF5;SQSTM1;FTCDNL1;ATF4;EPAS1;NPL;SLC1A5;CBS;STC2;MAP7;ZNF425;DCAF10;PCDH1;PCK2;DTNB;PYCR1;NR1D2;SNHG7;TDRD3;PER3;MYO15B;PSAT1;TCEA1;ULBP1;NFE2L1;PPIL6 |
| ESR1 CHEA | 5/154 | 0.0566 | 1.0000 | 0 | 0 | 2.50 | 7.18 | CYP27A1;C1QTNF6;CAPN5;PDK4;GAB2 |
| NFE2L2 CHEA | 18/1022 | 0.1445 | 1.0000 | 0 | 0 | 1.34 | 2.60 | SLC48A1;GRIA2;UNC5B;GPT2;SLC41A2;ARAP2;NR1D2;TUBE1;SLC30A1;SLC1A4;SLC7A11;SIPA1L2;GPCPD1;FAM210B;FTH1;STC2;GRB10;SQSTM1 |
| RUNX1 CHEA | 23/1294 | 0.1018 | 1.0000 | 0 | 0 | 1.36 | 3.12 | CCDC112;DENND2D;SH3GLB1;SLC38A1;UNC13A;DTNB;ACSL1;TSEN15;STRBP;ICA1;ELOVL6;FGR;LGALS3;MTHFD2;CCPG1;RPS6KA2;FTH1;TNIP3;MKNK2;GRB10;TBL1X;NFE2L1;NSMAF |
| SALL4 CHEA | 7/355 | 0.2005 | 1.0000 | 0 | 0 | 1.49 | 2.40 | UNC5B;RUFY3;SESN2;ASNS;ELOVL6;PDE8A;NSMAF |
| TCF3 CHEA | 17/1006 | 0.1943 | 1.0000 | 0 | 0 | 1.28 | 2.10 | UNC5B;FBXO27;SLC41A2;ASNS;ELOVL6;SLC1A5;TDRD3;SEL1L3;KLF8;SPIRE1;PLIN2;TBL1X;RFFL;PRKACB;PDE8A;ZBED3;NSMAF |
| TCF3 ENCODE | 15/840 | 0.1591 | 1.0000 | 0 | 0 | 1.36 | 2.50 | SH3GLB1;FBXW4;SLC48A1;ACSL1;NR1D2;TDRD3;MAN2A2;NAMPT;CHAC1;TBL1X;PDE8A;SQSTM1;NFE2L1;ATF4;RNF187 |
| TP53 CHEA | 7/319 | 0.1382 | 1.0000 | 0 | 0 | 1.67 | 3.30 | FBXW4;SMARCD3;UNC5B;VPS13C;SESN2;SLC30A1;SPATA18 |

**Transcription factors predicted by DEGs downregulated in macrophages**

| Term | Overlap | P-value | Adjusted P-value | Old P-value | Old Adjusted P-value | Odds Ratio | Combined Score | Genes |
|---|---|---|---|---|---|---|---|---|
| E2F4 ENCODE | 103/710 | 0.0000 | 0.0000 | 0 | 0 | 17.33 | 2972.50 | ERCC6L;DSCC1;CCNF;GMNN;HJURP;BUB1B;CDC20;NUSAP1;NEK2;FBXO5;GTSE1;ESCO2;CDC25C;WDR76;CDC25A;NEIL3;DEPDC1B;MELK;CCNE2;KIF20A;ASF1B;BLM;CDCA3;CDCA5;TROAP;CDCA8;NCAPG;PKMYT1;IQGAP3;NCAPH;SKA2;RACGAP1;ORC1;CLSPN;PLK4;FANCI;CDT1;PLK1;CDC7;NDC80;NET1;TPX2;KIF18A;DIAPH3;KIF18B;UBE2T;KIF4A;CDK1;MXD3;EZH2;ARHGAP11A;FEN1;NCAPG2;FOXM1;KIF15;LMNB1;TUBA1B;EXO1;NUF2;PBK;MYBL2;TK1;ZNF367;DLGAP5;HELLS;RMI2;KIF23;CCNA2;ESPL1;INCENP;SLC25A19;MCM3;MCM4;BIRC5;KIF2C;MCM5;DTL;MCM2;PCNA;UHRF1;TTK;TYMS;AURKB;RAD54B;AURKA;CDC45;RAD54L;E2F2;GINS1;GINS2;CENPW;SPAG5;ATAD2;MND1;SHCBP1;RAD51;PRC1;TRIP13;CENPQ;SPC24;CDKN3;MAD2L1;SPC25 |

| | | | | | | | | |
|---|---|---|---|---|---|---|---|---|
| FOXM1 ENCODE | 30/95 | 0.0000 | 0.0000 | 0 | 0 | 34.87 | 2530.14 | GPSM2;TROAP;CCNF;HJURP;CENPA;NCAPH;LMNB1;SKA2;CDC20;CCNB2;CCNB1;PTTG1;RACGAP1;NUF2;NEK2;GTSE1;CDKN2D;UBE2C;PLK1;KIF23;CCNA2;TPX2;KIF18B;INCENP;PRC1;UBE2T;CDK1;KIF20A;MXD3;CDKN3 |
| SIN3A ENCODE | 60/1131 | 0.0000 | 0.0000 | 0 | 0 | 4.54 | 187.95 | DSCC1;HJURP;BUB1B;KIF15;LMNB1;CDC20;TUBA1B;EXO1;NUF2;PBK;NEK2;FBXO5;GTSE1;ZNF367;DLGAP5;HELLS;KIF23;ESCO2;CDC25C;CDC25A;CCNA2;CCNE2;CKS2;MCM3;MCM4;BIRC5;KIF2C;KIF20A;DTL;BLM;CDCA5;CDCA8;NCAPG;HMMR;IQGAP3;NCAPH;AURKB;SKA2;AURKA;CCNB2;CCNB1;ORC1;CLSPN;PLK4;CENPW;PLK1;ATAD2;SHCBP1;TPX2;ANLN;CENPE;KIF18A;KIF18B;RAD51;ID1;TACC3;CDK1;TRIP13;CENPQ;MAD2L1 |
| NFYA ENCODE | 72/2250 | 0.0000 | 0.0000 | 0 | 0 | 2.66 | 62.84 | CCNF;GMNN;HJURP;HMGB3;CDC20;NUSAP1;NEK2;FBXO5;TK1;GTSE1;ZNF367;DLGAP5;RMI2;KIF23;ESCO2;CDC25C;CDC25A;CCNA2;DEPDC1B;CCNE2;ESPL1;INCENP;SLC25A19;CKS2;MCM3;MCM4;MCM5;KIF20A;SNRNP25;DTL;FAM72B;ASF1B;BLM;PCNA;CDCA3;UHRF1;TROAP;CDCA8;TTK;HMMR;CENPA;AURKB;SKA2;AURKA;CCNB2;CCNB1;CDC45;RAD54L;CLSPN;PLK4;CDKN2D;GINS1;ATAD2;SHCBP1;ZWINT;NET1;TPX2;KIF18A;DIAPH3;KIF18B;SCD;PRC1;UBE2T;ID1;CDK1;CORO7;CENPQ;SPC24;MXD3;EZH2;CDKN3;SPC25 |
| NFYB ENCODE | 99/3715 | 0.0000 | 0.0000 | 0 | 0 | 2.31 | 51.96 | ERCC6L;CCNF;GMNN;HJURP;CDC20;TUBB3;STMN1;NUSAP1;NEK2;FBXO5;GTSE1;MYLIP;ESCO2;CDC25C;CDC25A;DEPDC1B;CCNE2;KIF20A;SNRNP25;ASF1B;BLM;CDCA3;TROAP;CDCA8;ARHGAP18;HMMR;SKA2;CCNB2;CCNB1;RACGAP1;CLSPN;PLK4;UBE2C;PLK1;CDC7;ZWINT;BST2;NET1;TPX2;KIF18A;DIAPH3;KIF18B;UBE2T;KIF4A;ID1;CDK1;CORO7;MXD3;EZH2;GPSM2;NCAPG2;HMGB3;LMNB1;TK1;ZNF367;DLGAP5;RMI2;SHROOM2;KIF23;CCNA2;DEPDC7;PSRC1;ESPL1;INCENP;SLC25A19;CKS2;MCM3;MCM4;MCM5;FAM72D;DTL;FAM72B;MCM2;PCNA;UHRF1;IFI6;TTK;CENPA;AURKB;RAD54B;AURKA;CDC45;RAD54L;CDKN2D;GINS1;GINS2;SPAG5;MID1IP1;ATAD2;SHCBP1;CENPE;SCD;PRC1;APOC1;TACC3;CENPQ;SPC24;CDKN3;SPC25 |
| E2F1 CHEA | 34/859 | 0.0000 | 0.0000 | 0 | 0 | 3.04 | 48.55 | ARHGAP11A;PCNA;CDCA5;GMNN;BUB1B;PKMYT1;AURKB;SKA2;LMNB1;AURKA;CCNB2;TUBA1B;EXO1;TK1;CLSPN;ZNF367;DLGAP5;S100A10;FANCI;CDT1;GINS2;RMI2;UBE2C;ATAD2;MND1;SHCBP1;CDC25A;NDC80;KIF18A;ESPL1;SLC25A19;SPC24;EZH2;MCM2 |
| E2F6 ENCODE | 66/3245 | 0.0022 | 0.0316 | 0 | 0 | 1.53 | 9.36 | FEN1;ERCC6L;DSCC1;CCNF;HILPDA;GMNN;KIF15;LMNB1;TUBA1B;STMN1;MYB;NUF2;PBK;MYBL2;FBXO5;TK1;ZNF367;SREBF1;RMI2;KIF23;ESCO2;PARP9;WDR76;CDC25A;DEPDC1B;DEPDC7;CCNE2;SLC25A19;IRF7;MCM3;MCM4;BIRC5;SNRNP25;DTL;ASF1B;MCM2;BLM;CDCA3;CHRNA5;UHRF1;NCAPG;TYMS;CENPA;NCAPH;AURKB;SKA2;RAD54B;CDC45;RAD54L;E2F2;CLSPN;CDKN2D;GINS1;FANCI;CDT1;GINS2;ATAD2;CDC7;MND1;ANLN;RAD51;FABP5;CDK1;TRIP13;MXD3;EZH2 |

| | | | | | | | | |
|---|---|---|---|---|---|---|---|---|
| KLF4 CHEA | 26/987 | 0.0025 | 0.0316 | 0 | 0 | 1.92 | 11.51 | IFITM1;PCNA;UHRF1;DSCC1;SKA2;TUBA1B;TUBB3;STMN1;MYB;NUF2;SPP1;NEK2;MYBL2;ZNF367;TCF19;ANLN;DEPDC7;FABP5;APOC1;BIRC5;ACKR3;TRIP13;NBL1;CENPQ;DTL;IRF9 |
| IRF3 ENCODE | 19/663 | 0.0039 | 0.0435 | 0 | 0 | 2.08 | 11.53 | BLM;CDCA3;UBE2C;CENPA;SKA2;AURKA;TPX2;INCENP;SLC25A19;MCM3;RAD54L;MCM5;FBXO5;FAM72D;DLGAP5;SPC24;ASF1B;CDKN3;SPC25 |
| SPI1 CHEA | 26/1056 | 0.0060 | 0.0611 | 0 | 0 | 1.79 | 9.12 | TRAF3IP3;CDCA5;CELF2;PIK3CD;RAD54B;RGS1;ADORA3;TAGAP;CCL4;STK38L;CD14;FANCI;SP110;IPCEF1;KIF18B;RAD51;INCENP;KIFC1;SLC25A19;CKS2;TACC3;BIRC5;S100A4;DTL;MAD2L1;TRIM34 |

Red indicates adjusted P-value < 0.05

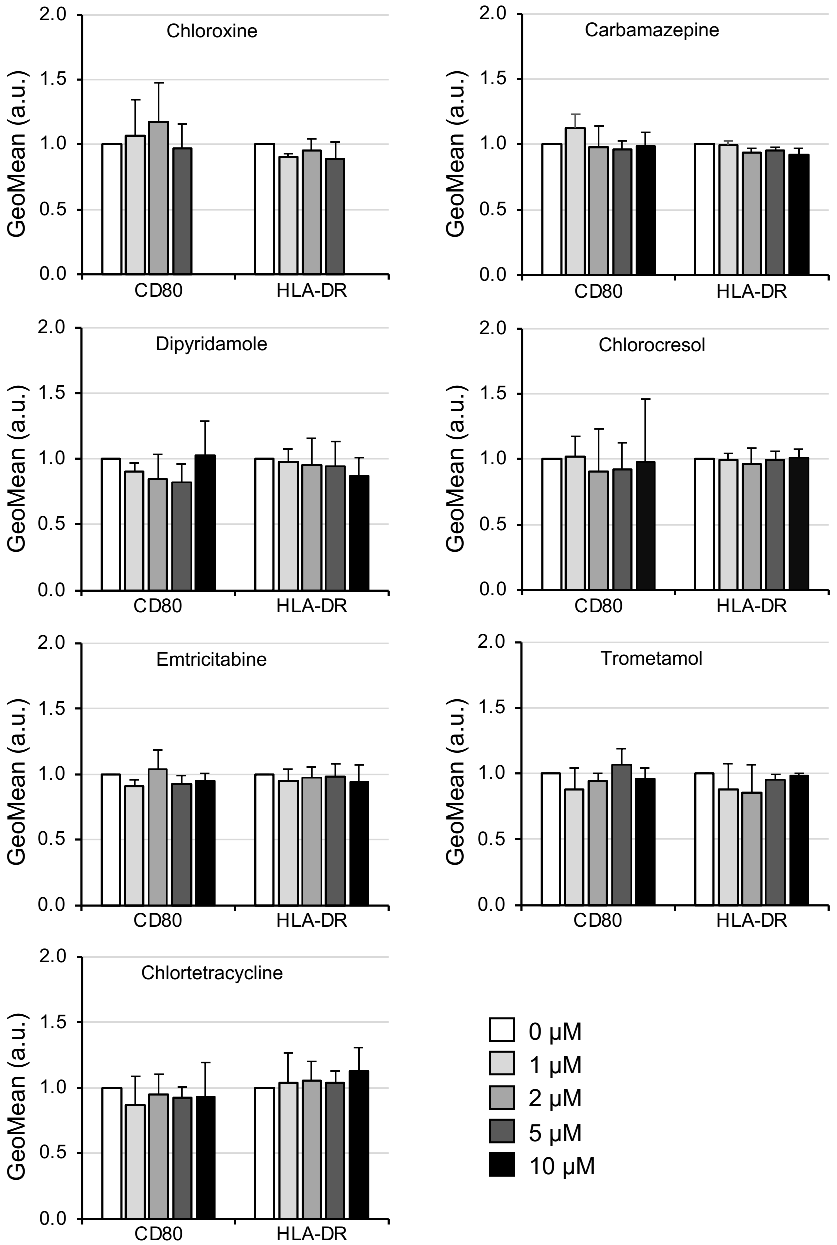


**Figure S1. Screening of TAM activators from an FDA-approved drug library.** CD80 and HLA-DR expressions on THP-1-derived macrophages in heterospheroids with HLF cells. The heterospheroids were incubated with the indicated concentrations of each drug for 3 days. No significant difference was observed vs. 0 μM, Dunnett's test ($n$ = 3).

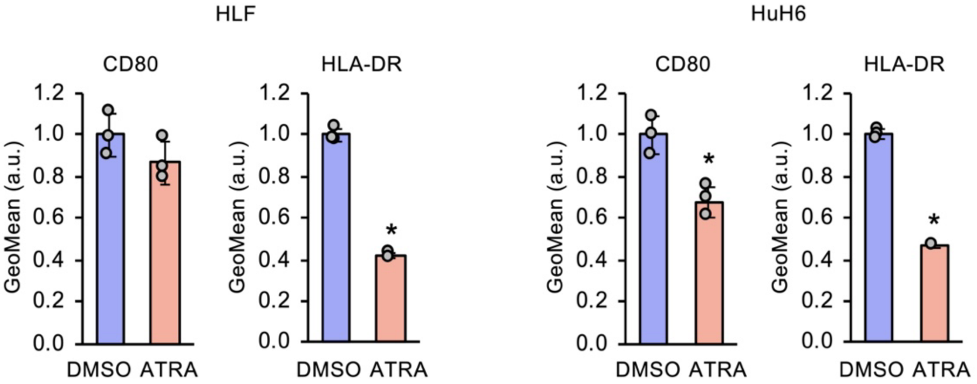


**Figure S2. Downregulation of M1 markers by all-trans-retinoic acid (ATRA).** CD80 and HLA-DR expressions on THP-1-derived macrophages in heterospheroids with HLF or HuH6 cells. The heterospheroids were incubated with 10 µM of ATRA for 3 days. *, $P < 0.05$ vs. DMSO, Student's *t*-test ($n = 3$).

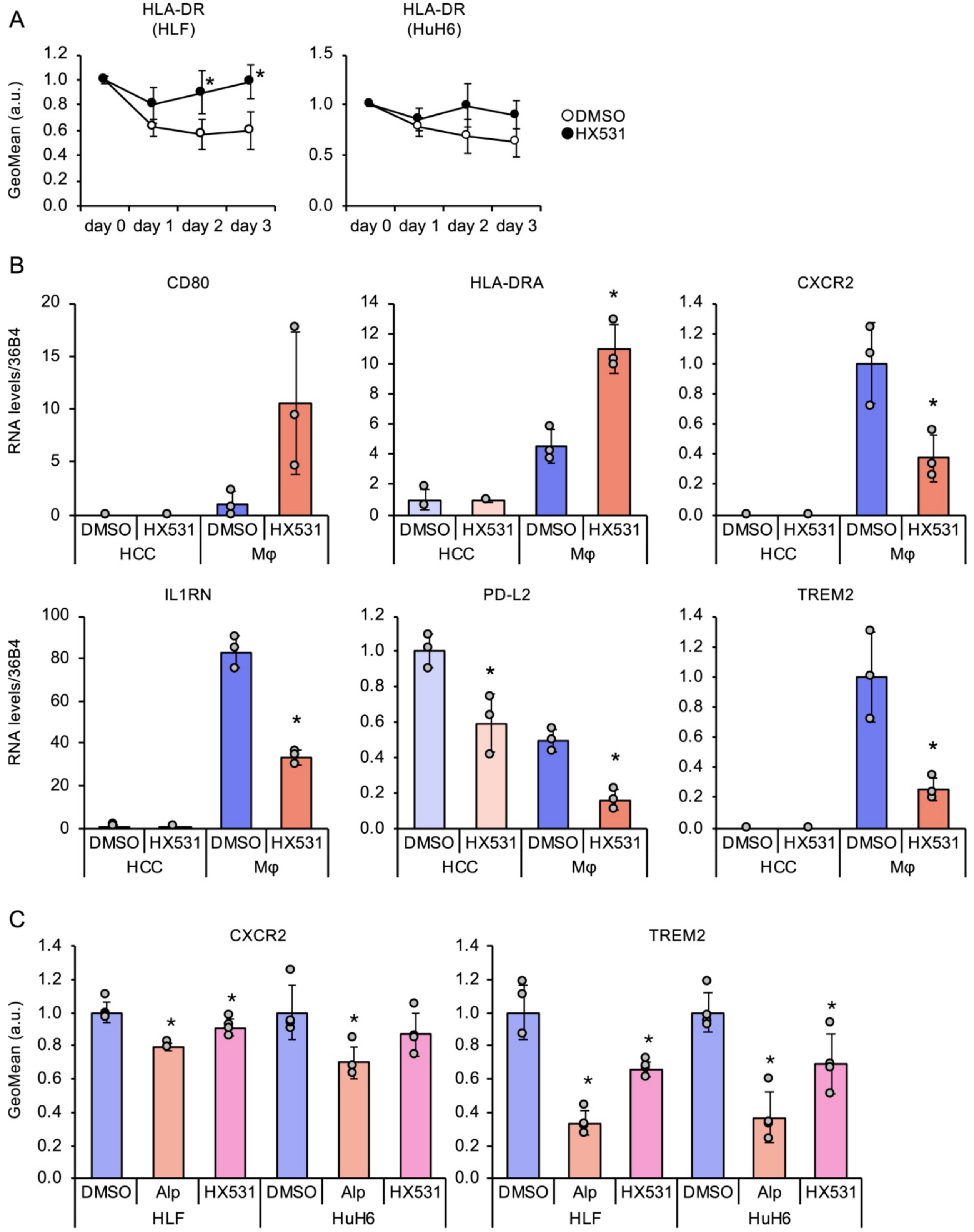


**Figure S3. Transcriptome analysis of heterospheroids treated with HX531.** (A) HLA-DR expressions on THP-1-derived macrophages in heterospheroids with HLF or HuH6 cells. The heterospheroids were incubated with DMSO or 2 μM of ATRA for the indicated days. *, $P < 0.05$ vs. DMSO, Student's *t*-test ($n = 3$). (B) Validation of RNA-seq analysis. MRNA were recovered from THP-1-derived macrophages and HLF cells in heterospheroids treated with DMSO or 1 μM HX531 for 2 days, and subjected to RT-qPCR. *, $P < 0.05$ vs. DMSO, Student's *t*-test ($n = 3$). (C) Cell surface CXCR2 and TREM2 expressions on THP-1-derived macrophages in heterospheroids with HLF or HuH6 cells. The heterospheroids were incubated with DMSO, 2 μM alprostadil (Alp), or 1 μM HX531 for 3 days. *, $P < 0.05$ vs. DMSO, Dunnett's test ($n = 4$).

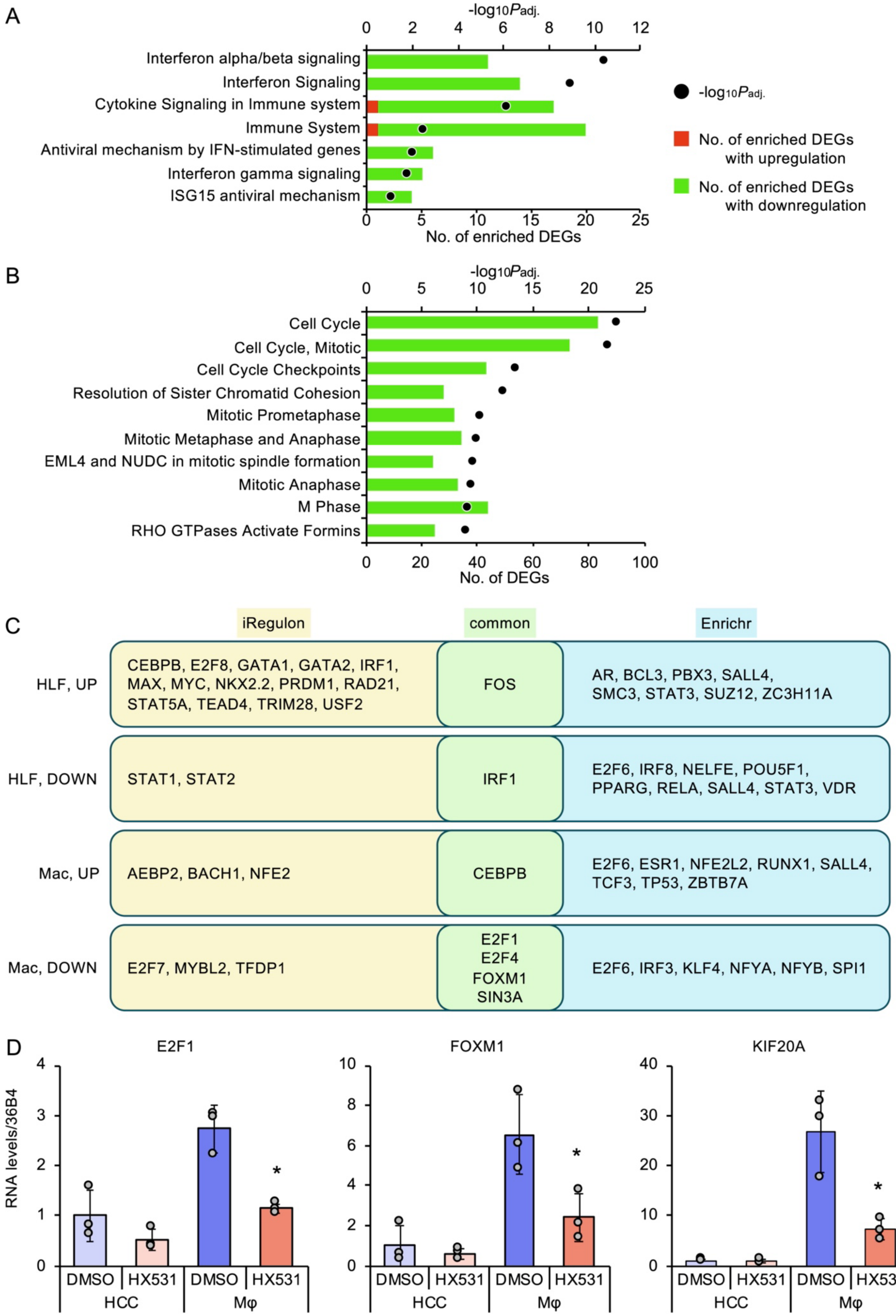


**Figure S4. Pathway analysis of heterospheroids treated with HX531.** (A,B) Reactome analyses of DEGs in HLF cells (A) and macrophages (B). (C) iRegulon and Enrichr analyses of DEGs in HLF cells and macrophages. (D) Validation of iRegulon and Enrichr analyses. MRNA were recovered from THP-1-derived macrophages and HLF cells in heterospheroids treated with DMSO or 1 μM HX531 for 2 days, and subjected to RT-qPCR. *, $P < 0.05$ vs.

DMSO, Student's *t*-test ($n = 3$).

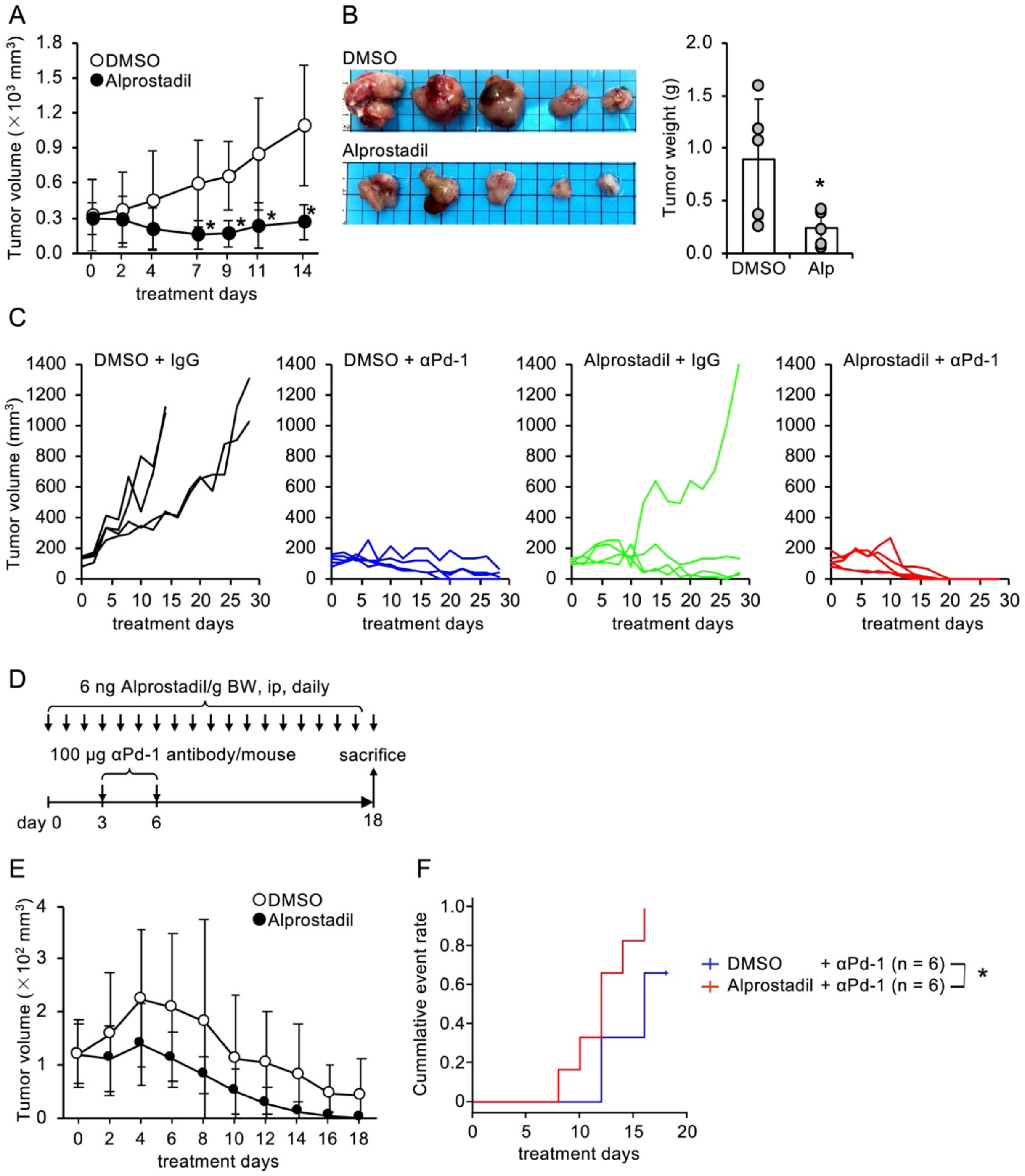


**Figure S5. In vivo efficacy of alprostadil.** (A) Tumor volume changes in mice treated with DMSO or alprostadil, as shown in Figure 7A. (B) Representative images and weights of tumor tissues after alprostadil (Alp) treatment. *, $P < 0.05$ vs. DMSO, Student's *t*-test ($n = 5$). (C) Tumor volume changes in each mouse treated with the combination of alprostadil and anti-Pd-1 antibody, as shown in Figure 7D. (D) Treatment schedule of combination of alprostadil and a reduced total dose of anti-Pd-1 antibody. (E,F) Tumor volume changes (E) and cumulative tumor free rate (F) of mice treated with the combination of alprostadil and anti-Pd-1 antibody, as shown in Figure S5D. *, $P < 0.05$, log-rank test.

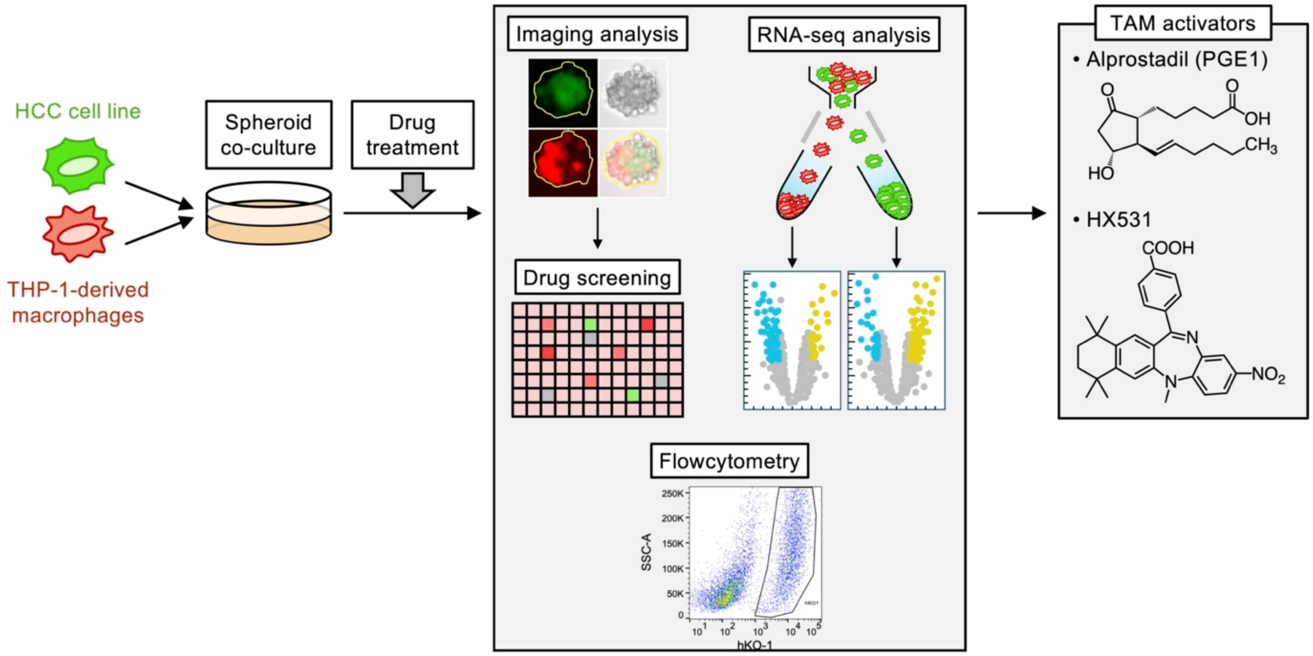


**Figure S6. Heterospheroids offer a new platform for oncoimmunological research and development.**
